# Understanding the biosynthesis of human IgMs through a combinatorial expression of mutant subunits that affect different assembly steps

**DOI:** 10.1101/2023.09.01.555973

**Authors:** Haruki Hasegawa, Songyu Wang, Eddie Kast, Hui-Ting Chou, Mehma Kaur, Tanakorn Janlaor, Mina Mostafavi, Yi-Ling Wang, Peng Li

## Abstract

Polymeric IgMs are secreted from plasma cells abundantly despite their structural complexity and intricate multimerization steps. To gain new insights into IgM’s assembly mechanics that underwrite the high-level secretion, we characterized the biosynthetic process of a natural human IgM, SAM-6, using a recombinant HEK293 cell system. By creating a series of mutant subunits that differentially disrupt specific sets of inter-chain disulfide bonds, we assessed their effects on various aspects of IgM biosynthesis in 48 different mutant subunit combinations. The analysis included the visualization of intracellular biosynthetic events such as steady-state subcellular subunit distribution, secretory trafficking bottlenecks, and the ER-associated Russell body formation by fluorescent microscopy. We also characterized various extracellular events including secreted IgM product quality, secretion output, and the release of various assembly intermediates using biochemical and biophysical assays. In this combinatorial mutagenesis approach, we unexpectedly found that the loss of multiple inter-chain disulfide bonds, including the one between μHC and λLC subunits, was tolerated in polymeric IgM formation and secretion. This finding revealed the vital role of underlying non-covalent protein-protein association not only during the orchestration of initial subunit interactions but also in maintaining the polymeric IgM product integrity during ER quality control steps, secretory pathway trafficking, and secretion. We suggest that the IgM assembly process is inherently robust and has a stopgap that permits the secretion of polymeric IgM even when not all the prescribed inter-chain disulfide bonds are formed. This study holistically presents the requirements and exemptions in polymeric IgM biosynthesis by encompassing the characterization of intracellular and extracellular events and the roles of covalent and non-covalent interactions. These findings can guide antibody engineering strategy when designing IgM-based multivalent modalities.

## 1. Introduction

Immunoglobulin M (IgM) is the most ancient antibody class conserved across all vertebrate species [1, 2]. IgMs are also unique among immunoglobulins in that they are produced as multimers. Depending on the availability of joining-chain (J-chain, JC) subunit during an assembly reaction in the endoplasmic reticulum (ER), μ heavy chain (HC) and λ or κ light chain (LC) oligomerize into pentamers or hexamers. Therefore, a single IgM protein can possess ten or twelve identical antigen binding sites per molecule [3]. Binding avidity gained from such polymeric formats is crucial during the primary immune response, where the multivalency is expected to compensate for the low affinity nature of the initial antibody repertoire elicited after a new antigen challenge. In addition to such ‘adaptive’ or ‘immune’ IgMs, there is also a class of IgMs called ‘innate’ or ‘natural’ IgMs that are produced in the absence of exogenous antigen exposure [4]. These natural IgMs play critical functions in immune surveillance by virtue of their low affinity target binding and poly-reactivity [4, 5]. The values of the IgM class for human therapeutics and diagnostic tools were recently re-recognized during the SARS-Cov-2 pandemic because the high mutation rate of surface glycoprotein and the concomitant generation of escape mutants was a significant concern for the antibody-based neutralization strategy using IgGs [6–8].

Besides the multivalency, additional attributes further distinguish IgMs from IgGs. For example, IgMs have a relatively short serum half-life of ∼4‒6 days in humans [5, 9] compared to ∼10‒21 days for IgGs [10]. Similarly, IgMs can activate complement pathways much more effectively than the IgG class and are perhaps better suited for immunotherapeutic strategies via complement-dependent cytotoxicity [11, 12]. While hexameric IgMs are reportedly 20-fold more potent than pentameric IgMs in activating the complement pathway to induce cytolysis [13, 14], only the pentameric IgM can cross the epithelial barrier by transcytosis through its JC component that interacts with the polymeric immunoglobulin receptor (pIgR) [15]. During the epithelial transport, pIgR ectodomain is released by proteolysis but remains associated with the pentameric IgM to give rise to a “secretory IgM,” which is released from the lumenal side of the epithelium as external secretions to serve as the first line of defense against pathogens that often favor respiratory and gastrointestinal mucosal surface as the portal of entry [16]. Randall et al. (1992) [17] showed that depending on how B cells were stimulated (i.e., LPS or IL-5), the activated cells produced hexameric and pentameric IgMs in different proportions. Altering the relative abundance of hexamers or pentamers under different inflammatory conditions indicates that it is an essential adaptive mechanism, with a trade-off between the two key functions—high cell lytic activity and epithelial transport. Because of such functional and physicochemical property differences, some investigators categorize pentameric IgM and hexameric IgM as two distinct subclasses [12].

IgM is also regarded as one of the most complex secretory proteins produced by the cell because of its megadalton size, multimeric nature, and intricate assembly cascade [18]. The complexity and the associated biosynthetic burden of a hexameric IgM formation can be appreciated by the need for a coordinated assembly of 12 μHCs and 12 LCs into one large hexameric IgM protein complex (∼1,100 kDa) while being post-translationally modified by 60 N-linked glycans, 84 intra-domain disulfides, and 30 inter-chain disulfide bridges concurrently. In the case of a pentameric IgM (∼950 kDa), one of the protomers is replaced by a JC that itself has 1 N-linked glycan site and 8 cysteine (Cys) residues involved in 3 intra-molecule and 2 inter-chain disulfide bridge formation. Despite the complex assembly requirements, plasma cells in the bone marrow manage to sustain a production of multimeric IgMs in the order of 10^8^ molecules/hour/cell (or ∼25,000 molecules/second/cell) [19, 20].

Partly because of the early studies reporting technical challenges on recombinant IgM expression [21–23], the IgM class has not been explored as a modality of human therapeutics as much as the IgG counterpart. Consequently, there is currently no approved marketed drug that uses IgM or IgM-like molecular design (https://www.antibodysociety.org/resources/approved-antibodies/). Nonetheless, constant efforts have been made to use IgMs and IgM-like molecules for therapeutic or diagnostic use [24]. Such attempts have ranged from (a) developing natural human IgMs isolated from patients to treat diseases [25–27], (b) augmenting the potency of IgGs through avidity by multimerizing IgGs into an IgM-like format by a tailpiece grafting [11, 28, 29], (c) fusing a scFv [30] or the ectodomain of a receptor [31] to the μHC’s constant region to enhance their target binding through avidity, and (d) using the pentameric IgM structure as a template to engineer novel multivalent bi-specific antibodies [32].

Prompted by the recent pandemic, there has been a renewed interest in using recombinant IgMs and IgM-like molecules as treatment options and diagnosis tools for human diseases [1, 24]. To extend our understanding of IgMs’ assembly processes and secretion requirements, we characterize the biosynthetic process of a natural human IgM called SAM-6 as our model cargo. SAM-6 was originally isolated from a stomach cancer patient as an antibody clone expected to possess anti-tumor activities [33]. Its sequence revealed that both HC and LC were encoded by unmutated germline genes [27, 34]. As expected from the polyreactive nature of natural IgM [35], SAM-6 was shown to bind to a tumor-specific 140 kDa membrane-anchored protein [34], oxidized LDL [33, 36], and the O-linked carbohydrate moiety of a novel cancer-specific variant of the ER-resident GRP78 that mislocalized on the cell surface [37, 38]. The proposed mechanism of action for SAM-6’s cancer-killing activity was also intriguing in that SAM-6 induces the over-accumulation of intracellular lipids in cancer cells and kills them by lipotoxicity [33, 34, 39]. Although subsequent attempts to test SAM-6 in human clinical trials did not show objective therapeutic responses as a single agent [40, 41], the IgM SAM-6 fulfilled a role as the benchmark molecule when exploring the secretory capacity of a novel human cell line [42] or a plant expression system [43]. SAM-6 also helped reveal site-dependent N-linked glycan type variations within an IgM molecule [44].

To gain a better mechanistic understanding and control over the production of IgMs, we investigate SAM-6 biosynthesis in a fully recombinant setting where all the input sequences are designed and selectively expressed in a heterologous HEK293 cell system. We are especially interested in determining whether all the inter-chain associations via disulfide bonds are essential in the formation and secretion of polymeric IgMs. To examine the importance of individual inter-chain disulfide bonds in polymeric IgM formation and secretion, μHC’s Cys-137, Cys-337, Cys-414, or Cys-575 residue is mutated singly or in combination with or without an additional CH1 domain deletion. Likewise, the λLC’s capacity to form inter-chain disulfide is ablated by deleting the C-terminal two amino acid residues containing the penultimate Cys-213. Using this collection of mutant subunits, we then assess their effects on various aspects of IgM biosynthesis in a total of 48 different construct combinations. For each setting, steady state subcellular distribution of involved subunit chain(s) is visualized to detect changes elicited by the mutation or the presence of co-expressed mutant subunit(s). We additionally focus on the induction and prevention of Russell body-related inclusion body phenotypes to elucidate their relations to IgM product quality, assembly intermediate production, and product secretion output. In this combinatorial mutagenesis approach, we unexpectedly demonstrate that not all the prescribed inter-chain disulfide bridge is required for the polymeric IgM formation. In other words, the loss of inter-chain disulfide bonding at specific positions is tolerated for IgM assembly and secretion. This finding illustrates the vital roles of underlying non-covalent protein-protein associations that not only orchestrate the initial subunit interactions and assembly, but also maintain the polymeric IgM product integrity during the ER quality control steps, secretory pathway trafficking, and secretion. We propose that the IgM assembly process is robust and has a stopgap that permits the secretion of polymeric IgM even when some inter-chain disulfide bond formation is incomplete. Our approach successfully combines the analysis of intracellular and extracellular events to holistically illustrate the requirements and exemptions for IgM assembly and secretion. Additionally, this study serves as an inclusive guide on recombinant IgM expression and provides insights into IgM-based novel designs to generate multivalent biologic modalities.

## 2. Materials and methods

### 2.1. Detection antibodies and reagents

Affinity-purified rabbit polyclonal anti-human IgG + IgM (H+L) (cat. 309-005-107) was purchased from Jackson ImmunoResearch Laboratories and was used for Western blotting. Affinity-purified rabbit polyclonal anti-human lambda-light chain (cat. A019302-2) was purchased from Dako and was used for Western blotting. FITC- and Texas Red-conjugated goat polyclonal anti-human mu-heavy chain (cat. 2020-02 and 2020-07) as well as FITC- and Texas Red-conjugated goat polyclonal anti-human lambda-light chain (cat. 2070-02 and 2070-07) were obtained from Southern Biotech and were used for immunofluorescent microscopy. Rabbit polyclonal anti-calnexin (cat. C4731) was from MilliporeSigma. Rabbit polyclonal anti-giantin polyclonal antibody (cat. PRB-114P) was from Covance. Mouse monoclonal anti-human JC (clone 3C7, aka OTI3C7) (cat. TA504168) was obtained from OriGene Technologies. FITC-conjugated mouse monoclonal anti-CD147 (clone HIM6) (cat. 555962) was obtained from BD Biosciences. Unless specifically mentioned, chemical, pharmacological agents, and biological reagents used in this study were obtained from MilliporeSigma.

### 2.2. Expression construct cloning

Amino acid sequence information for a natural human monoclonal IgM known as SAM-6 is publicly available in the US patent publication US20110207917A1. To generate a full-length μHC sequence, we used the VH sequence listed as “SEQ ID NO 18.” Because this VH sequence appeared to lack a portion of the FR1 region, we added back a stretch of missing amino acid sequence to make it free from gloss abnormality before reconstructing a full-length μHC sequence using the constant region reported in UniProt: P01871. To generate a full-length SAM-6 λLC sequence, we selected a VL sequence listed as “SEQ ID NO 14” and fused it to the λLC constant region reported in UniProt: P0DOY2. Although this VL sequence has an unpaired cysteine at position 22 in CDR1, we continued to use the reported original VL sequence because substituting the Cys to Ala or Ser did not make detectable difference in protein expression or protein quality (our unpublished data). Both μHC and λLC mature domain sequences were then fused to a heterologous VK1|O12 signal sequence (see V BASE website, https://www2.mrc-lmb.cam.ac.uk/vbase/) to facilitate ER targeting and entry into the secretory pathway. A full-length JC sequence reported in UniProt: P01591 (GenBank: NM_144646) was used to construct a human JC expression vector. All constructs were generated by gene synthesis and fragment assembly methods. All the recombinant genes of interest used in this study were sequence-verified and cloned into a pTT5 expression vector licensed from the National Research Council of Canada.

### 2.3. Cell culture, transient transfection, and protein production

The HEK293-EBNA1(6E) cell line used in this study was licensed from the National Research Council of Canada. HEK293 cells were cultured in a humidified Reach-In CO2 incubator (ThermoFisher) at 37 °C, 5% CO2 using FreeStyle 293 Expression Medium (ThermoFisher). Cells were maintained in suspension format using disposable shaker flasks placed on Innova 2100 platforms (Eppendorf) rotating at 130 rpm. Expression constructs were transfected into HEK293 cells using a PEI-based protocol described previously [45]. To express and produce recombinant hexameric IgM, a cognate pair of SAM-6 μHC and SAM-6 λLC encoding constructs or mutant constructs were co-transfected at a plasmid DNA weight ratio of 1-to-1. For the pentameric IgM expression, 3 subunit chains (i.e., SAM-6 μHC, SAM-6 λLC and JC) were co-transfected at a plasmid DNA ratio described in the text or the corresponding figure legend. Difco yeastolate cell culture supplement (BD Biosciences) was added to the cell culture at 24 hr post transfection. Cell culture media were harvested on day-7 post transfection and used for SDS-PAGE, Western blotting, and protein purification.

### 2.4. Purification of recombinant human IgM

*<u>Hexameric IgM purification.</u>* Putative hexameric IgM was first purified from the harvested culture medium by affinity chromatography using Capto L resin (Cytiva). It is important to note that although the LC isotype of SAM-6 is λ [46, 47], hexameric SAM-6 IgM is bound to Capto L resin marketed to purify immunoglobulins bearing κLC. See Results secretion for an explanation. Equilibration and washing buffers were 20 mM HEPES, pH 7.4, 150 mM NaCl. The elution buffer was 10 mM acetate, pH 3.5, 150 mM NaCl. IgM-containing Capto L fractions were pooled and subjected to cation exchange chromatography using Nuvia HR-S (Bio-Rad) with an equilibrating and wash buffer (20 mM acetate, pH 5.2, 30 mM NaCl), followed by elution with a linear NaCl gradient using the following elution buffer: 20 mM acetate, pH 5.2, 1 M NaCl. Purified hexameric IgM was concentrated and dialyzed into a formulation buffer (10 mM acetate, 0.26 M sucrose, pH 5.2). The quality of purified hexameric IgM was assessed by analytical SEC using a Superdex 200 10/300 GL column (Cytiva) using the formulation buffer mentioned above. *<u>Pentameric IgM purification.</u>* Putative pentameric IgM was first purified from the harvested culture medium by affinity chromatography using HiTrap Protein L (Cytiva). Like hexameric IgM, pentameric IgM is also bound to the HiTrap Protein L column mainly marketed to capture κLC-bearing immunoglobulins [46, 47]. The bound IgM was eluted from the column using 140 mM acetic acid, pH 2.8, and the eluate was neutralized by adding 1 M Tris-HCl, pH 8.0. IgM-containing fractions were pooled and diluted in 50 mM sodium acetate, pH 5.2 and loaded onto HiTrap S HP (Cytiva). IgM was eluted from the CEX column using a linear salt gradient in 50 mM sodium acetate, pH 5.2, followed by concentration and dialysis into 50 mM sodium acetate, pH 5.2, 200 mM NaCl. The quality of purified pentameric IgM was assessed by analytical SEC using HPLC (Agilent 1200 series) on a Yarra 3 µm SEC-2000 LC 300 x 4.6 mm column (Phenomenex) in 50 mM Tris-HCl, pH 7.0, 0.5 M arginine, 0.05% sodium azide.

### 2.5. Size exclusion chromatography coupled to multi-angle light scattering (SEC-MALS)

A TSKgel G4000SWXL column, 7.8 × 300 mm, 8-μm bead, 450-Å pore (TOSOH BIOSCIENCE) and an Agilent 1260 Infinity II HPLC system were used to determine the oligomeric state of purified IgM. The HPLC was coupled to a Wyatt miniDAWN multi-angle light scattering (MALS) detector and a Wyatt Optilab rEX refractive index detector. Protein samples at a concentration of ∼0.5 mg/mL (100 μL) were loaded on the column. All experiments were conducted at room temperature at a 0.5 mL/min flow rate in PBS, pH 7.0. The MW and mass distribution of the sample were then determined using the ASTRA software version 6.1.7.17 (Wyatt Technology).

### 2.6. Immunofluorescent microscopy

To perform imaging experiments, transfected cells grown in suspension format in shaker flasks were seeded onto poly-D-lysine coated glass coverslips at 48 hr post-transfection. Seeded cells were then statically cultured for 24 hr before the cells were fixed at 72 hr post transfection, using 0.1 M sodium phosphate buffer (pH 7.4) containing 4% paraformaldehyde for 30 min at room temperature. In some experiments, transfected cells were cultured in a growth medium containing 15 μg/mL Brefeldin A for 24 hr to block ER-to-Golgi transport before the cell fixation. After washing and quenching steps in PBS containing 0.1 M glycine, fixed cells were permeabilized in PBS containing 0.4% saponin, 1% bovine serum albumin, 5% fish gelatin for 15 min, followed by incubation with designated primary antibodies in the permeabilization buffer for 60 min. After three washes in the permeabilization buffer, the cells were stained with secondary antibodies for 60 min in the permeabilization buffer. Coverslips were mounted to microscope slide-glass using Vectashield mounting media (Vector Laboratories) and cured overnight at 4°C. The slides were analyzed on a Nikon Eclipse 80i microscope or Eclipse Ti-E microscope with a 60× or 100× CFI Plan Apochromat oil objective lens and Chroma FITC-HYQ or Texas Red-HYQ filter. Images were acquired using a Cool SNAP HQ2 CCD camera (Photometrics) and Nikon NIS-Elements imaging software.

### 2.7. SDS-PAGE and Western blotting

On day-7 post transfection, aliquots of suspension cell culture were withdrawn from shake flasks, and the culture media were separated from cell pellets by centrifugation at 1,000 g for 5 min. Harvested cell culture media were mixed with 2× NuPAGE lithium dodecyl sulfate (LDS) sample buffer (ThermoFisher). Cell lysate samples were prepared by lysing the cell pellets directly in 1× LDS sample buffer. To prepare samples under reducing conditions, 5% (v/v) beta-mercaptoethanol was included in the LDS sample buffer. To prepare samples under non-reducing conditions, 2 mM N-ethylmaleimide was included as an alkylating agent to prevent disulfide exchange reactions. All samples were heated at 75°C for 5 min. To normalize the cell lysate sample loading, whole cell lysates corresponding to 12,000-12,500 cells were analyzed per lane. A sample volume equivalent to 5 μl of the harvested culture medium was analyzed per lane to compare the differences in volumetric secretion titers. Unless specifically mentioned in the figure legend, SDS-PAGE was performed using NuPAGE 4–12% Bis-Tris gradient gel with a compatible MES SDS buffer system (both from ThermoFisher). Resolved proteins were electro-transferred to a nitrocellulose membrane, blocked, and probed with primary antibodies of choice. After three washes in PBS containing 0.05% (v/v) Tween-20, the nitrocellulose membranes were probed with AlexaFluor680-conjugated secondary antibodies (ThermoFisher). After three rounds of washing, the membranes were scanned to acquire Western blotting data using the Odyssey infrared imaging system (LI-COR Biosciences).

### 2.8. Negative-stain transmission electron microscopy

Grids covered with an amorphous carbon film (01843-F, Ted Pella) were glow discharged for 15 mA and 45 sec using PELOCO easiGlow. Protein sample solution (0.1 mg/mL, 3 μL) was applied to the treated grids and stained with 0.1% uranyl formate before imaged under Talos electron microscope operated at 200 kV. In total, 597 images for the hexameric IgM sample and 406 images for the pentameric IgM sample were recorded on the K3 detector using the Latitude S software package (Gatan Inc.) at nominal magnification 22,000× corresponding to 1.84 Å/pixel. For detailed analysis, 88,125 hexameric IgM particles and 52,657 pentameric IgM particles were picked on cisTEM [48] and exported to RELION [49] for 2D classification.

## 3. Results

### 3.1. Biosynthesis of model human IgM SAM-6 takes place without a major secretory bottleneck in HEK293 cells

To study the process of IgM biosynthesis in the most straightforward experimental system, we first co-expressed the μHC and λLC subunit of a natural human IgM SAM-6 in HEK293 cells. The third component, the JC subunit, will be brought into the assembly reaction later in Section 3.6. To detect if there is any overt trafficking bottleneck along the secretory pathway during SAM-6 overexpression, the subcellular distribution of μHC and λLC was visualized by immunofluorescent microscopy. At steady state, both μHC and λLC subunits were detected not only in the ER-like perinuclear structure and Golgi-like juxtanuclear region (Fig. 1A) but also in punctate structures distributed broadly in the cytoplasm (Fig. 1A). The punctate staining was more readily detectable when the recombinant expression level was high, when cells spread more flatly on coverslips, or when multiple focal planes were examined for a given image field. Even after a prolonged Brefeldin A (BFA) treatment to pharmacologically block the ER-to-Golgi cargo transport, the intracellular pool of SAM-6 did not show signs of aberrant accumulation that led to notable inclusion body formation (see Fig. 1B).

**Figure 1.**
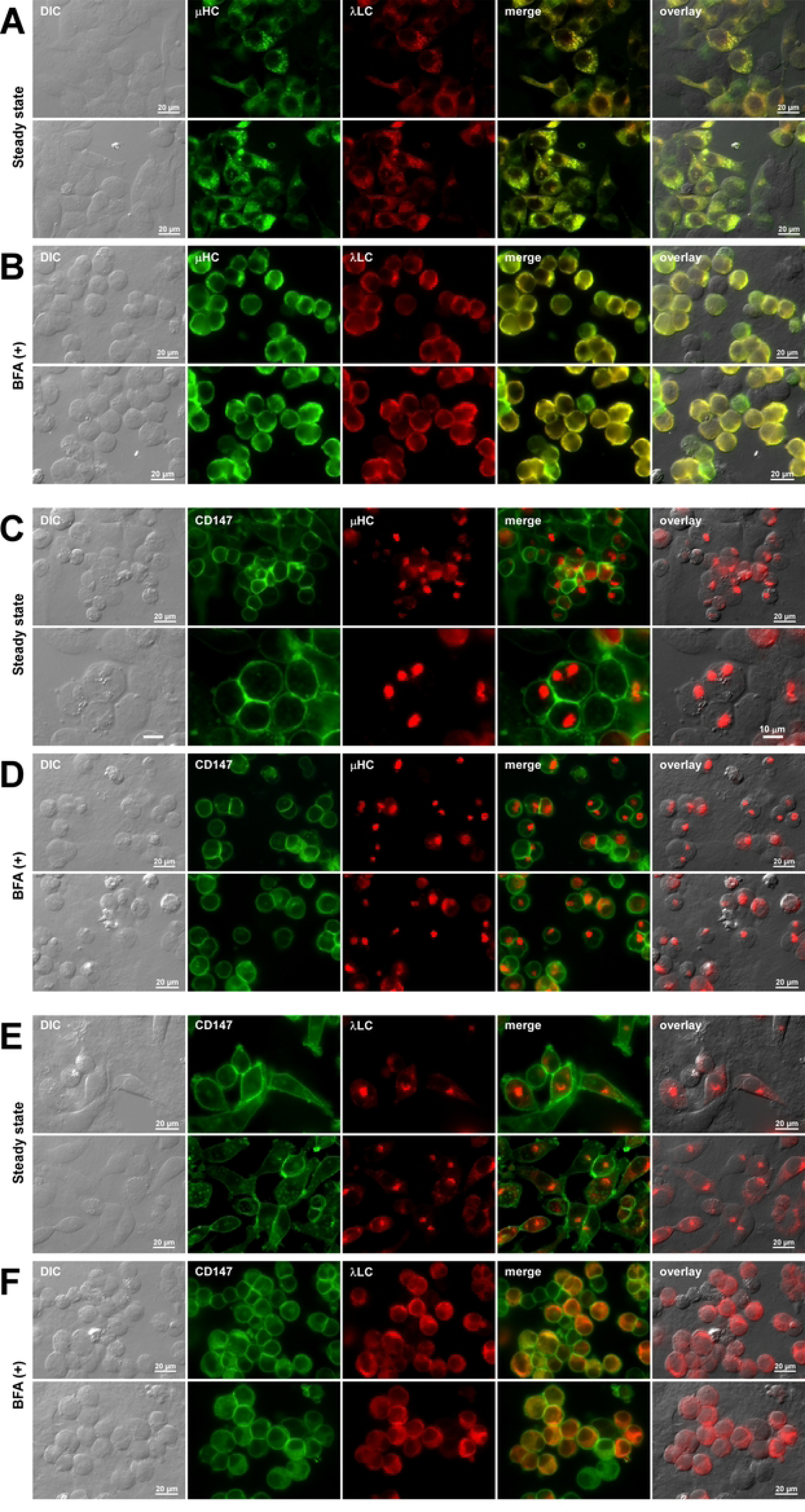
Steady state subcellular distribution of co-expressed μHC and λLC subunits during hexameric IgM product expression in HEK293 cells. Fluorescent micrographs of HEK293 cells transfected with (A, B) SAM-6 μHC and SAM-6 λLC construct pair, (C, D) SAM-6 μHC subunit alone, or (E, F) SAM-6 λLC subunit alone. On day-2 post-transfection, transfected cells were seeded onto poly-D-lysine coated coverslips in the absence (A, C, E) or presence (B, D, F) of 15 μg/mL Brefeldin A (BFA) and cultured statically for 24 hr. On day-3 post-transfection, cells were fixed, permeabilized, and immunostained. (A, B) Co-staining was performed using FITC-labeled anti-human μHC antibody and Texas Red-labeled anti-human λLC antibody. (C, D) Co-stained with FITC-labeled anti-CD147 antibody and Texas Red-labeled anti-human μHC antibody. (E, F) Co-stained with FITC-labeled anti-CD147 antibody and Texas Red-labeled anti-human λLC antibody. Green and red image fields were superimposed to create ‘merge’ views. DIC and ‘merge’ were superimposed to make ‘overlay’ views in A and B. DIC and red image fields were superimposed to create ‘overlay’ views in C‒F.

We next examined the intracellular behaviors of individual subunit chains. Similar to what has been shown for γHC subunits derived from various IgG antibodies [50–56], SAM-6’s μHC subunit induced globular spherules, referred to as Russell bodies, in almost all the μHC expressing cells (Fig. 1 C D). This was true when the cells were cultured under normal cell growth condition (Fig. 1C) or after a prolonged BFA treatment (Fig. 1D). By contrast, SAM-6’s λLC subunit distributed mainly to a juxtanuclear Golgi-like structure and to a lesser extent to the ER at normal growth condition (Fig. 1E). Unlike the extensively aggregated μHC protein that formed Russell body, the Golgi localization of λLC was sensitive to BFA treatment which expectedly redistributed the λLC to the ER-like localization (Fig. 1F).

To study the relationship between the steady state subcellular cargo distribution and the actual protein secretion outputs, we examined the secreted product accumulation in cell culture media at day-7 post-transfection. When the [μHC + λLC] construct pair was co-expressed, both subunit chains were readily detectable in the culture media when analyzed under reducing conditions (Fig. 2 A B, lane 1). Under non-reducing conditions, a large protein complex corresponding to a covalently assembled putative hexameric IgM was readily detectable as a major product just beneath the loading well (Fig. 2 A B, lane 4, red arrowhead) along with a couple of discrete assembly intermediates and free λLC monomers and dimers (Fig. 2 A B, lane 4, see labels for the assembly intermediate species). Secretion of assembly intermediates was suggested to be a normal and universal consequence of the IgM biosynthetic process [57], and our results agreed with such characteristics. Unlike the human IgG counterpart, there was no convenient assay that could directly and specifically quantitate the amount of assembled hexameric IgM in harvested cell culture media, but a comparison against a known amount of IgG antibodies in Coomassie blue stained gels estimated that a volumetric titer of secreted SAM-6 was consistently in the 25‒50 mg/L range including both the polymeric IgM and assembly intermediates. Given that the known secretion output for recombinant IgGs can vary from ∼1 mg/L for poor-secreting IgG mAbs and up to ∼300 mg/L for high-secreting IgG mAbs in this HEK293 transient expression system [50, 53, 54, 56], the observed 25‒50 mg/L secretion titer without any optimization was respectable. Although plasma cell-specific ER resident chaperones are proposed to specialize in assisting the assembly of IgMs [58–60], a non-professional secretory cell like HEK293 cell can also assemble and secrete this class of complex secretory cargo without overt difficulties.

**Figure 2.**
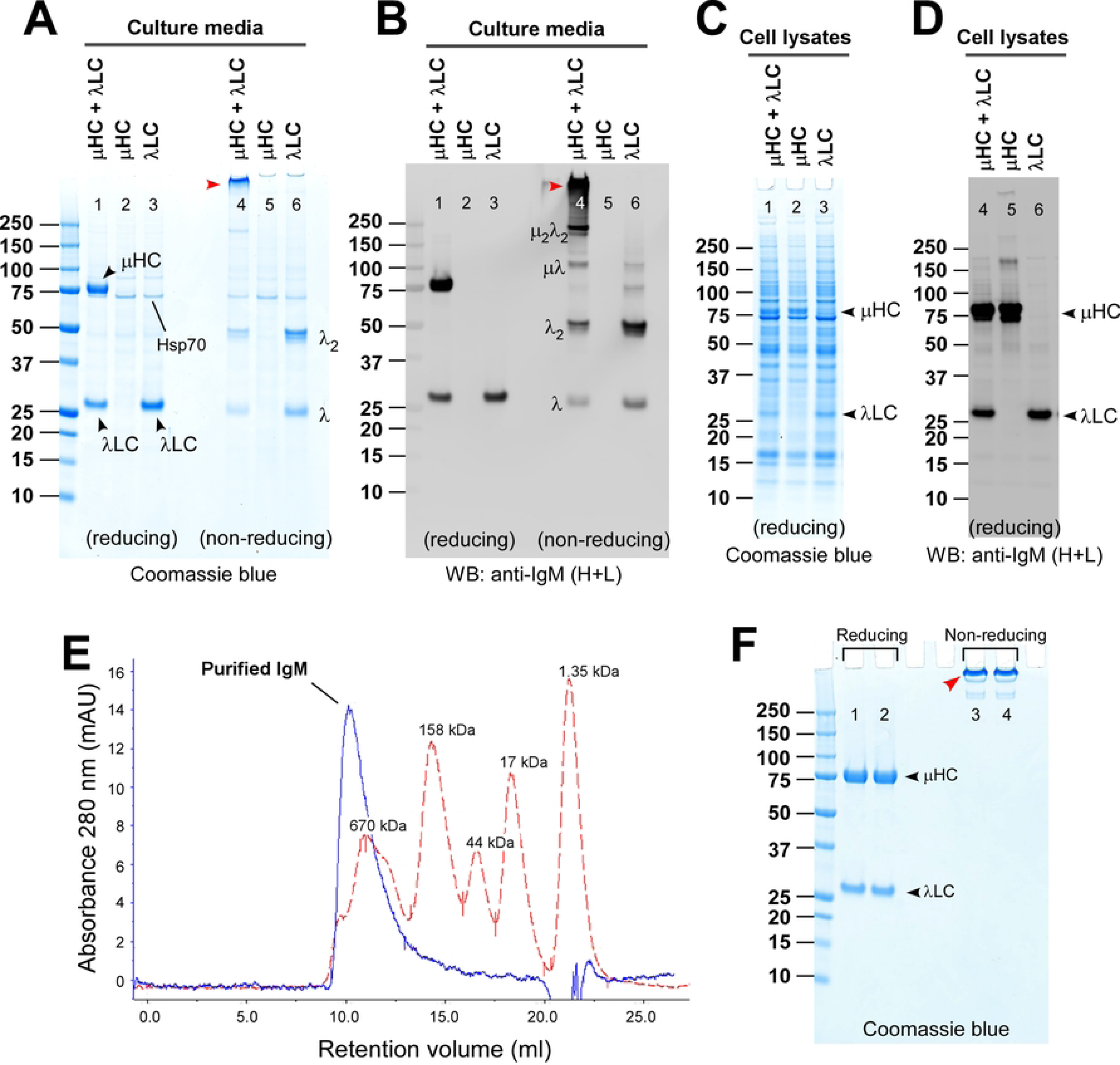
Recombinant expression and purification of hexameric SAM-6 IgM. (A) Coomassie blue stained gel showing the secreted IgM and subunits. HEK293 cells were transfected with the μHC and λLC construct pair (lanes 1 and 4), μHC construct only (lanes 2 and 5), or λLC construct only (lanes 3 and 6). Cell culture media were harvested on day-7 post-transfection and analyzed by SDS-PAGE under reducing conditions (lanes 1‒3) or non-reducing conditions (lanes 4‒6). The detectable subunit chain is pointed by an arrowhead and labeled in lanes 1‒3. Assembled hexameric IgM is pointed by a red arrowhead in lane 4. Monomeric and dimeric free λLC subunit is labeled in lane 6. A cell host-derived ∼74 kDa protein visible in all culture media lanes was identified as human Hsp70 by mass spectrometry (data not shown). (B) An identical sample set was analyzed by Western blotting. The blotted membrane was probed with polyclonal anti-IgM (H+L) antibodies to simultaneously detect both subunits and assembly intermediates composed of μHC or λLC or both. The expected protein band corresponding to the assembled hexameric IgM is pointed by a red arrowhead in lane 4. Identifiable assembly intermediates are labeled next to the corresponding protein bands in lane 4. Cell lysate samples were prepared on day-7 post-transfection and analyzed by (C) SDS-PAGE or (D) Western blotting after the proteins were resolved under reducing conditions. Subunit chains are pointed by arrowhead and labeled. (E) The overall purity of two-step purified hexameric IgM was assessed by analytical SEC. SEC profile of hexameric SAM-6 IgM is shown in a solid blue line. The size marker reference is shown in a dotted red line. (F) The purified hexameric SAM-6 IgM was analyzed by SDS-PAGE under reducing (lanes 1‒2) or non-reducing (lanes 3‒4) conditions. Sample loading was 2.5 μg per lane. The subunit chain is pointed by arrowhead and labeled (lanes 1‒2). Fully assembled hexameric IgM is pointed by a red arrowhead in lanes 3‒4.

Extensive Russell body formation in μHC-expressing cells (Fig. 1 C D) had alluded to a minimal free μHC secretion. In good agreement with such phenotypes, μHC was secretion-incompetent by itself, and thus, free μHC was hardly detectable in the cell culture media (Fig. 2 A B, lanes 2 and 5). By contrast, as implied by its predominant Golgi distribution at steady state (see Fig. 1E), the free λLC subunit was abundantly secreted to the culture media as a mixture of monomers and disulfide-linked dimers (Fig. 2 A B, lanes 3 and 6). Whole cell lysates were also analyzed by Western blotting to compare the relative expression levels of each subunit in different transfection settings (Fig. 2 C D). Expression level of μHC and λLC was high enough to be detectable by Coomassie blue staining (Fig. 2C, arrowheads) irrespective of whether μHC and λLC were co-transfected (Fig. 2C, lane 1) or transfected individually (Fig. 2C, lanes 2 and 3).

From the morphological and biochemical viewpoints, intracellular behaviors of SAM-6 μHC were very similar to those of various γHC subunits characterized in our laboratory using similar methods [50–56]. Namely, (1) full-length μHC had a strong propensity to induce Russell body by itself in the absence of LC expression, (2) LC co-expression rescued the μHCs from aggregating into Russell body, and (3) μHC acquired a secretion competency upon pairing with the LC subunit. Although the experiment itself was simple, the general guiding principles of immunoglobulin assembly were recapitulated precisely for the model IgM SAM-6.

To conduct various biochemical studies on IgMs in the subsequent sections, we produced and purified a putative hexameric form of SAM-6 (later shown as a bona fide hexameric IgM; see Section 3.7). Using the transfection conditions employed above, we carried out a small-scale production run and purified the IgM from 500 mL of the harvested culture medium. Although the LC isotype of SAM-6 IgM was λ, we found inadvertently that SAM-6 IgM can be purified using Capto^TM^ L resin (Cytiva) marketed to purify immunoglobulins with κLC isotype. Although there are reports on the weak binding of λLC to Protein L [46, 47], precise reasons why SAM-6 IgM could bind Capto^TM^ L resin were unclear (perhaps avidity?). Regardless, the binding was reproducible in that we were also able to purify a putative pentameric SAM-6 IgM (with a JC) using HiTrap^TM^ protein L resin (Cytiva), another affinity resin marketed to purify immunoglobulins bearing κLCs (see below, Section 3.7). Although unexpected, we took advantage of this finding to carry out the initial affinity capture step. As a second step, we performed cation exchange chromatography (see Materials and Methods). The purified yield was roughly 15 mg from the 500 mL culture medium. A representative analytical SEC chromatogram for the two-step purified IgM is shown in Fig. 2E (see also Fig. 7B). Purified IgM was also analyzed by SDS-PAGE both under reducing and non-reducing conditions and confirmed the expected redox-dependent subunit chain dissociation and gel mobility change (Fig. 2F).

### 3.2. Deletion of the CH1 domain alone does not make SAM-6 μHC subunit secretion competent

CH1 domain of the HC subunit is known to play critical roles in regulating the immunoglobulin biosynthesis at least by two mechanisms—the retention of free HCs by a BiP-mediated ER quality control mechanism and the covalent assembly with LC subunit via disulfide bond formation [61, 62]. It was demonstrated that CH1 domains remain largely unstructured and are bound by BiP until the BiP is displaced by a folded LC subunit [63, 64]. The displacement of BiP by the LC induces a spontaneous folding of the unstructured CH1 domain, followed by a disulfide linkage between the Cys residue in the CH1 domain and the Cys residue near the C-terminus of the LC subunit [65]. Therefore, the binding of BiP to the CH1 underpins the ER quality control mechanism that prevents the secretion of free HCs from the cells until the HCs assemble with the cognate LCs [65]. Despite such critical importance, the CH1 domain deletion can occur naturally via errors in gene recombination in animal models and homologous cellular systems [66]. In the case of IgGs, such CH1 deletion (often denoted as ΔCH1 mutation) alone renders the mutant HCs secretion-competent by bypassing the BiP-mediated ER quality control. Consequently, the mutant ΔCH1 HC dimers are secreted freely to the extracellular space even in the absence of LC expression or without assembling with LC subunits [66].

To understand the roles of the CH1 domain in μHC subunit biosynthesis and IgM assembly, we made a mutant μHC construct named μHC-ΔCH1 (Fig. 3A, third row) that lacked the entire CH1 domain (a.k.a., Cμ1) by deleting a corresponding stretch of amino acid residues as defined in UniProt: <u>P01871</u>. We then assessed the steady state subcellular distribution, Russell body formation, IgM assembly, and product secretion of this μHC-ΔCH1 mutant by expressing itself alone or by co-expressing with the λLC. Firstly, the morphology of μHC-ΔCH1 expressing cell became rounded, and the μHC-ΔCH1 protein aggregated into Russell body in those cells (Fig. 3B, second and third rows). Both the cell morphology and Russell body forming characteristics were indistinguishable from those of the intact μHC expressing cells (Fig. 3B, first row; see also Fig. 1C). Secondly, in contrast to γHC’s ΔCH1 mutants, the CH1 deletion alone did not make the μHC-ΔCH1 protein secretion-competent. As such, μHC-ΔCH1 protein was not detectable in culture media by Coomassie blue staining (Fig. 3D, lane 2) or by Western blotting (Fig. 3E, lane 2) despite the abundant expression of μHC-ΔCH1 protein in cell lysates (Fig. 3 D E, lane 6). Both the level of protein synthesis and the complete absence of protein secretion were indistinguishable between the intact μHC and μHC-ΔCH1 mutant (Fig. 3 D E, compare lanes 1 and 2; lanes 5 and 6). The only recognizable difference between them was their gel mobility (compare lanes 5 and 6 in Fig. 3E). Thirdly, upon co-expressing with λLC, the presence of λLC in the same ER milieu not only prevented the μHC-ΔCH1 mutant from aggregating into Russell body (Fig. 3C, second and third rows) but also changed its steady state subcellular distribution to the ER and cytoplasmic puncta that was similar to that of intact [μHC + λLC] co-expression setting (Fig. 3C, first row). Prevention of μHC-ΔCH1 Russell body formation by the co-expressed λLC suggested that their non-covalent interactions exerted a strong enough effect to discourage μHC-ΔCH1 aggregation (Fig. 3C, second and third rows). These results agreed with what Corcos et al. (2010) reported in their knock-in mouse models [67]. On the surface, the prevention of Russell body formation in [μHC-ΔCH1 + λLC] co-expressing cells disagreed with two influential reports [68, 69] where Sitia’s group conducted a similar transfection experiment and found an extensive Russell body formation. As shown below in Fig. 9F (top row), however, we were able to “restore” such extensive Russell body phenotypes in [μHC-ΔCH1 + λLC] expressing cells by additionally co-transfecting the JC subunit (see below). Fourthly, although Russell body phenotype was suppressed in [μHC-ΔCH1 + λLC] co-expression, μHC-ΔCH1 mutant remained secretion incompetent because it still cannot assemble with λLC. As a result, while free λLC was secreted as if it was expressed alone (Fig. 3 D E, lane 4), μHC-ΔCH1 secretion was prevented, except that a disulfide-linked μHC-ΔCH1 dimer was very faintly detectable by Western blotting (Fig. 3 E and G, lane 4, pointed by black arrowhead).

**Figure 3.**
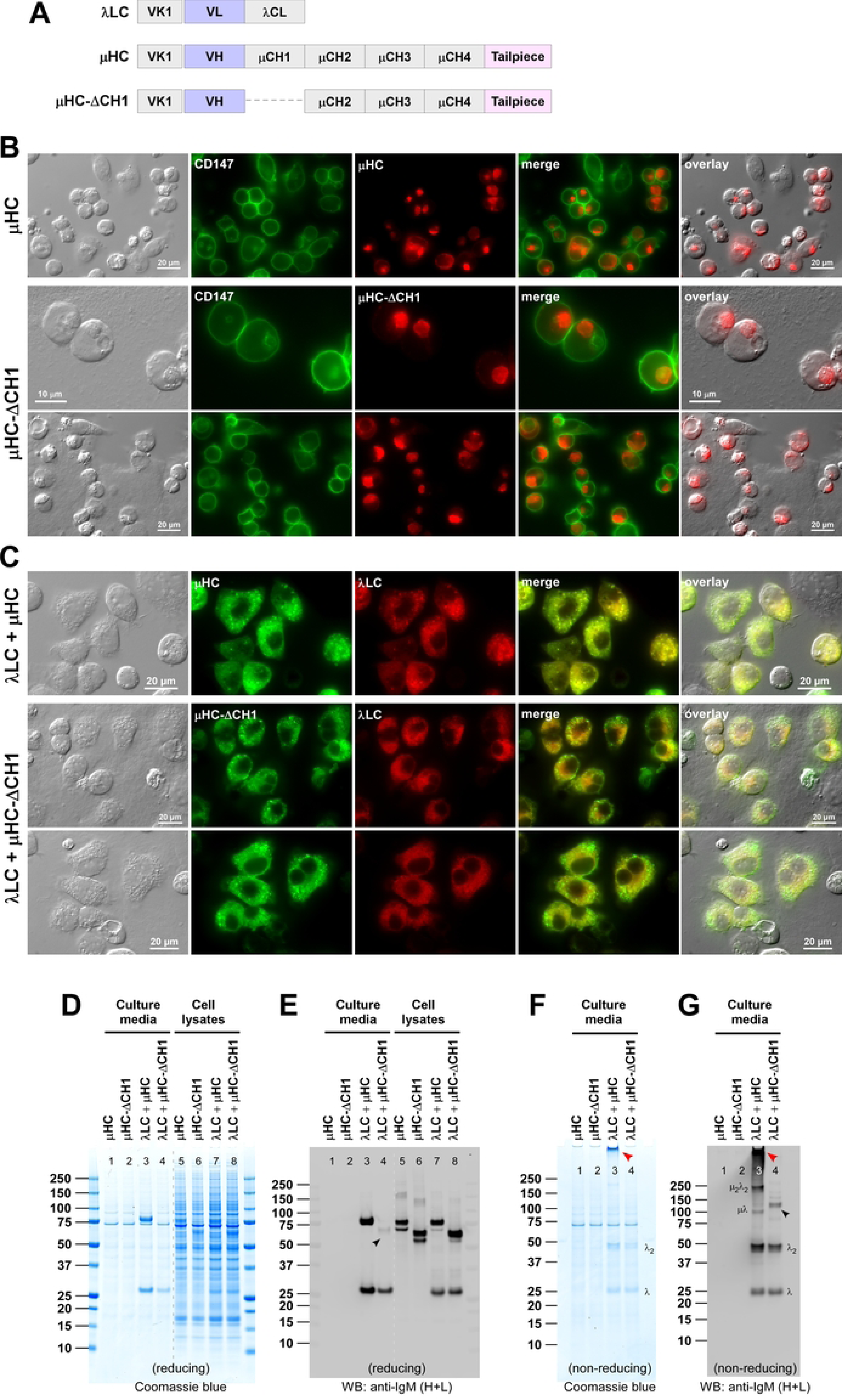
Effects of CH1 domain deletion on hexameric IgM assembly and secretion. (A) Schematic representation of SAM-6 λLC subunit (top), SAM-6 μHC subunit (middle), and SAM-6 μHC-ΔCH1 mutant which lacks the CH1 domain (bottom). The deleted CH1 domain is shown as a dotted line. Individual domain names are indicated in each box. ER targeting is driven by a heterologous signal sequence adapted from a VK1 encoding gene. (B) Fluorescent micrographs of HEK293 cells transfected with μHC (first row) or μHC-ΔCH1 mutant (second and third rows). On day-3 post-transfection, cells were fixed, permeabilized, and immunostained with FITC-labeled anti-CD147 antibody and Texas Red-labeled anti-human μHC antibody. Green and red image fields were superimposed to create ‘merge’ views. DIC and red image fields were superimposed to create ‘overlay’ views. (C) Fluorescent micrographs of HEK293 cells transfected with [λLC + μHC] pair (first row) or [λLC + μHC-ΔCH1] pair (second and third rows). Immunostaining was performed using FITC-labeled anti-human μHC antibody and Texas Red-labeled anti-human λLC antibody. Green and red image fields were superimposed to create ‘merge’ views. DIC and ‘merge’ were superimposed to create ‘overlay’ views. (D‒G) HEK293 cells were transfected with μHC (lanes 1 and 5), μHC-ΔCH1 (lanes 2 and 6), [λLC + μHC] pair (lanes 3 and 7), or [λLC + μHC-ΔCH1] pair (lanes 4 and 8). Cell culture media were harvested at day-7 post-transfection and analyzed by SDS-PAGE under reducing conditions (D, E; lanes 1‒4) or non-reducing conditions (F, G; lanes 1‒4). Cell lysate samples were also prepared on day-7 post-transfection and analyzed by SDS-PAGE (D, lanes 5‒8) or Western blotting (E, lanes 5‒8) after resolving the proteins under reducing conditions. Western blotting was performed using polyclonal anti-IgM (H+L) to detect both μHC and λLC subunits simultaneously as well as assembly intermediates composed of μHC or λLC or both. A faintly detectable μHC-ΔCH1 is pointed by a black arrowhead (E, lane 4). Likewise, faintly detectable μHC-ΔCH1 covalent dimers are pointed by black arrowhead (G, lane 4). The assembled hexameric IgM product is pointed by a red arrowhead (F, G; lane 3). Identifiable assembly intermediates are labeled next to the corresponding bands in panels F and G.

### 3.3. SAM-6 μHC-ΔCH1 is freed from ER retention when combined with C414/575S double mutation

Because of the presence of an 18-residue C-terminal tailpiece that drives the polymerization of IgMs, the biosynthesis of IgM is known to be more complex than that of IgGs [70, 71]. Because the penultimate Cys residue in the tailpiece of μHC is a known substrate for a free thiol-mediated ER retention mechanism [57, 72], this may explain why a mere deletion of the CH1 domain alone (which is a substrate for BiP-dependent ER retention mechanism) did not make the μHC subunit secretion competent (see above Fig. 3, and [62]).

To test whether the lack of μHC secretion was caused by an interplay between these two ER retention mechanisms, we combined the CH1 deletion and Cys-to-Ser mutations to disable BiP-mediated and free thiol-mediated retention simultaneously. In this scheme, the penultimate Cys-575 in the secretory tailpiece was mutated to Ser residue with or without an additional mutation to the Cys-414 located in the CH3 domain that also plays a role in inter-chain disulfide formation between the two adjacent protomers (see illustration of Fig. 4A, rows 1–3) [73]. Furthermore, to study the role of these two Cys residues in conjunction with the BiP-mediated ER retention, the same combination of Cys-to-Ser mutations was also introduced to the μHC-ΔCH1 backbone (see illustration of Fig. 4A, rows 4–6).

**Figure 4.**
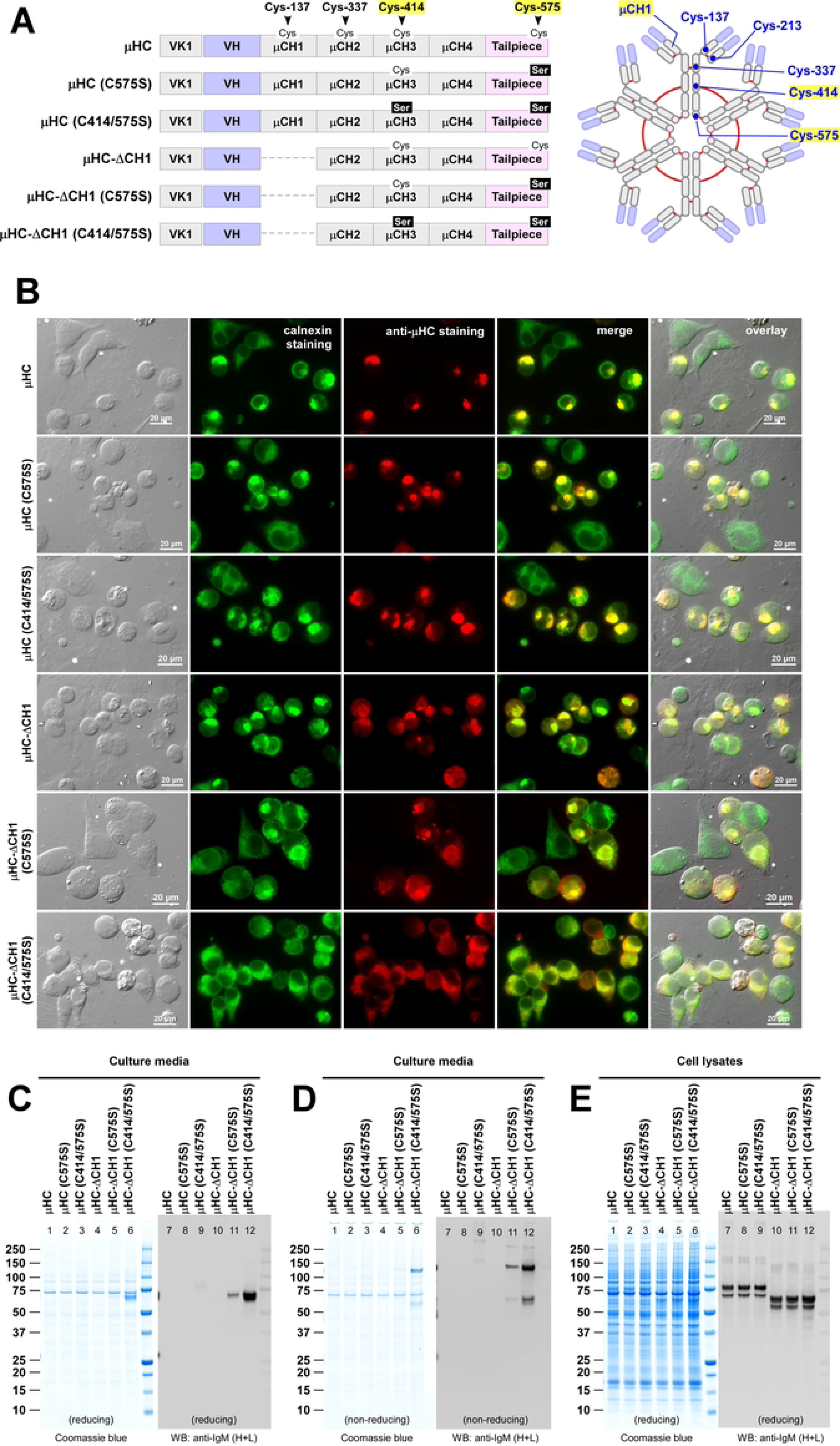
Roles of Cys-414 and Cys-575 residues in μHC subunit synthesis and secretion. (A) Schematic representation of parental SAM-6 μHC (top) and its CH1 deletion mutant μHC-ΔCH1 (fourth row) as well as their C575S and C414/575S mutant variants. The positions of key cysteine residue participating in the inter-chain disulfide bond formation are marked on the parental μHC. The deleted CH1 domain is indicated by a dotted line. Individual domain names are shown in respective grey boxes. The N-terminal “VK1” represents a heterologous signal sequence adapted from the VK1 encoding gene. (A, right diagram) The positions of cysteine residue involved in inter-chain disulfide bond formation are illustrated in the context of the hexameric IgM diagram. Solid red lines represent the inter-chain disulfide bonds and their connectivity. (B) Fluorescent micrographs of HEK293 cells transfected with the six constructs shown in panel A, left. The transfected construct name is shown on the left side of each row. On day-3 post-transfection, cells were fixed, permeabilized, and immunostained with polyclonal anti-calnexin antibody (green) and Texas Red-labeled anti-human μHC antibody (red). Green and red image fields were superimposed to create ‘merge’ views. DIC and ‘merge’ were superimposed to create ‘overlay’ views. (C, D) Cell culture media were harvested at day-7 post-transfection and analyzed by SDS-PAGE and Western blotting after resolving the proteins under reducing (C) or non-reducing (D) conditions. (E) Cell lysate samples were also prepared on day-7 post-transfection and analyzed by SDS-PAGE (lanes 1‒6) or Western blotting (lanes 7‒12) after resolving the proteins under reducing conditions. Western blotting was performed using polyclonal anti-IgM (H+L).

Although C575S point mutation or C414/575S double mutation was introduced to abrogate μHC’s polymerization capacity, both mutant μHCs continued to show high-level Russell body formation that was indistinguishable from the parental μHC (Fig. 4B, rows 1–3, red). Similar to the γHC-induced Russell bodies [50, 51, 53, 55, 56], those μHC globular structures co-aggregated with ER-resident proteins such as calnexin (Fig. 4B, rows 1–3, green and merge) and others (data not shown). In good agreement with the Russell body phenotypes, these mutants were almost entirely retained in the ER and undetectable in the culture media (Fig. 4 C D, lanes 1–3 and 7–9) in spite of abundant protein synthesis in the cell (Fig. 4E, lanes 1–3 and 7–9).

Similarly, C575S single point mutation was not able to prevent μHC-ΔCH1 from aggregating into Russell body (Fig. 4B, fifth row), and the extent of Russell body formation was indistinguishable from that of the parental μHC-ΔCH1 (Fig. 4B, fourth row). Unsurprisingly, the secretion of μHC-ΔCH1 (C575S) remained undetectable by Coomassie blue staining (Fig. 4 C D, lane 5), but its low-level leakage in the form of monomers and dimers was revealed by Western blotting (Fig. 4 C D, lane 11). By contrast, the simultaneous introduction of ΔCH1 and C414/575S not only rescued the μHC from aggregating into Russell bodies (Fig. 4B, sixth row, red) but also finally liberated the μHC-ΔCH1 from the ER retention, thereby resulting in efficient secretion into culture media in a mixture of monomers and covalent-dimers (Fig. 4 C D, lanes 6 and 12). Despite such dramatic secretion level difference, the steady state level of synthesized proteins inside the cell was comparable (Fig. 4E, lanes 4‒6 and 10‒12). In the absence of λLC, the SAM-6 μHC subunit was released from the ER retention only when simultaneously uncoupled from both BiP-mediated and thiol-mediated ER retention mechanisms.

### 3.4. C575S and C414/575S mutations progressively block the formation of polymeric IgM

To evaluate the effect of Cys-to-Ser mutations on various aspects of IgM biosynthesis, such as μHC–λLC assembly, polymerization, Russell body formation, secretion output, product quality, etc., we co-expressed the SAM-6 λLC with each one of the SAM-6 μHC mutants illustrated in Fig. 4A. Firstly, when [μHC (C575S) + λLC] pair was co-expressed, both subunit chains distributed broadly to the ER and cytoplasmic puncta at steady state (Fig. 5A, second row) and it was indistinguishable from that of parental [μHC + λLC] co-expression pair (Fig. 5A, first row). When co-expressed with λLC, the μHC (C575S) subunit was abundantly secreted at a level equivalent to the parental μHC (Fig. 5C, lanes 1–2 and 7–8). However, the μHC (C575S) mutant could no longer covalently assemble the polymeric IgM species as efficiently as the parental μHC (Fig. 5D, lanes 1–2) and, as such, it mainly generated smaller covalently-assembled complexes such as ∼200 kDa μ2λ2 protomers and ∼100 kDa μλ half-molecules (Fig. 5D, lanes 1–2 and 7–8). Although Cys-575 is known to possess essential functions to drive the covalent polymerization of mouse IgM [70, 73, 74], knocking out the Cys-575 residue alone could only partially abolish the covalent assembly of polymeric human IgM. Whether the [μHC (C575S) + λLC] product retains the ability to form polymeric IgM complexes through non-covalent association cannot be assessed in the presence of a denaturant such as SDS. This critical issue will be addressed later using analytical SEC under physiological conditions.

**Figure 5.**
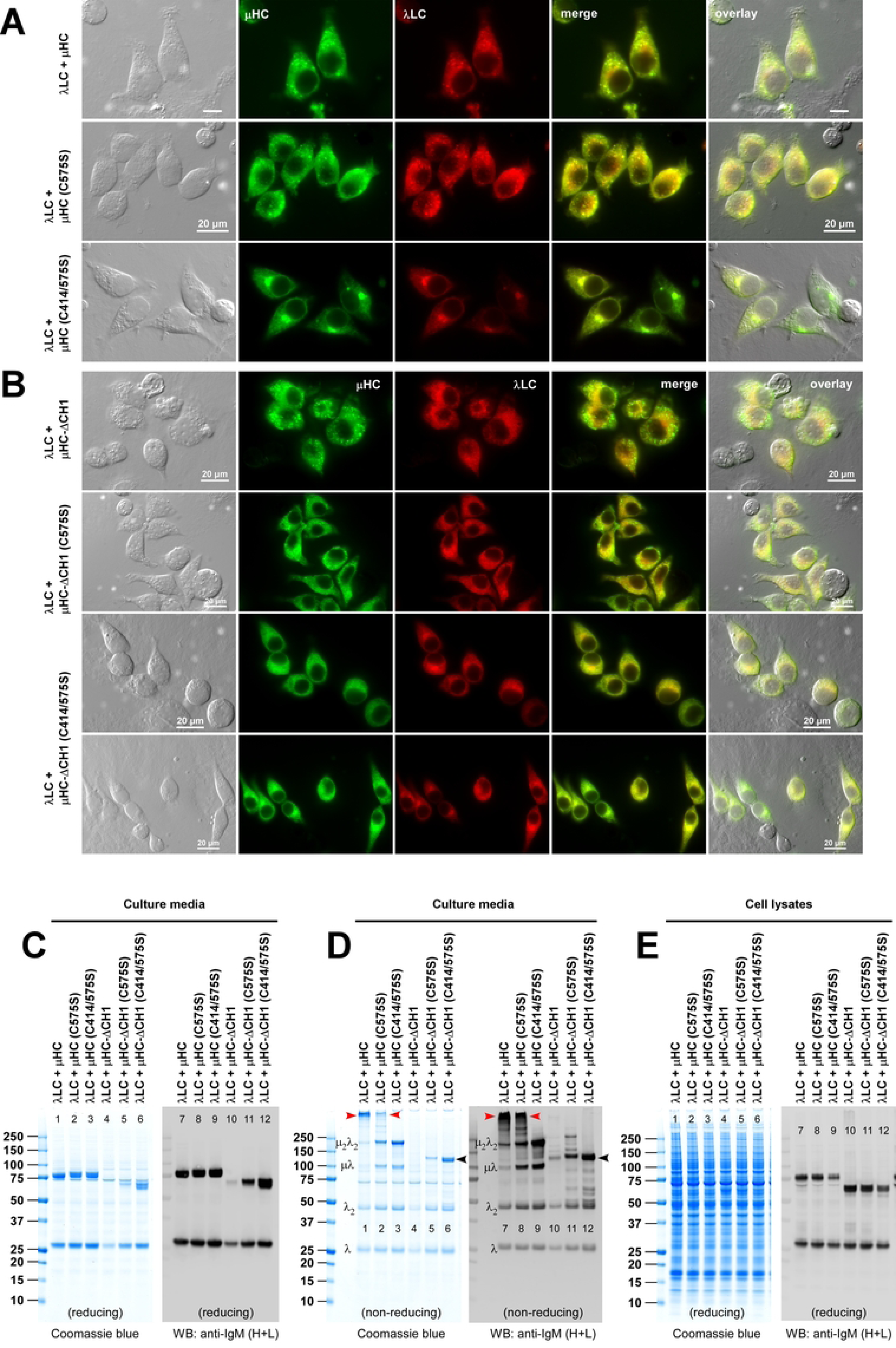
Effect of Cys-414 and Cys-575 mutations on covalent assembly and secretion of hexameric IgM. (A) Fluorescent micrographs of HEK293 cells co-transfected with [λLC + μHC] pair (top row), [λLC + μHC (C575S)] pair (middle row), and [λLC + μHC (C414/575S)] pair (bottom row). On day-3 post-transfection, cells were fixed, permeabilized, and co-stained with FITC-labeled anti-human μHC antibody and Texas Red-labeled anti-human λLC antibody. Green and red image fields were superimposed to create ‘merge’ views. DIC and ‘merge’ were superimposed to create ‘overlay’ views. (B) Fluorescent micrographs of HEK293 cells co-transfected with [λLC + μHC-ΔCH1] pair (top row), [λLC + μHC-ΔCH1 (C575S)] pair (second row), and [λLC + μHC-ΔCH1 (C414/575S)] pair (third and fourth rows). Cell fixation, immunostaining, and image processing were performed as described above. (C‒E) Analysis of protein expression and secreted protein quality for six different subunit co-expression pairs. The transfected construct pair is shown at the top of each lane. Cell culture media (C, D) and cell lysates (E) were harvested at day-7 post-transfection and analyzed by SDS-PAGE and by Western blotting after resolving the proteins under reducing conditions (C and E) or non-reducing conditions (D). Western blotting was performed using polyclonal anti-IgM (H+L) to detect μHC and λLC subunits simultaneously as well as assembly intermediates composed of μHC or λLC or both. A protein band corresponding to the assembled hexameric IgM is pointed by a red arrowhead in panel D, lanes 1, 2, 7, and 8. Likewise, identifiable assembly intermediates are labeled next to the corresponding protein bands in panel D, lanes 1 and 7. Secreted μHC-ΔCH1 (C414/575S) mutant dimers are pointed by black arrowhead in panel D, lanes 6 and 12.

Secondly, when co-expressed with λLC, instead of forming the Russell bodies, μHC (C414/575S) double mutant distributed to a juxtanuclear Golgi-like structure and reticular ER-like structures (Fig. 5A, third row, green), whereas the λLC mainly distributed to the Golgi as if it was expressed by itself (Fig. 5A, third row, red). Although the secretion of the μHC (C414/575S) subunit was comparable to the parental μHC when analyzed under reducing conditions (Fig. 5C, lanes 1 and 3; lanes 7 and 9), secreted product quality was markedly different. When tested under a non-reducing setting, [μHC (C414/575S) and λLC] pair generated ∼200 kDa μ2λ2 protomer species predominantly and could no longer assemble or secrete any covalently polymerized products (Fig. 5D, lanes 3 and 9). Evidently, the μHC (C414/575S) mutant completely lost the ability to polymerize into higher-order species covalently.

### 3.5. C575S and C414/575S mutation progressively increase μHC-ΔCH1 homodimer secretion regardless of λLC co-expression

Because of the missing CH1 domain, μHC-ΔCH1 series constructs cannot assemble covalently with λLC subunit even if λLC was co-expressed; and consistent with this, μHC-ΔCH1 series mutants can only be secreted as covalently linked homodimers at best. Although protein synthesis looked comparable (see Fig. 5E, lanes 4–6 and 10–12), the secretion of μHC-ΔCH1 homodimers varied significantly depending on which Cys residue was mutated (Fig. 5D, lanes 4– 6 and 10–12). While the secretion of μHC-ΔCH1 was only faintly detectable by Western blotting (Fig. 5 C D, lanes 4 and 10; see also Fig. 3 D–G, lane 4), it increased notably when C575S point mutation was introduced (Fig. 5 C D, lanes 5 and 11) and then more profoundly after C414/575S double mutations were introduced (Fig. 5 C D, lanes 6 and 12, black arrowhead). Without the λLC subunit co-expression, both μHC-ΔCH1 and its C575S mutant aggregated into Russell bodies (see above, Fig. 4B); however, when they were co-expressed with the λLC, the co-presence of λLC in the ER lumen effectively prevented the Russell body formation (Fig. 5B, first and second rows). This illustrated that the non-covalent interactions between the variable regions of μHC-ΔCH1 and λLC were sufficient to intervene in the extensive aggregation of μHC-ΔCH1 protein into Russell bodies. Lastly, when both Cys-414 and Cys-575 were mutated simultaneously in the μHC-ΔCH1 background, the μHC-ΔCH1 (C414/575S) mutant distributed to the ER (Fig. 5B, third and fourth rows, green) and co-localized extensively with the co-expressed λLC — and such distribution agreed well with abundant secretion level (Fig. 5 C D, lanes 6 and 12).

### 3.6. J-chain subunit is readily incorporated into polymerizing IgM via disulfide bonds to generate pentameric IgMs

It has been reported that hexameric IgM is preferentially produced when JC subunit availability is limited in the ER lumen [71, 72], while pentameric IgM predominates if sufficient JC subunit is available [17, 75, 76]. However, how much JC is required to tip the balance from hexamer to pentamer formation has not been shown. The fate of excess free JC subunit during IgM biosynthesis is also unknown. To determine the fine line between the hexamer and pentamer IgM formation, we introduced a varying amount of JC-encoding construct into the IgM assembly reaction and then monitored the secretion output and the quality of secreted products.

The JC subunit has eight reactive cysteine residues participating in intra- and inter-molecular disulfide bond formation [77]. Because of such cysteine-rich configuration, it is important to understand the fate and behavior of free JC inside and outside the cells. To this end, we examined JC’s intrinsic aggregation propensity by testing if a JC overexpression leads to intracellular aggregation or if the JC can reach a secretion-competent status without being dependent on other subunit chains. Firstly, Western blotting analysis using a mouse anti-human JC (clone 3C7) detected a ∼20–23 kDa protein in the JC transfected cells but not in the mock-transfected cells (Fig. 6A). Other mAb clones such as 3H7, 3B3, and 2B1 all detected the protein of the same size (data not shown). Although JC is often referred to as a 15 kDa protein consisting of 137 amino acids, due to its rich content of negatively charged amino acids and an N-linked glycosylation site at Asn-49, the JC protein is known to run aberrantly on SDS-PAGE. (On a side note, we found that this anti-JC mAb 3C7 or the rabbit anti-mouse IgG secondary antibody had minor cross-reactivity with λLC and μHC in Western blotting but did not hinder our data interpretation.) Secondly, the mAb 3C7 worked well in immunofluorescent microscopy to visualize the intracellular distribution of transfected JC in HEK293 cells. At steady state, overexpressed JC predominantly filled the ER-like reticular structures that were continuous with the nuclear membrane (Fig. 6B, green) without showing co-localization with giantin, a cis-Golgi marker (Fig. 6B, red). Despite the apparent steady state distribution to the ER, the free JC did not induce notable intra-ER protein aggregates such as Russell body. Thirdly, free JC was released to the culture media by itself (see below), suggesting that it can acquire secretion competency even without assembling with other subunits.

**Figure 6.**
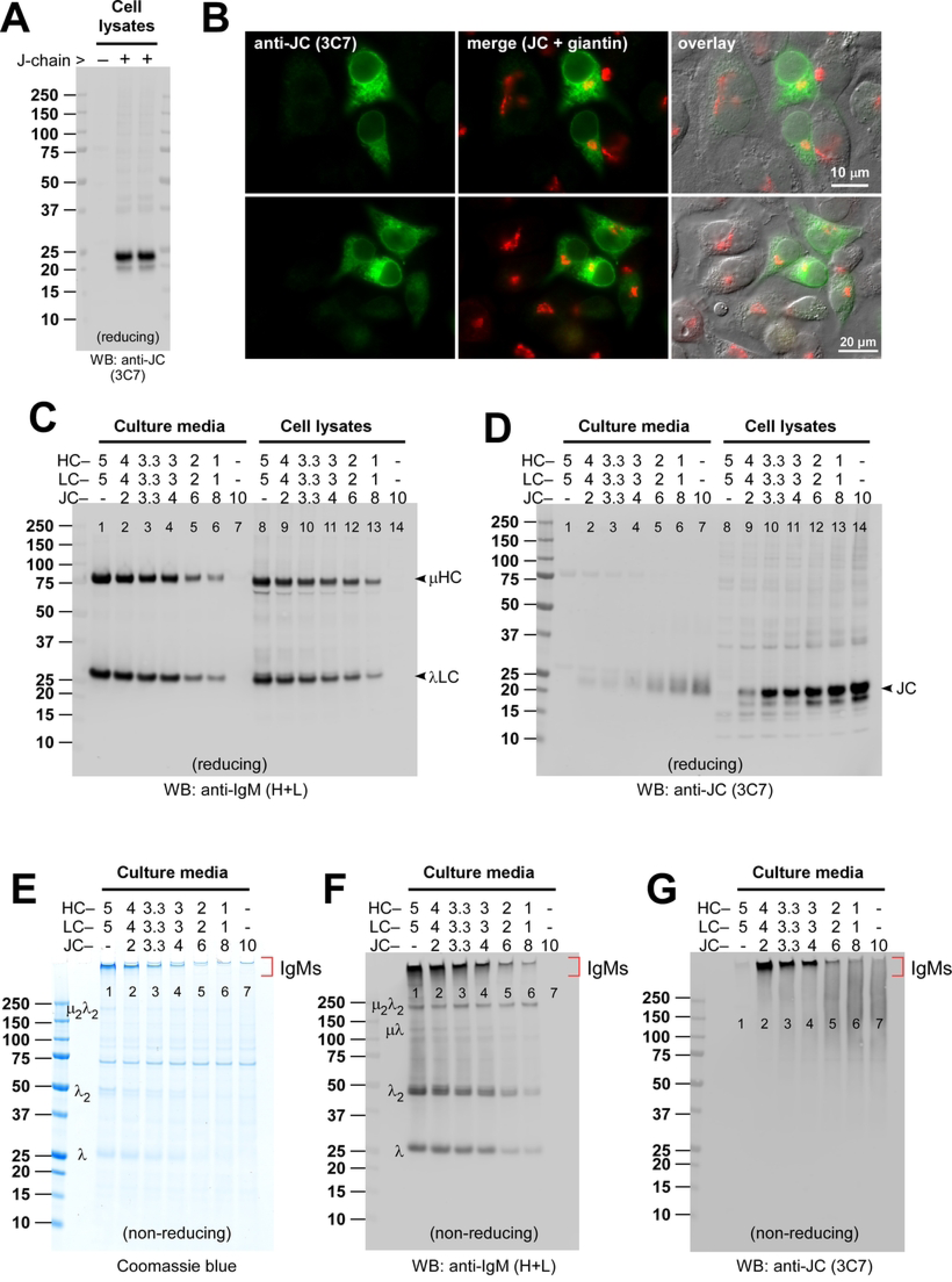
Optimization of pentameric IgM formation by titrating the expression of J-chain subunit. (A, B) HEK293 cells were transfected with a human JC encoding construct, and the expression was verified by Western blotting on cell lysates and by immunofluorescent microscopy on fixed cells. (A) At day-3 post-transfection, cell lysate samples were prepared and analyzed by Western blotting using anti-JC monoclonal antibody (clone 3C7) after resolving the proteins under reducing conditions. Mock-transfected cell lysate was analyzed in parallel as a control for anti-JC detection specificity. (B) On day-3 post-transfection, cells were fixed, permeabilized, and co-stained with anti-JC monoclonal antibody (clone 3C7) (shown in green) and anti-giantin antibody (shown in red). DIC and ‘merge’ images were superimposed to create ‘overlay’ views. (C, D) To produce a pentameric form of IgM, three subunit chains (SAM-6 μHC, SAM-6 λLC, and JC) were co-transfected using varying plasmid DNA ratios indicated at the top of individual lanes without changing the total amount of transfected DNA (= 10 μg). The numbers shown at the top of each lane represent the amount of each construct in μg and should add up to 10. At day-7 post-transfection, cell culture media were harvested (lanes 1‒7), and cell lysates were prepared (lanes 8‒14) to run SDS-PAGE under reducing conditions followed by Western blotting. Blotted membranes were probed (C) with polyclonal anti-IgM (H+L) antibodies to detect both μHC and λLC subunits simultaneously or (D) with monoclonal anti-JC antibody. The detected subunit chain is pointed by an arrowhead and labeled in both panels C and D. (E‒G) Cell culture media harvested at day-7 post-transfection were resolved by SDS-PAGE under non-reducing condition and Coomassie blue stained (E) or analyzed by Western blotting (F, G). Membranes were probed with polyclonal anti-IgM (H+L) to simultaneously detect assembled IgM species and assembly intermediates composed of μHC or λLC or both (panel F) or with monoclonal anti-JC antibody (clone 3C7) to detect JC containing protein complex or free JC (panel G). In panels E and F, protein bands corresponding to λLC monomers, dimers and assembly intermediates are labeled on the left side of lane 1. The protein band corresponding to hexameric or pentameric IgM is marked with a bracket on the right side of lane 7 in panels E‒G.

**Figure 7.**
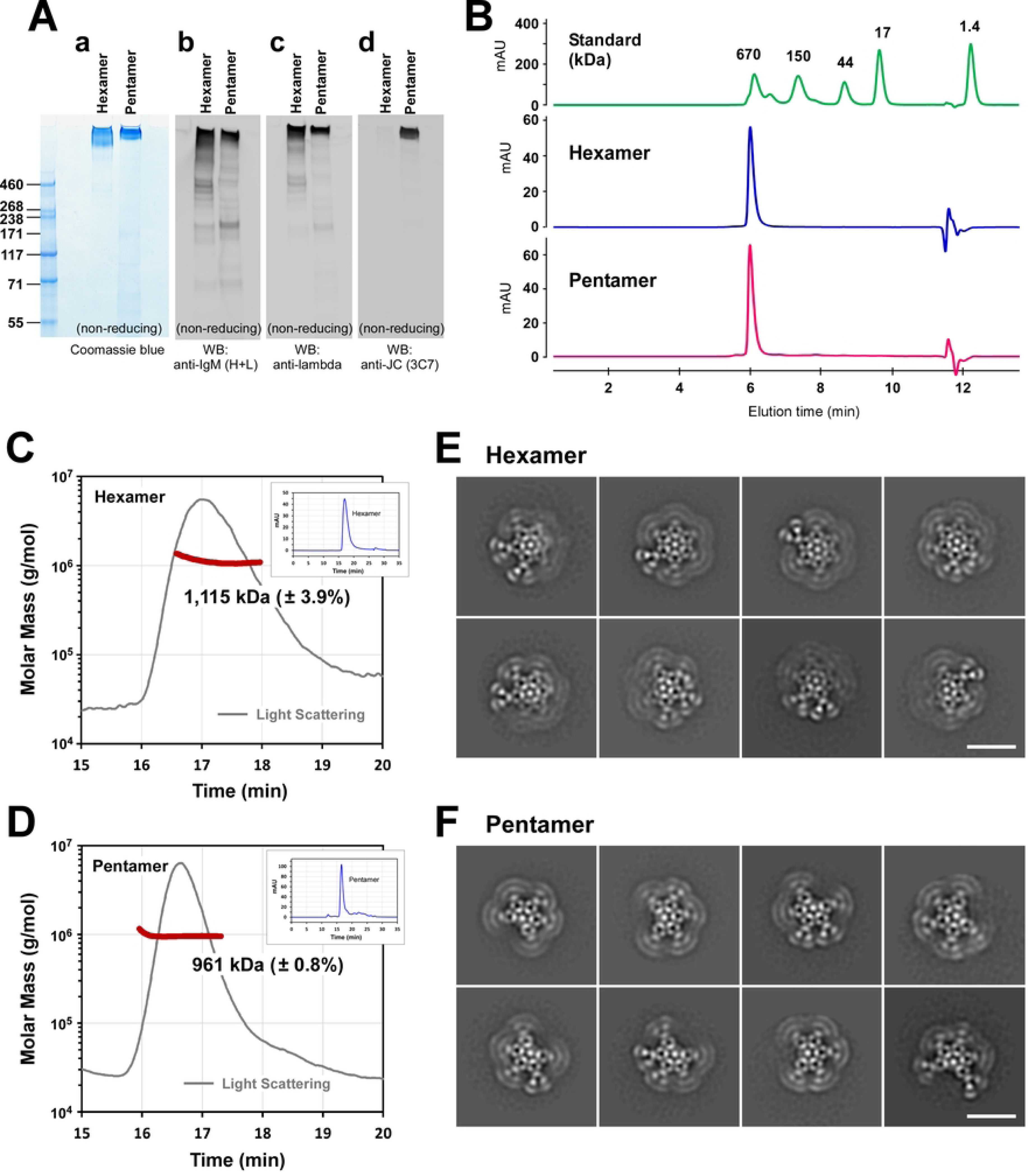
Characteristics of purified hexameric and pentameric IgM proteins. (A, a) Two-step purified hexameric and pentameric IgM preparations were resolved by SDS-PAGE under non-reducing conditions using a 3‒8% Tris-Acetate gradient gel followed by Coomassie blue staining. Sample loading was 2.5 μg per lane for both hexameric and pentameric IgMs. (A, b‒d) Identically run samples were analyzed by Western blotting using three different detection probes: (A, b) polyclonal anti-IgM (H+L), (A, c) polyclonal anti-λLC, and (A, d) monoclonal anti-JC antibody. (B) Overall protein purity and elution profile of two-step purified hexameric or pentameric IgMs were assessed by analytical SEC. About 5 μg of purified protein was injected for each IgM preparation. (C, D) Purified hexameric and pentameric IgM proteins (100 μg each) were analyzed by SEC-MALS. Light scattering and estimated molecular mass are plotted by solid gray line and bold brown line, respectively. The average molecular mass for (C) hexameric IgM peak and (D) pentameric IgM peak is stated in respective graphs. For calculation purposes, we arbitrarily used the mass of Man9GlcNac2 (= 1,865.64 Da) for the N-glycan because it represents one of the species exported out of the ER. Simultaneously collected SEC UV trace data are shown in the inset of C and D. (E, F) Negative-stain transmission electron micrographs of purified recombinant IgM. Eight representative 2D class averages for (E) hexameric IgM and (F) pentameric IgM are shown. Scale bar, 25 nm.

To test the effect of varying amounts of JC subunit on IgM product quality, we co-transfected an increasing amount of JC construct on top of the [μHC + λLC] pair while maintaining the total amount of transfecting DNAs constant (= 10 μg) and keeping the DNA ratio of μHC-to-λLC to be 1-to-1. In this scheme, the amount of DNA for the [μHC + λLC] pair decreases as the DNA for JC increases in a compensatory fashion (see Fig. 6 C D, top of the lanes). As expected, the amount of secreted [μHC + λLC] pair in culture media (Fig. 6C, lanes 1–7) and the expressed proteins in cell lysates (Fig. 6C, lanes 8–14) declined as the specific DNAs for the subunit pair decreased. By contrast, the secreted JC (Fig. 6D, lanes 1–7) and the JC protein level in cell lysates (Fig. 6D, lanes 8–14) increased as the relative amount of transfected JC DNA increased.

To assess the quality of secreted IgM products under the varied subunit chain ratio settings shown above, we analyzed the cell culture media under non-reducing settings to find a condition that promotes the incorporation of JC into the assembling polymeric IgM most effectively. In the range of DNA ratio tested, the lowest amount of JC DNA (2 μg) and the highest amount of [μHC + λLC] DNA pair (4 μg each) produced the greatest amount of JC-positive IgM product (Fig. 6 E–G, lane 2). Whenever the amount of JC exceeded the need for pentamer IgM formation, the excess free JC was released into the culture media in covalently linked polymers without being incorporated into any discrete assembly intermediates (Fig. 6G, lanes 2–6). Especially when the JC construct was expressed by itself, secreted JCs produced a smear-like Western blot that was broadly distributed over a wide range of molecular weight (Fig. 6G, lane 7). The fact that the smear was detected only under non-reducing conditions (compare Fig. 6, D and G, lane 7) indicated that secreted free JCs were extensively disulfide-linked, perhaps joined like concatemers using two or more of its eight reactive cysteine residues. While Cys-15 (the second Cys) and Cys-69 (the third Cys) are known to participate in the inter-chain disulfide bond formation in the context of pentameric IgM [77] and secretory IgA [78], whether they are responsible for this extensive concatemer formation is unknown. This type of redox-sensitive smearing in Western blotting was similar to the characteristics reported for cysteine-rich cytokine IL-31 [45]. Interestingly, this observation came at odds with the assertion made back in 1975 that JC can only be secreted into culture media in covalent association with polymeric IgM [79]. Importantly, however, we did not detect any extracellularly released assembly intermediates that were positive for JC (Fig. 6 F G). In other words, when JC was incorporated into assembling IgM, it was found only in the context of fully assembled IgM pentamers.

### 3.7. Production and characterization of J-chain-positive pentameric IgM

Because our secretion yield was sufficiently high, we continued to employ the same condition used above (see Fig. 6G, lane 2) without any additional optimization to produce the pentameric IgMs. Like the hexameric IgM species (see above), the fact that secreted pentameric SAM-6 IgM bound to a Protein L column served as a convenient initial affinity purification method despite the LC isotype being λ [46, 47]. From a batch of 300 mL harvested cell culture medium, the final purified yield was about 4 mg (see Materials and Methods).

SDS-PAGE under non-reducing conditions did not reliably discriminate the hexamers from the pentamers even when a lower % gradient gel was used to help resolve high molecular weight proteins (Fig. 7A, a). Likewise, Western blotting using polyclonal anti-IgM (H+L) or polyclonal anti-λLC could not differentiate the hexamers from the pentamers because both forms of IgM were detected equally by these two probes (Fig. 7A, b and c). In the Western blotting method, pentamers and hexamers were distinguishable only when these IgM products were probed with the monoclonal anti-JC antibody (Fig. 7A, d). Although the presence or absence of JC could differentiate the two different IgM preparations, the evidence was still missing as to whether the JC-positive IgMs were indeed pentamers and the JC-negative IgMs were hexamers. Similarly, the result did not rule out the possibility that the JC-positive IgM preparation was a heterogeneous mixture of pentamers and hexamers—because we do not have a method to detect the JC-negative IgM species selectively.

In another attempt to differentiate the two types of IgMs, we examined the purified IgM products by analytical SEC. As shown in Fig. 7B, both preparations showed a single main peak indicating high protein purity, but this method still did not distinguish the two types of IgMs based on the elution time or the shape of the elution peak. To look for a definitive analytical approach that differentiates the two forms of IgMs, we next carried out SEC-MALS to detect the molecular mass of protein complexes present in each purified IgM preparation. The average mass of IgMs detected in the putative hexamer peak was 1,115 kDa ± 3.9% (see Fig. 7C), and it was very close to the calculated mass of 1,135.2 kDa for a SAM-6 hexameric IgM (= 12 μHCs with 60 N-glycans + 12 λLCs). For the N-linked glycan, we arbitrarily used the mass of Man9GlcNac2 (= 1,865.64 Da) as it is one of the common N-glycan species exported out of the ER. Likewise, the average mass for the putative pentameric IgM protein peak was 961 kDa ± 0.8% (see Fig. 7D) and, again, it was very close to the calculated theoretical mass of 965.9 kDa for SAM-6 pentamer IgM (= 10 μHCs with 50 N-glycans + 10 λLCs + 1 JC with 1 N-glycan). From the standpoint of molecular mass, the IgMs produced without the JC consisted mainly of hexamers, while the IgMs produced with the JC were predominantly made of pentameric IgMs. To test if this conclusion is supported by an orthogonal method, we visualized both types of IgM by negative-stain transmission electron microscopy (EM) to obtain morphological evidence on IgM polymer status. To this end, a total of 88,125 hexamer particles and 52,657 pentamer particles were picked on cisTEM [48] and exported to RELION [49] for 2D classification analysis (data not shown). The hexameric form was indeed predominant in the purified hexameric IgM preparation, as expected from the MALS data. Eight representative micrographs shown in Fig. 7E are the class averages obtained from hexameric IgM preparation. The images indicated the features of the hexameric arrangement of six protomers. Interestingly, these images suggested the difficulty of visualizing all 12 Fab regions simultaneously on the same focal plane. Similarly, EM data showed that pentamers were predominant in the JC-positive pentameric IgM preparation. Eight representative class averages obtained from the pentameric IgM preparation are shown in Fig. 7F. Although the pentamers were predominant, we also obtained a few class averages showing a hexameric form (data not shown). Additionally, ring-like structures that did not conform to known IgM structures were obtained (data not shown). SAM-6 IgM pentamers’ morphology was similar to the recently reported structure of five protomer arrangements of mouse IgM-Fcμ [80] and human IgM-Fcμ [81–83]. These pentamers have a space gap where a JC replaces a single protomer, saving space for accommodating one AIM protein through a disulfide bond and charge-based interactions [80]. These morphological data strengthened that a suitable range of JC expression was all it took to shift the production of hexamers to pentamers. However, similar to what has been reported in homologous cellular models of plasma cells [17, 75, 76], a small fraction of hexameric IgM was still inevitably assembled by chance, even in the presence of abundant JC subunit during IgM biosynthesis in the ER lumen.

### 3.8. Without λLC, J-chain co-aggregates with μHC via Cys-575 and induces Russell body

The previous two sections revealed that IgM pentamer is readily assembled and secreted when three subunits [μHC + λLC + JC] were co-expressed in a permissible stoichiometry range. In this section, we mimicked the situations where either μHC or λLC expression level becomes insufficient and deviates far from the optimal subunit stoichiometry balance during the pentamer production. Specifically, we examined how JC’s intracellular behavior was altered if we dropped out the λLC or μHC subunit from the 3-chain transfection scheme.

When μHC was omitted from the 3-chain expression scheme, JC’s steady state distribution was not influenced by the remaining λLC subunit. Namely, λLC continued to distribute to the ER and Golgi (Fig. 8A, green), while the JC continued to localize in the ER without accumulating to the Golgi (Fig. 8A, red). In other words, their respective steady state dynamics remained the same as if they were expressed individually (see also Fig. 1E and Fig. 6B). The results suggested that λLC and JC did not interact, or at least their interaction was not significant enough to influence each other’s biosynthesis. As a result, λLC continued to be secreted as a mixture of monomers and covalent dimers (Fig. 8 H I, lane 1), while the JC was secreted in a covalently concatemerized format that shows up as a smear (Fig. 8 J K, lane 1). Even if the μHC expression was lost entirely during the pentamer IgM production, such an event can go unnoticed at the cellular level because it will not be manifested as prominent cell phenotype alternation.

**Figure 8.**
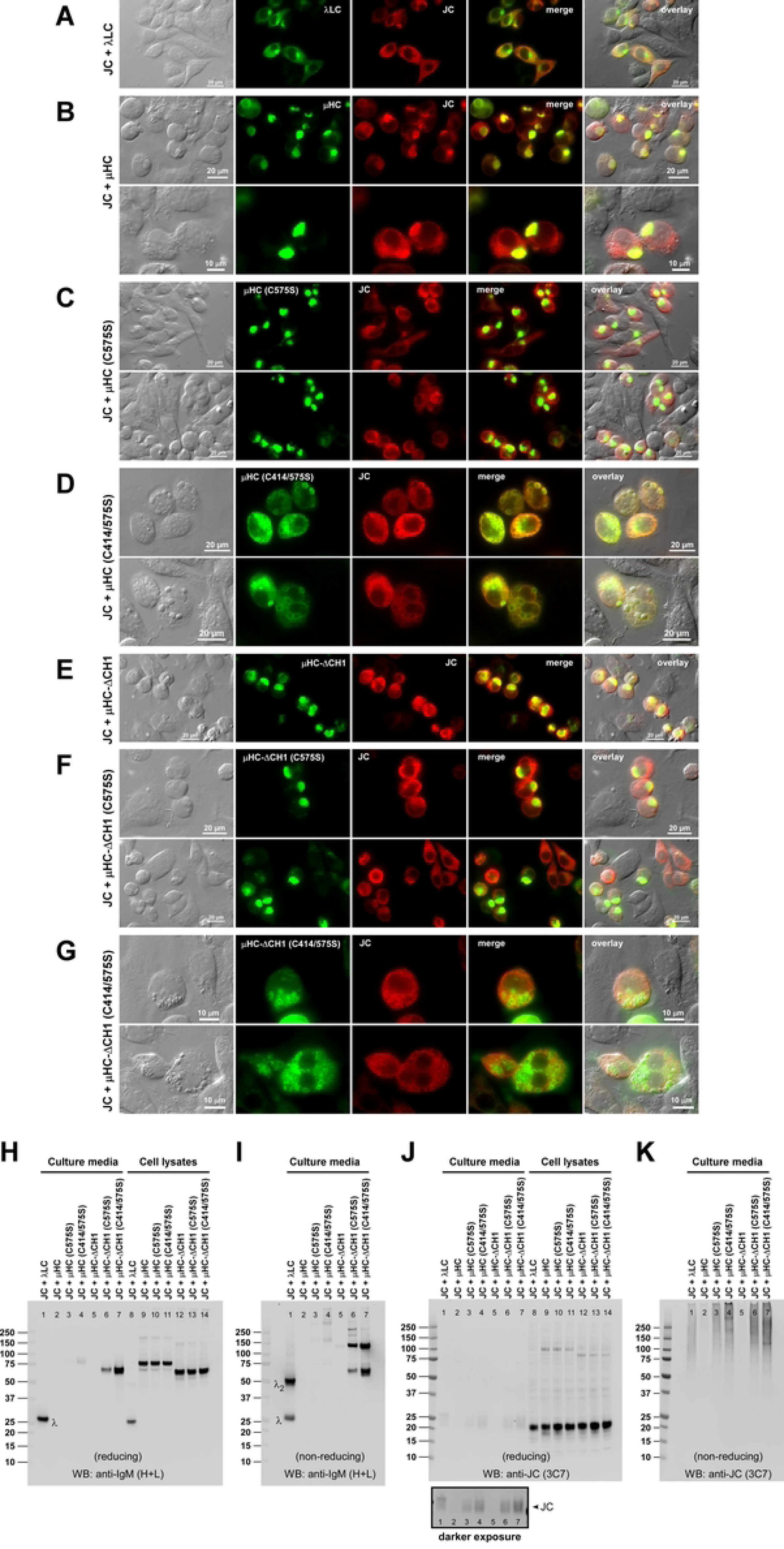
The intracellular distribution of the J-chain is markedly influenced by the characteristics of the co-expressed μHC subunit. Fluorescent micrographs of HEK293 cells co-transfected with JC and one of the following constructs: (A) λLC, (B) μHC, (C) μHC (C575S) mutant, (D) μHC (C414/575S) mutant, (E) μHC-ΔCH1 mutant, (F) μHC-ΔCH1 (C575S) mutant, or (G) μHC-ΔCH1 (C414/575S) mutant. A co-transfected construct pair is also shown on the left of each row. On day-3 post-transfection, cells were fixed, permeabilized, and co-stained with FITC-labeled anti-human λLC antibody (A) or FITC-labeled anti-human μHC antibody (B‒G) and monoclonal anti-human JC antibody followed by AlexaFluor594-conjugated secondary antibody (A‒G, shown in red). Green and red image fields were superimposed to create ‘merge’ views. DIC and ‘merge’ were superimposed to create ‘overlay’ views. (H, I) At day-7 post-transfection, cell culture media were harvested (lanes 1‒7), and cell lysate samples were prepared (lanes 8‒14) to run SDS-PAGE under reducing condition (panel H) or non-reducing condition (panel I) followed by Western blotting. Blotted membranes were probed with polyclonal anti-IgM (H+L) antibodies to detect both μHC and λLC subunits and assembly intermediates simultaneously. A co-transfected construct pair is shown at the top of each lane. (J, K) The same set of culture media and cell lysate samples were analyzed by Western blotting using anti-JC monoclonal antibody after protein samples were resolved by SDS-PAGE under (J) reducing or (K) non-reducing conditions. A longer exposed Western blot result is shown underneath the corresponding lanes in a black box for the cell culture media analyzed under reducing conditions (panel J, lanes 1‒7).

In an earlier section, the μHC subunit was shown to aggregate into Russell body by itself (see Fig. 1C and 3B). In the [μHC + JC] pair co-transfection setting, the μHC subunit still aggregated into Russell body even when the JC was co-expressed (Fig. 8B, green). On the other hand, JC no longer distributed to reticular ER as it usually would, but instead, it was co-aggregated with the μHC into Russell body (Fig. 8B, red and merge). Under this setting, not only was the μHC secretion blocked as expected (Fig. 8 H I, lane 2), but the JC secretion was also prevented almost completely (Fig. 8 J K, lane 2). These two proteins clearly interacted heavily to the point of co-aggregating into Russell body, which led to the retention of JC in the ER. This suggests that if λLC expression was lost during the pentamer IgM production process, such an event is readily noticeable because of the dramatic change in JC’s steady state subcellular distribution.

To dissect the mechanics behind the co-aggregation of μHC and JC into Russell body in the absence of λLC, we tested the mutant μHC bearing C575S or C414/575S mutation in the same experimental condition. While μHC (C575S) mutant continued to aggregate into the Russell body as before (Fig. 8C, green), the co-expressed JC no longer accumulated into the globular structure as judged from its lack of significant co-localization with the Russell body produced by the μHC (C575S) mutant (Fig. 8C, red and merge). In this co-expression setting, although μHC (C575S) remained secretion incompetent (Fig. 8 H I, lane 3), the JC secretion was restored to a normal level (Fig. 8 J K, lane 3). Expectedly, the penultimate Cys-575 of the μHC tailpiece interacted covalently with the JC, forcing the JC to co-aggregate with μHC into Russell body and making the JC non-secreting.

When the μHC (C414/575S) double mutant was expressed by itself, it induced Russell bodies extensively (see above, Fig. 4B, third row). However, when co-expressed with the JC subunit, the μHC (C414/575S) mutant now induced droplet-like inclusion bodies (Fig. 8D) instead of Russell bodies. The apparent droplet morphology was similar to the previously characterized cryoglobulin-like scFv-Fc-stp (stp, secretory tailpiece) that induced intra-ER protein droplet by liquid–liquid phase separation (LLPS) [84]. Immunostaining with anti-μHC or anti-JC antibodies could not stain the interior of the droplet inclusion bodies but only managed to stain their surface (Fig. 8D); again, similar to the reported case for scFv-Fc-stp [84]. The fact that the presence of JC altered the intra-ER behavior of μHC (C414/575S) indicated that these two proteins interacted in the ER lumen and influenced each other’s solution behavior, resulting in such a notable change in inclusion body type. Because Cys-414 and Cys-575 of μHC were mutated to Ser, the interactions were unlikely to be covalent. In fact, while μHC (C414/575S) remained largely non-secreting, the co-expressed JC retained secretion competence (Fig. 8 J K, lane 4), suggesting that their interactions are weak and transient.

We also performed the same JC co-expression experiments for the μHC-ΔCH1 mutant series. Regarding the intracellular protein distribution, μHC-ΔCH1 series behaved just like the full-length μHC variants did in the presence of JC. Namely, μHC-ΔCH1 continued to aggregate into Russell body (Fig. 8E, green) even when JC was co-expressed. The co-expressed JC, in turn, was drawn into the Russell body and co-aggregated with μHC-ΔCH1 (Fig. 8E, red and merge). Similarly, μHC-ΔCH1 (C575S) mutant continued to induce Russell body (Fig. 8F, green) in the presence of JC, but the co-aggregation of JC into Russell body was prevented (Fig. 8F, red and merge) because the culprit Cys-575 was mutated to Ser. In the presence of JC, the double mutant construct μHC-ΔCH1 (C414/575S) no longer distributed to reticular ER as it usually would (see above, Fig. 5B). Instead, the co-expression of JC changed the intracellular dynamics of μHC-ΔCH1 (C414/575S) protein into inducing droplet-like inclusion bodies (Fig. 8G). The induction of droplet inclusion body through the interaction with JC was therefore a CH1-independent event.

The secretion pattern of μHC-ΔCH1, μHC-ΔCH1 (C575S), and μHC-ΔCH1 (C414/575S) in the presence of JC was identical to when these constructs were expressed by themselves (compare Fig. 8H, lanes 5–7 and Fig. 4C, lanes 10–12). Likewise, the protein quality of secreted μHC-ΔCH1, μHC-ΔCH1 (C575S), and μHC-ΔCH1 (C414/575S) was not influenced by the presence of JC either (Fig. 8I, lanes 5–7). For example, whenever JC was drawn into the Russell body via μHC’s Cys-575, JC secretion was blocked (Fig. 8K, lane 5); but if the JC was excluded from the Russell body by C575S mutation, the JC secretion took place as usual (Fig. 8K, lane 6). Similarly, when μHC-ΔCH1 (C414/575S) formed a similar droplet inclusion body in the presence of co-expressed JC, only the JC was secreted (Fig. 8K, lane 7).

### 3.9. Roles of CH1 domain and Cys-414/Cys-575 residues in pentameric IgM assembly and secretion

Secretory production of pentameric IgM was surprisingly straightforward. It only took a co-expression of three comprising subunit chains at a permitted stoichiometry range. To appreciate the importance of μHC-encoded key structural determinants in pentameric IgM formation and secretion, we tested the five μHC mutants shown in Fig. 4A and looked for which assembly step is affected in the 3-chain co-transfection experiments.

Before analyzing the secreted product quality, the expression of all three proteins was validated by Western blotting on cell lysate samples, as shown in Fig. 9A, lanes 1–6. The secretion outputs of individual protein components varied depending on each mutation’s specific effect on the pentameric IgM assembly (Fig. 9A, lanes 7–12). Of note, the JC secretion was detectable in the culture media only when co-expressed with the parental [μHC + λLC] pair (Fig. 9A, bottom panel, lane 7). This was, in fact, the only condition that allowed the productive incorporation of JC into the assembling pentameric IgMs (see Fig 9 B–D, lane 1, red arrowhead). In all other settings, the JC was not detectable in the culture media (Fig. 9A, bottom panel, lanes 8–12), and, as mentioned above, the JC was not incorporated into any assembly intermediates (Fig. 9D, lanes 2–6). Similarly, overall protein secretion was poor when μHC-ΔCH1 mutant was co-expressed with λLC and JC (Fig. 9A, lane 10). This was because all three proteins extensively co-aggregated into Russell body (Fig. 9F, first row).

Regarding protein quality, covalently assembled pentameric IgM was formed only when the parental μHC, λLC, and JC subunits were co-expressed (Fig. 9 B–D; lane 1, red arrowhead). In other words, the JC subunit was covalently incorporated into a higher-order protein complex only when co-expressed with the parental [μHC and λLC] pair (Fig. 9D, lane 1). The blockade of covalent assembly by C575S or C414/575S mutation effectively disrupted the covalent association between the μHC homodimers and between JCs and the two μHC homodimers. As a result, these mutations prevented the formation of covalent IgM pentamers (Fig. 9 B C, lanes 2– 3) but instead led to the secretion of ∼200 kDa μ2λ2 protomers and ∼100 kDa μλ half-molecules as main products (Fig. 9 B C, lanes 2–3). By contrast, in the ΔCH1 mutant series, the λLC–μHC covalent assembly was abolished, as shown by the lack of λLC-containing protein complexes larger than λLC dimers (Fig. 9C, lanes 4–6). Although μHC-ΔCH1 mutant was non-secreting (Fig. 9B, lane 4), the μHC-ΔCH1 became secretable as a mixture of monomers and homodimers when C575S or C414/575S mutation was introduced (Fig. 9B, lanes 5–6; black and blue arrowheads, respectively). These mutant HC monomers and homodimers are devoid of λLC subunit, and the corresponding protein bands were not detected when probed with anti-λLC antibody (Fig. 9C, lanes 4‒6). Lastly, regardless of the μHC mutations, the excess free λLC was consistently secreted as a mixture of monomers and disulfide-bonded dimers under all conditions tested (Fig. 9 B C).

Co-expression of 3 parental subunit chains resulted in the distribution of [μHC + λLC] to the ER and the characteristic cytoplasmic puncta (Fig. 9E, first row, green), while the JC remained in the ER (Fig. 9E, first row, red). By contrast, when the μHC carried the C575S or C414/575S mutation, the μHC and λLC distributed to the ER and Golgi without showing the punctate staining (Fig. 9E, second and third rows), while the JC remained distributed to the ER. Although μHC-ΔCH1’s aggregation into Russell body was prevented by the presence of λLC (see above, Fig. 5B, first row), the additional expression of JC made these subunit chains heavily aggregated into Russell body again (Fig. 9F, first row). The same Russell body phenotype in similar settings was first observed in 1991 [68, 69], and Sitia’s group proposed that the ΔCH1 mutation was the leading cause of Russell body formation without realizing that the presence of JC was another requirement in this particular case of Russell body phenotype induction. Because Russell body formation was suppressed by introducing the C575S mutation to the μHC-ΔCH1 subunit (Fig. 9F, second row, green), it was clear that covalent association between JC and μHC-ΔCH1 via Cys-575 was the underlying cause of the extensive Russel body formation in the presence of λLC. By contrast, in the 3-chain expression involving ΔCH1 (C575S) or ΔCH1 (C414/575S) subunit, both μHC mutants and λLC showed the ER and Golgi distribution (Fig. 9F, second and third rows, green), while JC was consistently in the ER as expected (red).

Although expected, this section showed that in order to produce a covalently assembled pentameric IgM species, not only does the CH1 domain need to be intact, but the Cys-414 and Cys-575 residues must also remain available for intricate inter-chain disulfide bond formation among the comprising three subunit chains.

### 3.10. Covalent association between μHC and λLC is not required for IgM-like product polymerization and secretion

To characterize the role of covalent interaction between μHC and λLC in IgM biosynthesis, we disabled the λLC’s capacity to form inter-chain disulfide bridge by truncating the C-terminal two amino acids (i.e., Cys-213 and Ser-214), of which the penultimate Cys-213 is essential for bridging the λLC to μHC by a disulfide bond. We designated such mutant as λLC-ΔCS (see Suppl. 1A). A similarly engineered λLC-ΔCS mutant of different clonal origin previously demonstrated the effectiveness of the ΔCS mutation in abolishing λLC’s inter-chain disulfide bond formation [56]. The ΔCS version of SAM-6 λLC also showed negligible ability to form disulfide-linked homodimers (see Suppl. 1 B C, lane 6) without affecting the total secretion level (Suppl. 1 B C, lanes 3 and 4) or the steady state subcellular distribution during protein synthesis (Suppl. 1 D E). By using this SAM-6 λLC-ΔCS mutant, we aimed to demonstrate the criticality of covalent linkage between μHC and λLC in IgM biosynthesis, while keeping the CH1 domain of μHC completely intact.

As a prerequisite, the protein expression level of all three constructs was first validated by Western blotting on cell lysates (Fig. 10A; top, second, third panels; lanes 1–4). Likewise, co-transfected subunit chains were all found secreted and were detectable in the culture media when similarly analyzed under reducing conditions (Fig. 10A; top, second, third panels; lane 5– 8, black arrowheads). At this point, however, we were already puzzled by the observation that the μHC subunit was abundantly detected in the culture media when co-transfected with λLC-ΔCS (Fig. 10A; top panel, lanes 6 and 8) ―because we presumed that μHC would not exit the ER unless the μHC subunit assembles covalently with λLC via a disulfide bond. Nonetheless, when it comes to the product secretion level, both 2-chain and 3-chain transfected products gave a comparable secretory output regardless of whether full-length λLC or its ΔCS mutant was used (Fig 10A, compare lanes 5 and 6 for 2-chain setting; lanes 7 and 8 for 3-chain setting).

**Figure 9.**
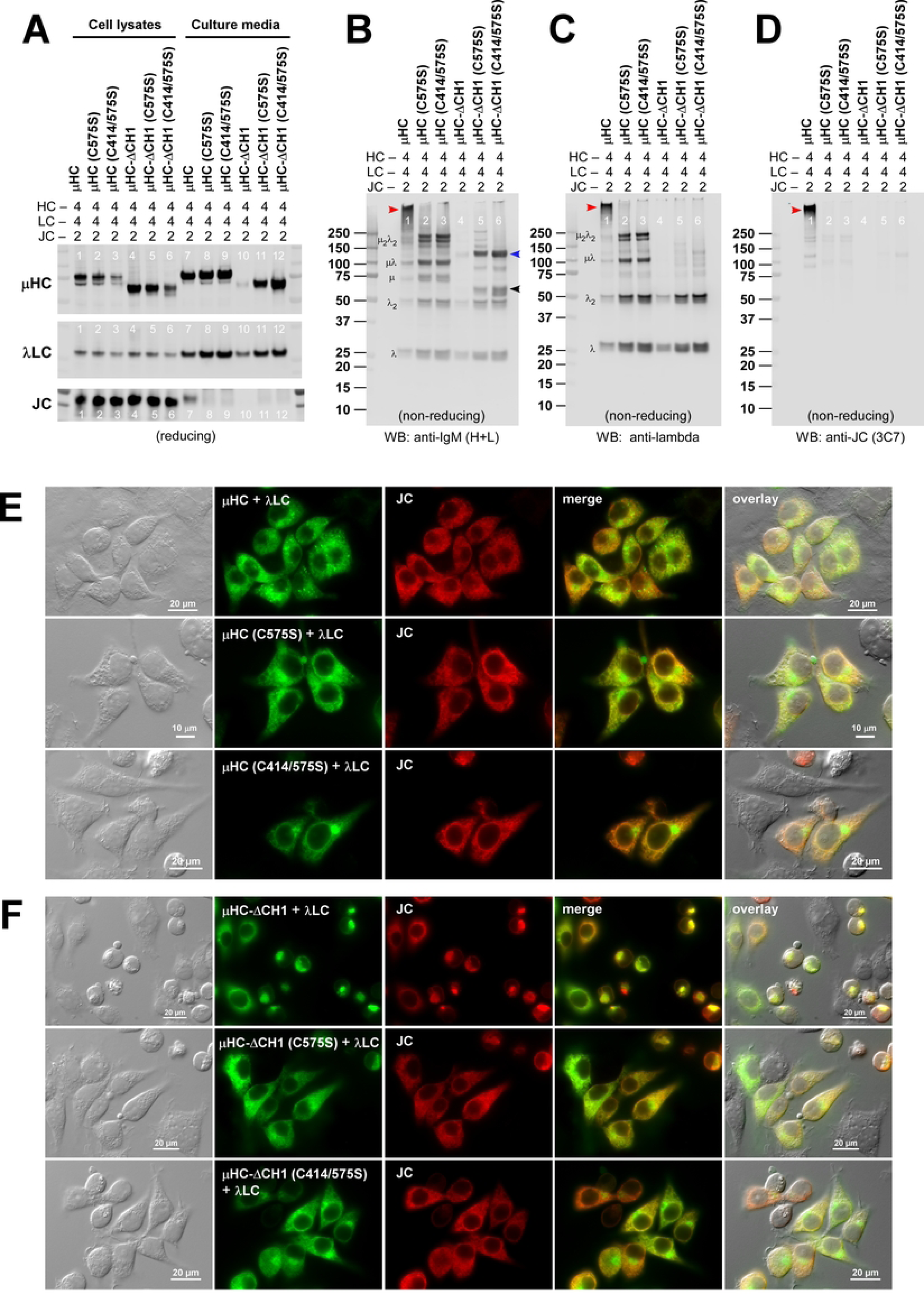
Differential effects of ΔCH1, C575S, and C414/575S mutations on pentameric IgM assembly and secretion. (A‒D) The effect of ΔCH1, C575S, or C414/575S mutation on IgM pentamer formation was tested. As before, three subunit chains were co-transfected to produce pentameric IgM species at the ratio of μHC : λLC : JC = 4 : 4 : 2. (A) At day-7 post-transfection, cell lysates (lanes 1‒6) and cell culture media samples (lanes 7‒12) were prepared and resolved by SDS-PAGE under reducing conditions followed by Western blotting. Blotted membranes were probed with polyclonal anti-IgM (H+L) antibodies to detect μHC and its mutants (top panel) and λLC subunit (middle panel) as well as with monoclonal anti-JC antibody (bottom panel). (B‒D) Day-7 cell culture media were also analyzed by Western blotting after resolving the proteins under non-reducing conditions. Blotted membranes were probed with (B) polyclonal anti-IgM (H+L) to simultaneously detect IgM and various assembly intermediates, (C) polyclonal anti-λLC to detect IgM and a subset of assembly intermediates containing λLC subunit, or (D) monoclonal anti-JC antibody to detect pentameric IgM or assembly intermediates containing JC subunit. Identifiable assembly intermediates are labeled next to the corresponding protein bands in panels B and C. The protein band corresponding to the assembled pentameric IgM is pointed by a red arrowhead in lane 1 of panels B‒D. Secreted μHC-ΔCH1 (C414/575S) mutant monomers and dimers are pointed by black and blue arrowheads in panel B (lanes 6), respectively. (E) Fluorescent micrographs of HEK293 cells co-transfected with the following set of 3 constructs: (top row) [JC + λLC + μHC] combination; (middle row) [JC + λLC + μHC (C575S)] combination; (bottom row) [JC + λLC + μHC (C414/575S)] combination. As before, suspension-cultured transfected cells were seeded onto poly-D-lysine coated coverslips on day-2 post-transfection and then statically grown for 24 hr. On day-3 post-transfection, cells were fixed, permeabilized, and co-stained with a 1-to-1 mix of FITC-labeled anti-human μHC antibody and FITC-labeled anti-human λLC antibody to stain both subunits simultaneously (shown in green) and monoclonal anti-JC antibody followed by AlexaFluor594-conjugated secondary antibody (shown in red). Green and red image fields were superimposed to create ‘merge’ views. DIC and ‘merge’ were superimposed to create ‘overlay’ views. (F) Fluorescent micrographs of HEK293 cells co-transfected with the following set of 3 constructs: (top row) [JC + λLC + μHC-ΔCH1] combination; (middle row) [JC + λLC + μHC-ΔCH1 (C575S)] combination; (bottom row) [JC + λLC + μHC-ΔCH1 (C414/575S)] combination. Suspension-cultured transfected cells were seeded, fixed, and immunostained as above. Captured images were processed as described above.

**Figure 10.**
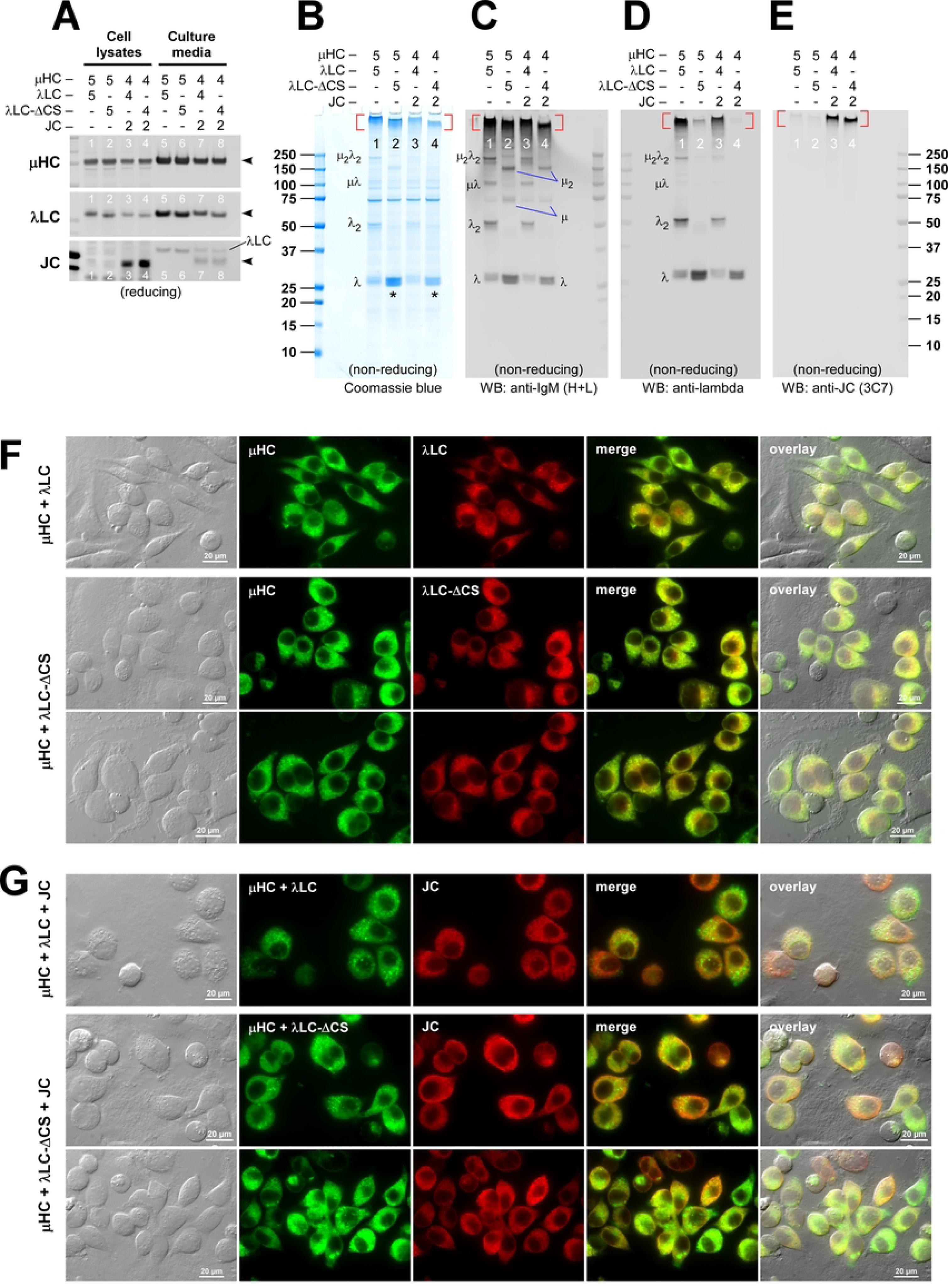
Polymeric IgM-like product is assembled and secreted without an inter-chain disulfide linkage between μHC and λLC. (A) Effect of LC-ΔCS mutation on protein expression and secretion was tested in a 2-chain co-expression scheme (lanes 1‒2 and 5‒6) and a 3-chain co-expression setting (lanes 3‒4 and 7‒8). Subunit chains were co-transfected at the DNA ratio indicated at the top of each lane. At day-7 post-transfection, cell lysates (lanes 1‒4) and cell culture media samples (lanes 5‒8) were prepared and resolved by SDS-PAGE under reducing conditions followed by Western blotting. Blotted membranes were probed with polyclonal anti-IgM (H+L) antibodies to detect μHC (top panel) and λLC or λLC-ΔCS (second panel) or with monoclonal anti-JC antibody (third panel). (B‒E) Day-7 cell culture media were resolved by SDS-PAGE under non-reducing conditions, and Coomassie blue stained in panel B and analyzed by Western blotting in panels C‒E. Blotted membranes were probed with (C) polyclonal anti-IgM (H+L) to simultaneously detect IgM and various assembly intermediates composed of μHC or λLC or both; (D) polyclonal anti-λLC to selectively detect IgM and subsets of assembly intermediates containing λLC or λLC-ΔCS; or (E) monoclonal anti-JC antibody to detect pentameric IgM or any assembly intermediates containing the JC subunit. The hexameric or pentameric IgM proteins are indicated by red brackets in panels B‒E. Protein bands corresponding to λLC monomers and dimers (or λLC-ΔCS monomers) and assembly intermediates are labeled on individual gels shown in panels B‒D. (F) Fluorescent micrographs of HEK293 cells co-transfected with (top row) [μHC + λLC] pair, (second and third rows) [μHC + λLC-ΔCS] pair. (G) Likewise, (first row) [μHC + λLC + JC] combination, (second and third rows) [μHC + λLC-ΔCS + JC] combination. On day-3 post-transfection, cells were fixed, permeabilized, and co-stained with (F) FITC-labeled anti-human μHC antibody and Texas Red-labeled anti-human λLC antibody or (G) a 1-to-1 mix of FITC-labeled anti-human μHC and FITC-labeled anti-human λLC antibodies (shown in green) and monoclonal anti-JC antibody followed by AlexaFluor594-conjugated secondary antibody (shown in red). Green and red image fields were superimposed to create ‘merge’ views. DIC and ‘merge’ were superimposed to create ‘overlay’ views.

To delineate the quality of secretion-competent products made with the λLC-ΔCS mutant, we examined the secreted products under non-reducing conditions to preserve pre-existing disulfide bonds (Fig. 10, B‒E). As repeatedly shown above, both hexameric and pentameric IgMs were secreted abundantly when the designated 2-chain and 3-chain parent constructs were co-transfected, respectively, at the indicated stoichiometry (Fig. 10B, lane 1 for hexamer; lane 3 for pentamer, see red bracket area). Interestingly, the formation and secretion of high molecular weight IgM-like product were not impacted by the use of λLC-ΔCS mutant in the 2-chain and 3-chain settings (Fig. 10B, compare lanes 1 and 2 for hexamer; lanes 3 and 4 for pentamer)—indicating that IgM biosynthesis took place at a similar rate regardless of the λLC was full-length or ΔCS mutant. However, the apparent gel mobility of hexameric IgM was detectably different depending on when full-length λLC or λLC-ΔCS was used (Fig. 10B, compare lanes 1 and 2). Likewise, the apparent size of secreted pentameric IgM products was clearly different depending on when the full-length λLC or ΔCS mutant was co-transfected (Fig. 10B, compare lanes 3 and 4). Namely, the size of IgM-like products expressed with the λLC-ΔCS subunit was consistently smaller than the bona fide IgMs expressed with the full-length λLC. In addition, the amount of detectable free λLC monomer was consistently more abundant whenever the ΔCS version λLC was used (Fig. 10B, compare lanes 1 and 2; lanes 3 and 4, marked by asterisks), thereby hinting at a possibility that the association of λLC-ΔCS mutant with the IgM-like polymer backbone was labile in the presence of SDS. The simplest way to explain such product size difference in SDS-PAGE was that (1) λLC-ΔCS associated with the μHC via non-covalent interactions, and (2) the non-covalently associated λLC-ΔCS readily dissociated from the polymeric μHC backbone when subjected to SDS-PAGE. This observation, in turn, indicated that (3) such non-covalent interactions between λLC-ΔCS and μHC were sufficient to hold together an IgM-like product at physiological conditions in the ER lumen and during secretory trafficking, and even after secreted into extracellular space. (4) It also suggested that when λLC-ΔCS associated with polymeric μHC backbone, such “quasi-IgM” behaved indistinguishably from bona fide IgM during biosynthesis, met the ER quality control criteria and was able to negotiate abundant secretion. (5) During SDS-PAGE analysis, however, the λLC-ΔCS dissociated and migrated separately from the rest of the polymeric μHC backbone. (6) As such, we detected a disproportionately more abundant free monomeric LC subunit whenever λLC-ΔCS was used.

To test if such interpretations are experimentally supported, we performed a set of Western blotting using the probes against different components of IgM and assembly intermediates (Fig. 10 C‒E). Probing with polyclonal anti-IgM (H+L) antibody detected both bona fide polymeric IgMs (Fig. 10C, lanes 1 and 3) and quasi-IgMs at comparable signal intensity (Fig. 10C, lanes 2 and 4) which was in good agreement with the results of Coomassie blue stained gels (Fig. 10B). Notably, in Fig. 10C, the assembly intermediates in lanes 2 and 4 were consistently ∼50 kDa or ∼25 kDa smaller than those intermediate species found in lanes 1 and 3; thereby clearly indicated that two or one non-covalently associated λLC-ΔCS subunit(s) dissociated from these assembly intermediates during SDS-PAGE (Fig. 10C, compare lanes 1 and 2; lanes 3 and 4). When the same samples were probed with polyclonal anti-λLC antibodies, the quasi-IgM polymers were indeed found devoid of LC subunit (Fig. 10D, compare lanes 1 and 2; lanes 3 and 4). Although a residual amount of λLC-ΔCS was detected in quasi-IgM products (Fig. 10D, lane 2), this was likely caused by an opportunistic off-pathway disulfide bonding via non-canonical cysteine connectivity, as reported previously for other class of immunoglobulin [85]. This residual opportunistic incorporation of LC-ΔCS was a reproducible event (see Fig. 13C, lanes 2 and 6). Lastly, based on the fact that JC was incorporated into the quasi-IgM pentamers at a comparable level as the bona fide IgM pentamers (Fig. 10E, compare lanes 3 and 4), it was clear that covalent incorporation of JC into polymerizing μHCs was independent of the mode of λLC‒μHC associations (i.e., covalent or non-covalent).

We next examined the steady state subcellular distribution of participating subunit chains during the 2-chain and 3-chain transfection settings to see if subunit localization was altered when the association between λLC and μHC was only mediated by non-covalent forces. Firstly, co-expression of λLC-ΔCS effectively suppressed the formation of μHC-induced Russell body (Fig. 10F, second and third rows). This suggested that non-covalent interactions between the λLC-ΔCS mutant and the full-length μHC were sufficient to support IgM assembly by repressing the aggregation of μHC into Russell body. Thus, the morphological evidence agreed very well with quasi-IgM hexamer’s seemingly normal assembly and secretion. Secondly, the [μHC + λLC-ΔCS] pair distributed to the ER and cytoplasmic punctata during biosynthesis, and the distribution pattern was indistinguishable from the biosynthesis of bona fide IgM hexamer (Fig. 10F, first row). Again, the results agreed with the comparable secretion output between bona fide hexamer IgM and quasi-IgM hexamer. Thirdly, even in the presence of JC to produce pentameric IgMs, the ΔCS mutation did not produce noticeable changes in steady state subcellular distribution of [μHC + λLC-ΔCS] pair and JC subunit (Fig. 10G, second and third row). The subunit distribution pattern was again indistinguishable from that of bona fide IgM pentamers (Fig. 10G, first row).

Surprisingly enough, the deletion of C-terminal Cys-Ser residues from the λLC was not detrimental to the overall biosynthetic processes of IgM assembly and secretion output despite the absence of covalent linkage between λLC-ΔCS and μHC. As a result, pentameric and hexameric forms of IgM-like products (or quasi-IgMs) were generated and secreted at an equivalent rate to the bona fide IgM products. In other words, overall biosynthesis of IgM in the ER lumen and secretory pathway trafficking proceeded expectedly even without the covalent linkage between λLC and μHC.

### 3.11. The inter-chain disulfide bond between μHC and λLC is ablated by introducing a minimally invasive C137S point mutation

Dispensability of inter-chain covalent interaction between μHC and λLC was unexpected because the earlier results of μHC-ΔCH1 mutant (see Fig. 3) and generally accepted knowledge [68] appeared to endorse the critical role of μHC–λLC covalent assembly in IgM product formation and secretion. In retrospect, however, deletion of the entire CH1 domain might have been too invasive to retain intricate non-covalent forces that were also at play in holding the μHC and λLC subunits together. To reproduce the effect of ΔCH1 in a much less invasive way, we introduced a C137S point mutation to μHC in order to prevent the disulfide bond formation with the Cys-213 residue of λLC. Because this will maintain the integrity of the CH1 domain, we expected this would preserve exquisite non-covalent interactions between the CH1 and CL as much as possible. Using this μHC (C137S) mutant subunit (see Suppl. 2A, second row), we reciprocally evaluated the dispensability of inter-chain covalent disulfide bonding between μHC and λLC during IgM biosynthesis.

When expressed by itself, μHC (C137S) mutant subunit showed high aggregation propensity and induced Russell body to a similar degree as the parental μHC (see Suppl. 2D). Likewise, μHC (C137S) mutant subunit was not able to be secreted to the culture media (see Suppl. 2 B C). The C137S mutation by itself did not alter the basic characteristics of μHC. In fact, when μHC (C137S) was co-expressed with JC, μHC (C137S) and JC co-aggregated into Russell body owing to the intact Cys-575 (Suppl. 3D, second row), and the secretion of JC was prevented as a result (Suppl. 3 A‒C, lanes 5 and 8).

To assess the effect of C137S mutation on IgM product formation, the μHC (C137S) mutant construct was co-expressed with the parental λLC or its disabled λLC-ΔCS counterpart in the absence of JC to generate hexamers or in the presence of JC to produce pentamers. As before, the protein expression level of each transfected construct was validated using the cell lysates by Western blotting to ensure that designated proteins were expressed at equivalent amounts under different construct combinations (Suppl. 4C; lanes 1‒4 for 2-chain; lanes 5‒8 for 3-chain). We also verified the secretion of each transfected subunit as part of the secreted protein complex (Suppl. 4 A B, lanes 1‒4 for 2-chain; lanes 5‒8 for 3-chain). In 2-chain co-expression settings, the secreted amount of μHC and λLC was comparable regardless of whether the μHC had the C137S mutation or not and whether λLC or λLC-ΔCS was used in any combinations (Suppl. 4 A B, lanes 1‒4). Likewise, in a 3-chain transfection scheme, a comparable amount of individual subunit chains was detected in the culture media as part of the secreted protein complex regardless of whether parental subunit or mutant subunit was used (Suppl. 4 A B, lanes 5‒8). When the secreted products were analyzed under reducing conditions, the effects of μHC (C137S) mutation were unnoticeable regarding protein secretion.

To define the effect of μHC (C137S) mutation on the covalent polymerization of IgM, we evaluated the secreted products under non-reducing conditions (Fig. 11, A‒C). Both in 2-chain (Fig. 11A, lanes 1‒4) and in 3-chain (Fig. 11A, lanes 5‒8) expression settings, the secreted amount of high molecular weight IgM-like species was equivalent to the parental IgM even when the parental μHC was substituted with μHC (C137S) mutant subunit (compare lanes 1 and 2 for 2-chain; lanes 5 and 6 for 3-chain). Moreover, the amount of secreted IgM-like product was comparable even when μHC (C137S) mutant was co-expressed with λLC-ΔCS mutant both in 2-chain (Fig. 11A, lane 4) and 3-chain (Fig. 11A, lane 8) settings. These results suggested, as before, that IgM-like covalent polymer assembly and secretion took place at a similar rate regardless of whether the parental μHC and λLC subunits were used or if μHC (C137S) and λLC-ΔCS mutants were used in any HC‒LC combinations. In other words, covalent pairing of μHC and λLC by disulfide bridge was not required for the IgM-like protein assembly, ER quality control steps, and product secretion.

**Figure 11.**
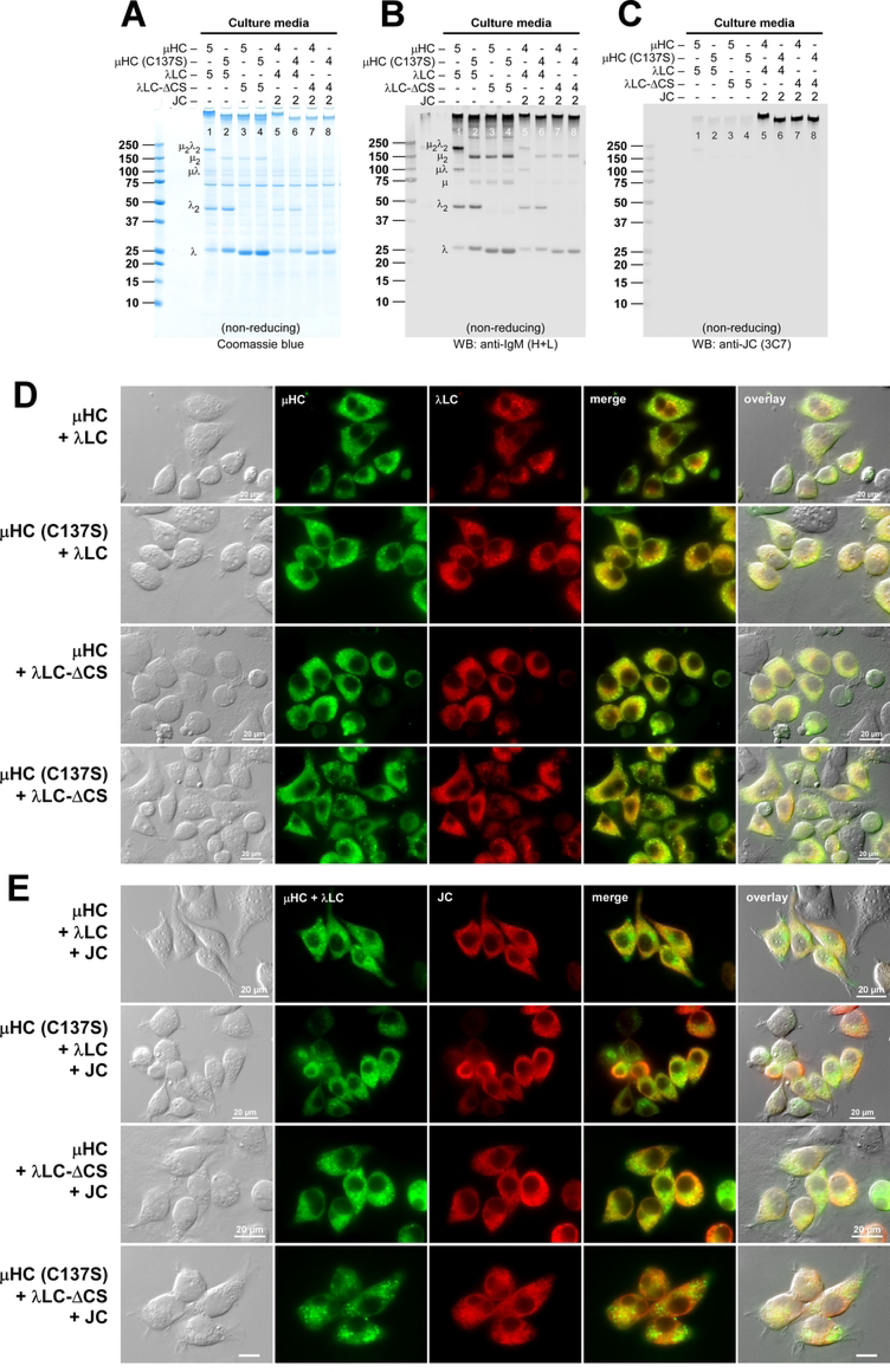
Polymeric IgM-like product is assembled and secreted independent of the covalent association between μHC and λLC. (A‒C) The effect of μHC (C137S) mutant subunit on covalent polymeric IgM assembly and secretion was assessed in a 2-chain co-expression scheme (lanes 1‒4) and a 3-chain co-expression setting (lanes 5‒8). Parental subunit chains and mutants were co-transfected at the DNA ratio indicated at the top of each lane. Day-7 cell culture media were resolved by SDS-PAGE under non-reducing conditions and Coomassie blue stained (A) and analyzed by Western blotting (B, C). Blotted membranes were probed with (B) polyclonal anti-IgM (H+L) and (C) monoclonal anti-JC antibodies. Protein bands corresponding to λLC monomers and dimers, as well as assembly intermediates, are labeled in panels A and B. (D, E) Fluorescent micrographs of HEK293 cells co-transfected with a combination of parental or mutant subunit chains that are intended to abolish inter-chain disulfide bond formation between μHC and λLC. Cells were transfected with a 2-chain construct set (D) and 3-chain construct set (E) to generate hexameric and pentameric IgM-like products, respectively. The transfected subunit combination is shown on the left of each row. On day-3 post-transfection, cells were fixed, permeabilized, and co-stained with (D) FITC-labeled anti-human μHC antibody and Texas Red-labeled anti-human λLC antibody or (E) a 1-to-1 mix of FITC-labeled anti-human μHC and FITC-labeled anti-human λLC antibodies (shown in green) and monoclonal anti-JC antibody followed by AlexaFluor594-conjugated secondary antibody (shown in red). Green and red image fields were superimposed to create ‘merge’ views. DIC and ‘merge’ were superimposed to create ‘overlay’ views.

Judging from the gel mobility differences, the size of hexameric IgM-like products made with the [μHC (C137S) + λLC] pair was smaller than the parental hexameric IgM (Fig. 11A, compare lanes 1 and 2). Importantly, the [μHC (C137S) + λLC] product size was indistinguishable from another hexameric IgM-like product assembled by [μHC + λLC-ΔCS] pair (compare lanes 2 and 3) or a product generated with [μHC (C137S) + λLC-ΔCS] subunit pair where the inter-chain disulfide-forming Cys residues were ablated on both subunits simultaneously (Fig 11A, compare lanes 2 and 4). Likewise, the apparent gel mobility of secreted pentameric IgM-like products was clearly smaller than the bona fide pentameric IgM when μHC (C137S) or λLC-ΔCS or both chains were used to replace the corresponding parental subunits (Fig. 11A, compare lanes 5‒8). These results all underscored the previous section’s finding that, in the presence of SDS, the association of λLC to the polymeric IgM-like backbone became labile because the inter-chain disulfide bond formation was abrogated. This conclusion was also indirectly supported by the observation that detectable free λLC monomer was consistently more abundant whenever the disulfide-forming Cys residues were ablated singly or doubly (Fig. 11A, compare lane 1 to lanes 2‒4). Again, the simplest way to explain this observation was that because of the μHC (C137S) or λLC-ΔCS mutation, μHC and λLC subunits were held together only by non-covalent interactions, and the non-covalently associated λLCs dissociated from the polymeric IgM-like backbone when subjected to SDS-PAGE. Conversely, non-covalent interactions between λLC and μHC were sufficiently strong enough to hold the subunits together during ER quality control processes and secretory pathway trafficking and even after secretion.

Western blotting using the probes against different constituents of the IgM and assembly intermediates further supported this conclusion (Fig. 11 B C). Probing with polyclonal anti-IgM (H+L) antibodies comparably detected both bona fide polymeric IgMs (Fig. 11B, lanes 1 and 5) and the IgM-like products (Fig. 11B, lanes 2‒4 for hexamers and lanes 6‒8 for pentamers). Expectedly, covalent λLC dimers were generated only when the parental λLC was used (Fig. 11B, lanes 1 and 2; lanes 5 and 6), whereas such dimer formation was suppressed when λLC-ΔCS was used (Fig. 11B, lanes 3 and 4; lanes 7 and 8). The major assembly intermediates for the bona fide IgMs were ∼200 kDa μ2λ2 and ∼100 kDa μλ for both hexamers and pentamers (Fig. 11B, lanes 1 and 5), whereas the detectable assembly intermediates for the IgM-like products were ∼150 kDa μ2 and ∼75 kDa μ species where two and one λLC subunit had dropped off from the μ2λ2 and μλ intermediates, respectively (Fig. 11B, lanes 2‒4; lanes 6‒8). Clearly, the inter-chain disulfide bridge between μHC and λLC was abolished in these IgM-like products and, as a result, the λLCs dissociated from the μHC during SDS-PAGE. Importantly, all three IgM-like products shared the same assembly intermediates, namely, ∼150 kDa μ2 and ∼75 kDa μ (Fig. 11B, lanes 2‒4; lanes 6‒8). Likewise, because non-covalently associated λLC and λLC-ΔCS dissociated from the IgM-like polymer backbone during SDS-PAGE, the detectable free LC subunit was again disproportionately more abundant in lanes 2‒4 and 6‒8. As before, JC was incorporated normally into the IgM-like pentamers at a comparable level as the bona fide IgM pentamers (Fig. 11C, lanes 5‒8), thereby showing that covalent incorporation of JC into the polymerizing μHC via Cys-575 was independent of the inter-chain disulfide bridge formation between λLC and μHC.

Abolishing the inter-chain disulfide bridge formation between μHC and λLC in three different mutant combinations did not alter the steady state subcellular distribution of participating subunits during IgM and IgM-like products biosynthesis (Fig. 11 D E). For the 2-chain co-expression setting to generate hexameric IgM-like products, regardless of whether the μHC or the λLC or both chains had the disulfide-ablating mutation, steady state subunit distribution was indistinguishable from that of the bona fide IgM (Fig. 11D, compare the top row with the second, third, fourth rows). The localization was mainly in the ER and cytoplasmic puncta as before. Similarly, for the 3-chain co-expression to produce pentameric IgM-like species, JC was consistently localized in the ER irrespective of whether μHC or λLC had the inter-chain disulfide-abrogating mutation (Fig. 11E, red), while the localization of [μHC + λLC] pair was generally in the ER and the puncta (Fig. 11E, green). The morphological evidence agreed well with the seemingly normal assembly and comparable secretion output of IgM-like products despite the lack of an inter-chain disulfide bridge between μHC and λLC.

To summarize, the prevention of inter-chain disulfide bond formation between μHC and λLC was tolerated in IgM covalent polymerization and secretion. In other words, non-covalent associations between μHC and λLC were strong enough to support IgM biosynthesis by meeting the ER quality control criteria and enduring the subsequent secretory pathway trafficking steps.

### 3.12. Disruption of μHC‒μHC covalent pairing by C337S mutation effectively prevents the formation of covalently associated IgM polymers

To further assess the roles of different inter-chain covalent associations in IgM biosynthesis, we next mutated the Cys-337 located in the CH2 domain. This Cys-337 is responsible for the inter-chain disulfide bridge that connects a pair of μHC monomers to generate a μHC homodimer comprising a single IgM monomer unit (see Suppl. 2A, third row). As the position of Cys-337 would implicate, the C337S mutation splits one IgM monomer molecule into two half-molecules at the midline. However, how the C337S point mutation affects the overall IgM product quality and secretion output is not clear. The types of assembly intermediates generated from this mutation are also unknown. Before characterizing the effect of C337S in the context of IgM biosynthesis, we first examined the behavior of μHC (C337S) mutant alone. Expectedly, μHC (C337S) showed a high propensity to induce Russell body (see Suppl. 2D) and was prevented from getting secreted into the culture media (see Suppl. 2 B C, lane 3). Even when μHC (C337S) was co-expressed with JC, due to its intact Cys-575, the μHC (C337S) mutant retained the ability to entangle JC into the Russell body (Suppl. 3D) and suppressed the JC secretion (Suppl. 3C, lanes 6 and 9).

As before, the protein expression level of each transfected subunit was comparable among cells that received different combinations of constructs in 2-chain and 3-chain settings (Suppl. 4F; lanes 1‒3 for 2-chain; lanes 4‒6 for 3-chain). Likewise, the amount of secreted subunit chains in culture media was equivalent regardless of whether the parental or mutant subunits were used (Suppl. 4 D E, lanes 1‒3 for 2-chain; lanes 4‒6 for 3-chain). Regarding the total amount of secreted subunits, the C337S point mutation had no harmful effect even when co-expressed with λLC-ΔCS mutant (Suppl. 4 D E, lanes 3 and 6).

Due to its critical position that can split the IgM monomer into two halves, it was not surprising that μHC (C337S) mutant subunit could no longer support the formation of covalently associated IgM-like proteins (Fig. 12A, compare lanes 1 and 2 for 2-chain; lanes 4 and 5 for 3-chain). In addition, having the C337S point mutation in the μHC subunit suddenly generated a much more diverse array of complex assembly intermediates (Fig. 12 A‒D, lane 2 for 2-chain; lane 5 for 3-chain). One such assembly intermediate species was much larger than ∼200 kDa, but its subunit composition or stoichiometry was unclear (Fig. 12 A B, lanes 2 and 5). When μHC (C337S) was co-expressed λLC-ΔCS, the complexity of secreted intermediates increased even further for the 2-chain setting (Fig. 12 A B, lanes 3). Because λLC-ΔCS would dissociate from μHC (C337S) in SDS-PAGE, the apparent laddering is caused by the difference in the number of μHC units covalently linked via intact Cys residues. Due to such narrow increments of size differences, it was unfeasible to annotate them reliably even when different components were separately probed (Fig. 12 B‒D). Of note, while all the tested μHC mutants so far failed to generate any assembly intermediates covalently associated with the JC, the μHC (C337S) mutant yielded two JC-positive assembly intermediates (Fig. 12D, lanes 5 and 6). A similar finding on the JC-containing assembly intermediates was reported by Davis et al. [73] back in 1989. However, according to Brewer and Corley (1997) [86], the JC incorporation into a polymerizing IgM can take place only at the final stage of IgM pentamer assembly because the JC incorporation is suggested to be thermodynamically favored over the incorporation of a sixth monomeric unit. Based on their size and reactivity to different probes, these intermediates are most likely to be μ2λ2J and μ2J (Fig. 12D, lanes 5 and 6). Although JC is usually not supposed to be found in smaller assembly intermediates, our results suggest that a unique configuration of μHC (C337S) open homodimers somehow mimics a particular conformation that favors the incorporation of JC via disulfide bond (see Fig. 15C).

**Figure 12.**
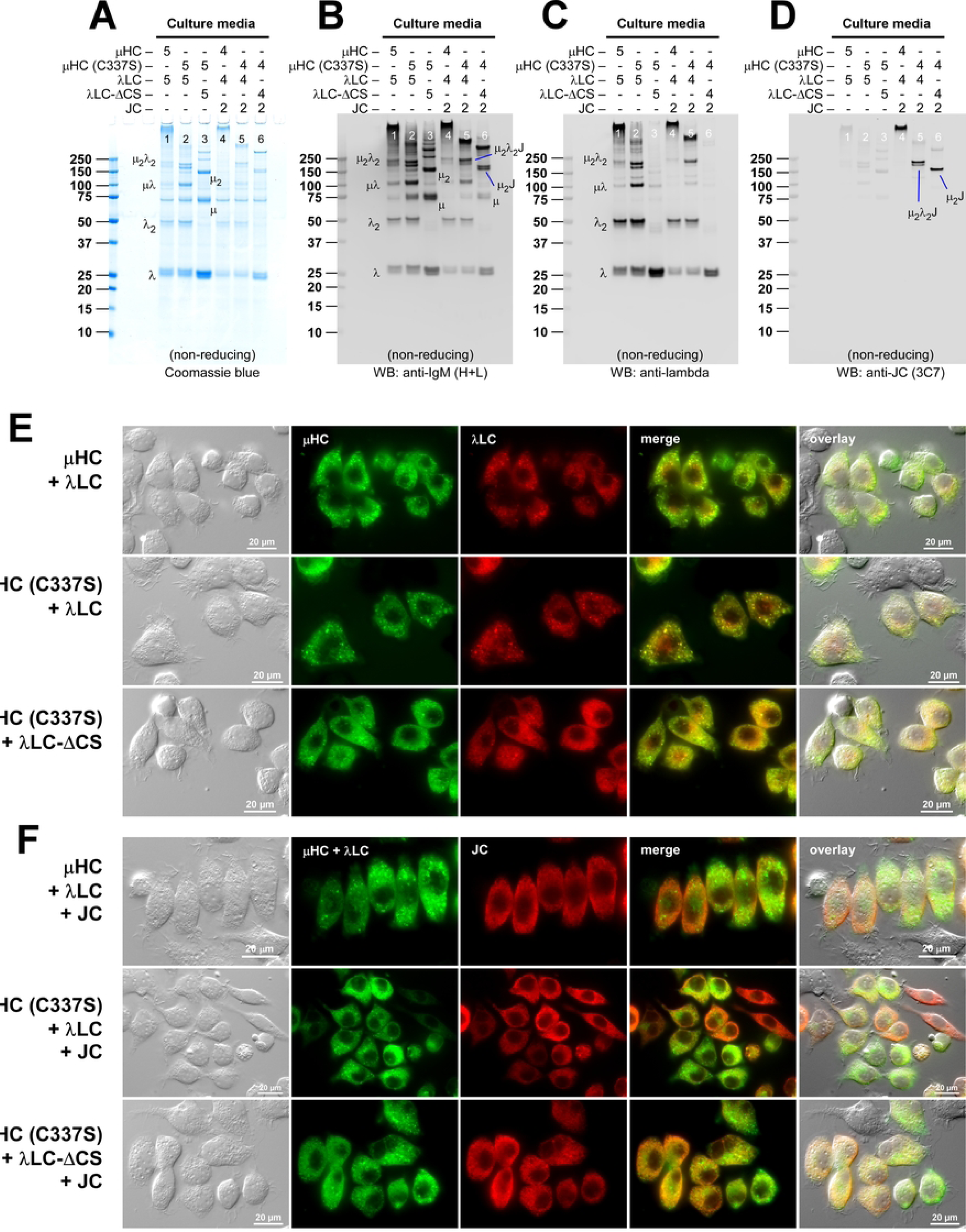
Effects of C337S point mutation on hexameric and pentameric IgM product formation and secretion. (A‒D) The role of Cys-337 in polymeric IgM formation was tested by replacing the parental μHC subunit with μHC (C337S) mutant in a 2-chain co-expression scheme (lanes 1‒3) and a 3-chain co-expression setting (lanes 4‒6). Subunit chains were co-transfected at the DNA ratio indicated at the top of each lane. Day-7 cell culture media were resolved by SDS-PAGE under non-reducing conditions, and Coomassie blue stained in panel A and analyzed by Western blotting in panels B‒D. Blotted membranes were probed with (B) polyclonal anti-IgM (H+L), (C) polyclonal anti-λLC, or (D) monoclonal anti-JC antibody. Protein bands corresponding to λLC monomers and dimers (or λLC-ΔCS monomers) and identifiable assembly intermediates are labeled on individual gels. (E, F) Fluorescent micrographs of HEK293 cells co-transfected with a combination of parental or mutant subunit chains. Cells were transfected with a 2-chain construct set (E) and 3-chain construct set (F) to generate hexameric and pentameric IgM-like products, respectively. The transfected subunit combination is shown on the left of each row. On day-3 post-transfection, cells were fixed, permeabilized, and co-stained with (E) FITC-labeled anti-human μHC antibody and Texas Red-labeled anti-human λLC antibody or (F) a 1-to-1 mix of FITC-labeled anti-human μHC and FITC-labeled anti-human λLC antibody cocktail (shown in green) and monoclonal anti-JC antibody followed by AlexaFluor594-conjugated secondary antibody (shown in red). Green and red image fields were superimposed to create ‘merge’ views. DIC and ‘merge’ were superimposed to create ‘overlay’ views.

**Figure 13.**
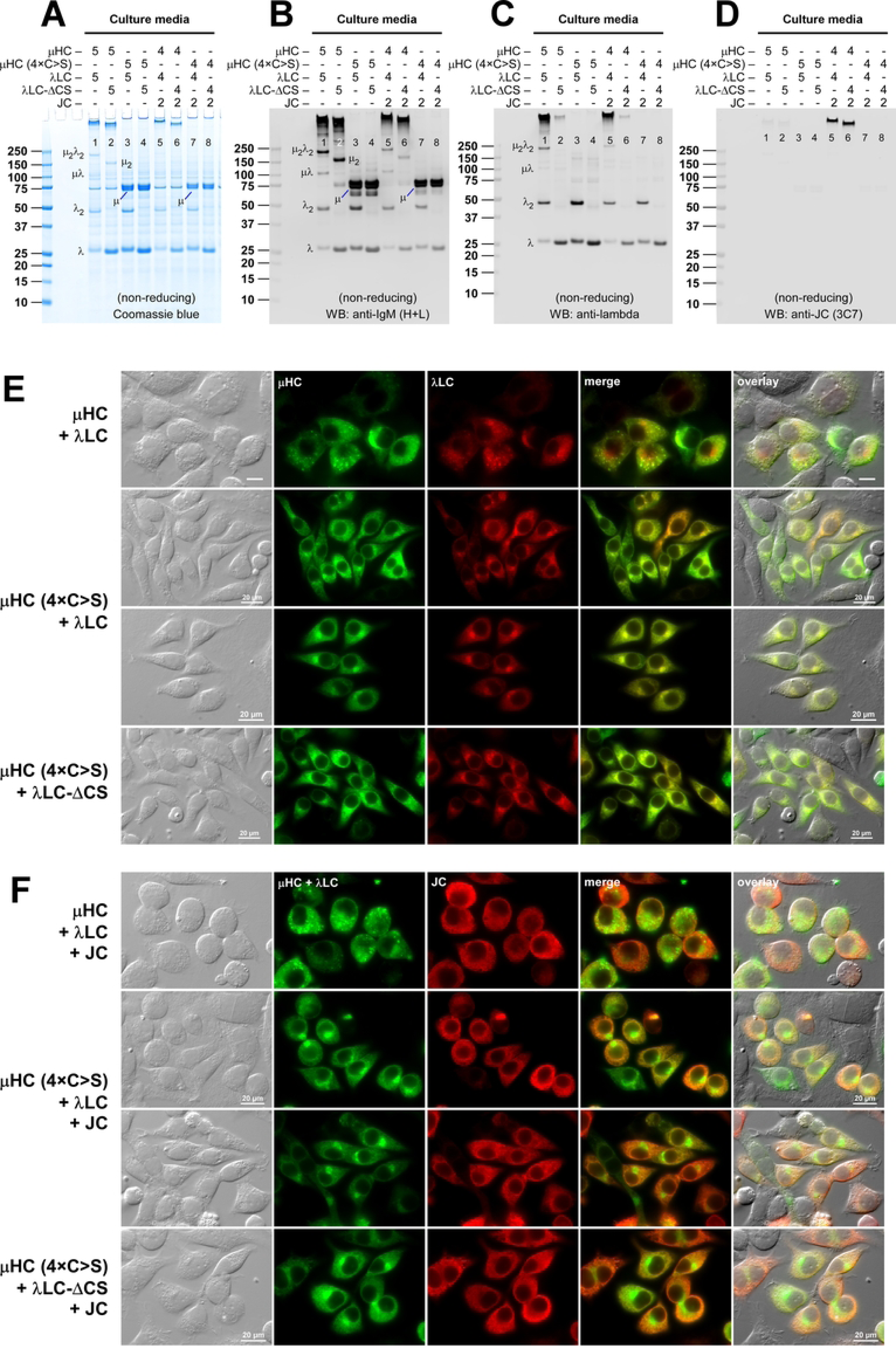
μHC subunit failed to produce covalently linked assembly intermediates when all four cysteine residues involved in inter-chain disulfide formation were mutated. (A‒D) All four Cys residues involved in inter-chain disulfide bond formation were abrogated by mutating Cys-137, Cys-337, Cys-414, and Cys-575 residues simultaneously into Ser to create μHC (4×C>S) mutant subunit. The effect of simultaneous inter-chain disulfide bridge ablation was tested on polymeric IgM formation by replacing the parental μHC subunit with μHC (4×C>S) mutant in a 2-chain co-expression scheme (lanes 1‒4) and in a 3-chain co-expression setting (lanes 5‒8). Subunit chains were co-transfected at the DNA ratio indicated at the top of each lane. Day-7 cell culture media were resolved by SDS-PAGE under non-reducing conditions, and Coomassie blue stained in panel A and analyzed by Western blotting in panels B‒D. Blotted membranes were probed with (B) polyclonal anti-IgM (H+L), (C) polyclonal anti-λLC, or (D) monoclonal anti-JC antibody. Protein bands corresponding to λLC monomers and dimers and identifiable assembly intermediates are labeled on individual gels. (E, F) Fluorescent micrographs of HEK293 cells co-transfected with a combination of parental or mutant subunit chains. Cells were transfected with (E) 2-chain and (F) 3-chain construct sets to generate hexameric and pentameric IgM-like products, respectively. The transfected subunit combination is shown on the left of each row. On day-3 post-transfection, cells were fixed, permeabilized, and co-stained with (E) FITC-labeled anti-human μHC antibody and Texas Red-labeled anti-human λLC antibody or (F) a 1-to-1 mix of FITC-labeled anti-human μHC and FITC-labeled anti-human λLC cocktail (shown in green) and monoclonal anti-JC antibody followed by AlexaFluor594-conjugated secondary antibody (shown in red). Green and red image fields were superimposed to create ‘merge’ views. DIC and ‘merge’ were superimposed to create ‘overlay’ views.

**Figure. 14.**
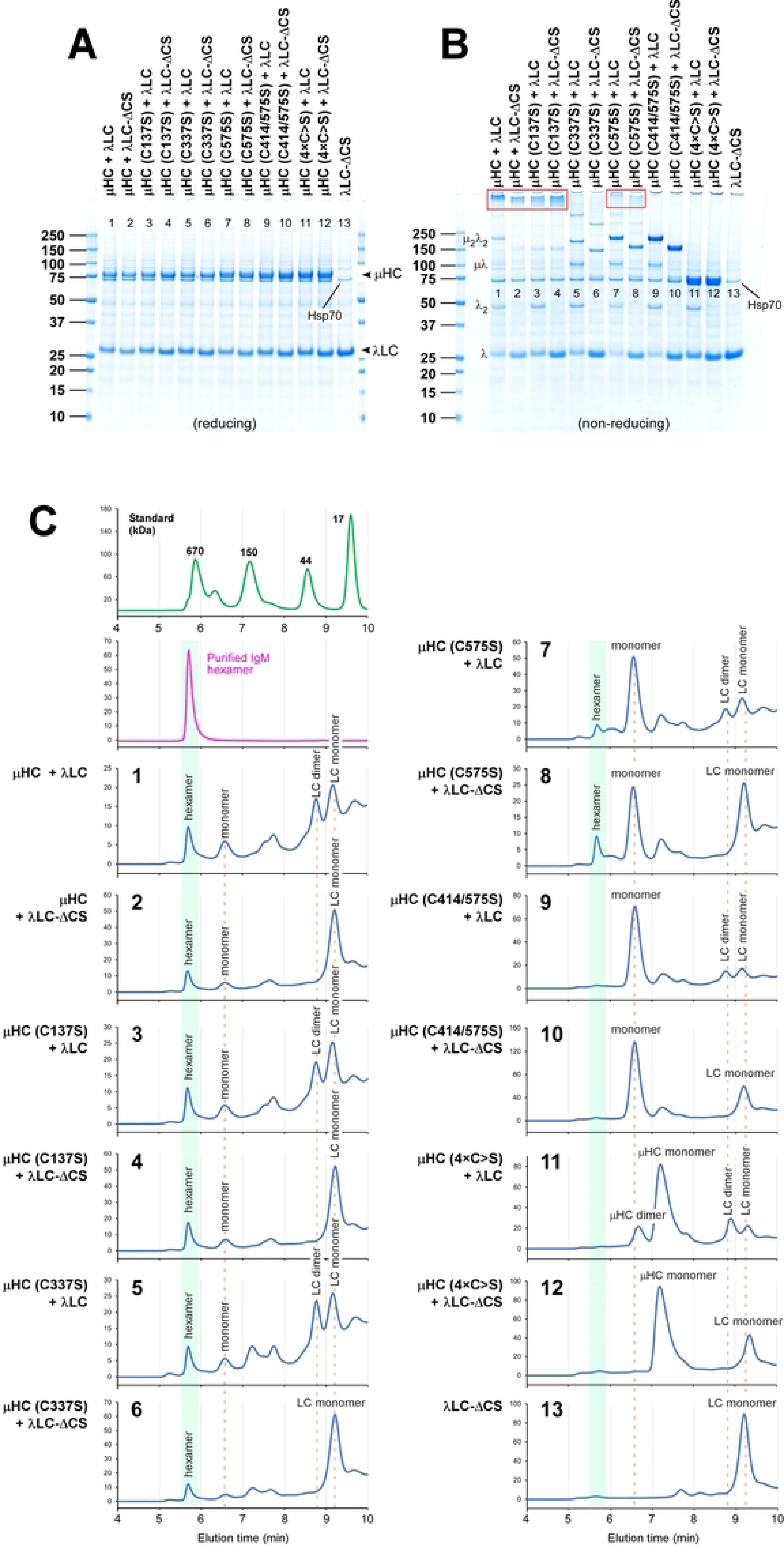
Assembly and maintenance of polymeric IgM products partially through non-covalent forces. (A, B) Secretion level and product quality of IgMs, IgM-like products, and assembly intermediates were compared in SDS-PAGE by resolving the day-7 cell culture media under (A) reducing and (B) non-reducing conditions. Parental μHC (lanes 1‒2) and five different μHC mutants (lanes 3‒12) were co-transfected with the parental λLC (odd number lanes) or λLC-ΔCS (even number lanes) in a 2-chain co-transfection scheme. In lane 13, the λLC-ΔCS construct was transfected by itself as a product quality control in the analytical SEC assay below. Covalently associated polymeric IgM and IgM-like products preserved in SDS-PAGE are marked by red box in panel B. (C) To assess the roles of non-covalent protein-protein interactions in the assembly and maintenance of polymeric IgM configuration, we examined the quality of secreted products by analytical SEC under non-denaturing and physiological pH assay conditions. Along with the day-7 culture media obtained from 13 different [μHC + λLC] subunit chain combinations shown in panels A and B, purified hexameric IgM (see Figs. 2E and 7B) was analyzed as a reference for polymeric IgM elution (panel C, second panel). SEC elution profiles for each culture medium sample shown in panel B, lanes 1‒13, are displayed in the chromatograms with the corresponding numbering in C, panels 1‒13. The transfected subunit chain combination is also shown on the left side of each chromatogram. The elution peak corresponding to hexameric IgM is shaded in light green in individual chromatograms.

**Figure 15.**
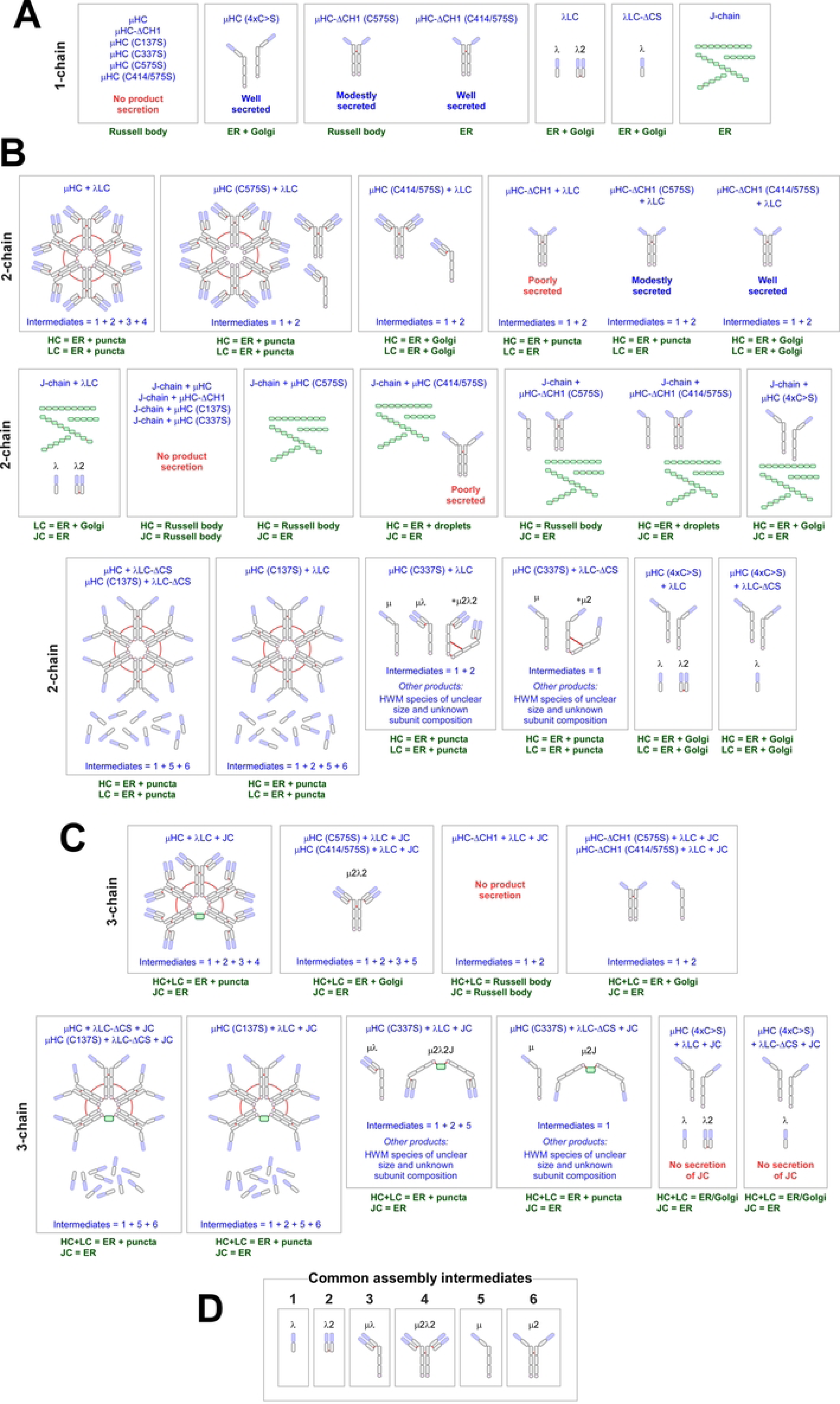
Diagrams of covalently assembled main secretory products, by-products, and assembly intermediates under different subunit combinations. The types of secreted main products and significant by-products released to culture media (or lack thereof) are illustrated. (A) single-construct expression setting. (B) 2-chain co-expression setting. (C) 3-chain co-expression setting. The name of the transfected construct (or a set of constructs) is listed in blue at the upper-most area of each box. At the lower-most area, common types of assembly intermediate in each setting are shown using the numbering system categorized in panel D. Steady state subcellular distribution of individual subunits for each transfection setting is shown in the space under each box. When there is no mention of the secretion outputs, it suggests that the products shown in the panel were secreted abundantly. (D) Six commonly produced assembly intermediates released to the culture media are illustrated in each box, from 1 to 6. This numbering scheme describes a set of secreted assembly intermediates in panels B and C. Solid red lines represent the inter-chain disulfide bond connectivity.

When it comes to the steady state subcellular distribution of μHC (C337S) and various assembly intermediates containing the μHC (C337S) subunit, overall distribution remained the same as the parental [μHC + λLC] pair co-expression, regardless of 2-chain (Fig. 12E) or 3-chain (Fig. 12F) expression setting. In all cases, μHC and λLC were collectively found in the ER and cytoplasmic puncta (Fig. 12 E F), while the JC was predominantly in the ER (Fig. 12F). A blockade of inter-chain disulfide formation between the two μHC monomers within an IgM monomer unit disturbed the covalent polymerization of IgM product, but the effect of C337S was still tolerated by the ER quality control mechanisms to allow the secretion of various covalent assembly intermediates.

### 3.13. Simultaneous ablation of all four cysteine residues involved in inter-chain disulfide formation prevents the μHC from producing covalent assembly intermediates

To assess the effect of complete ablation of inter-chain disulfide bonds on polymeric IgM biosynthesis, we created a μHC (4×C>S) mutant subunit in which Cys-137, Cys-337, Cys-414, and Cys-575 were simultaneously mutated to serine (Suppl. 5A). Before tested in IgM assembly reaction, we first examined the characteristics of μHC (4×C>S) subunit by itself. Unlike the parental μHC and other μHC mutants that induced Russell body phenotype (Suppl. 5F, first row), the cells expressing μHC (4×C>S) mutant maintained normal cell morphology, and the μHC (4×C>S) protein distributed to the reticular ER without showing any sign of aggregating into Russell body (Suppl. 5F, second and third rows). Likewise, unlike the parental μHC and other μHC mutants that were barred from secretion (Suppl. 5 D E, lane 1), μHC (4×C>S) mutant was abundantly secreted by itself as monomers, and to a lesser extent as dimers and tetramers when expressed alone (Suppl. 5E, lane 2). If μHC (4×C>S) was co-expressed with JC, unlike the parental μHC that co-aggregated with JC into Russell body (Suppl. 5G, top row), both μHC (4×C>S) and JC maintained their respective ER distribution as if they were expressed individually (Suppl. 5G, second and third rows) and both proteins were secreted to the culture media as they normally would (Suppl. 5 D E, lanes 3‒4).

To assess the biosynthetic assembly of polymeric IgM in the complete absence of inter-chain disulfide bond formation, μHC (4×C>S) mutant was co-expressed with the parental λLC or its disabled λLC-ΔCS counterpart, both in the absence and presence of JC. As always, the protein expression level of each transfected construct was validated using the cell lysates by Western blotting to ensure that designated proteins were expressed at equivalent amounts under different construct combinations (Suppl. 4 I). We also verified the secretion of each transfected subunit under reducing conditions by Coomassie blue staining and Western blotting (Suppl. 4 G H) and found that there was no marked effect of μHC (4×C>S) mutation on total protein secretion levels (Suppl. 4G).

To evaluate the effect of μHC (4×C>S) mutation on polymeric IgM assembly and product quality, the secreted proteins were resolved under non-reducing conditions (Fig. 13A). Because μHC (4×C>S) can no longer form any inter-chain disulfide bridge with other proteins including itself, the detectable secretory products were predominantly the μHC (4×C>S) monomers without producing any covalent assembly intermediates (Fig. 13B, lanes 3‒4 and lanes 7‒8). Unlike when μHC (4×C>S) was expressed alone or with JC (Suppl. 5E), dimers or tetramers were not detected in the culture media whenever co-expressed with λLCs (Fig. 13B, lanes 3‒4 and lanes 7‒8). The co-expressed λLC or λLC-ΔCS was independently secreted by following their dimer/monomer forming rules (Fig. 13C, lanes 3‒4 and 7‒8). Expectedly, JC did not form any covalent assembly intermediates with μHC (4×C>S) (Fig. 13D, lanes 7‒8). Therefore, by the ablation of all five cysteine residues involved in inter-chain disulfide formation (four on μHC and one on λLC), we can completely block the formation of any covalently assembled intermediates (Fig. 13 A B, lanes 4 and 8).

Steady state subcellular distribution of co-expressed [μHC (4×C>S) + λLC] pair and [μHC (4×C>S) + λLC-ΔCS] pair (Fig. 13E, second to fourth rows) was clearly different from that of the parental [μHC + λLC] pair which reproducibly showed the ER and cytoplasmic puncta localization (Fig. 13E, top row). Instead, μHC (4×C>S) distributed to the ER and Golgi (Fig. 13E, second to fourth rows, green), while the co-expressed λLC or λLC-ΔCS distributed to the ER and Golgi as if these subunits were expressed separately (Fig. 13E, second to fourth rows, red). Co-expression of λLC and λLC-ΔCS did not influence the subcellular and secretory behaviors of μHC (4×C>S) at all. Likewise, when JC was co-expressed in the 3-chain expression setting, μHC (4×C>S) and λLC or λLC-ΔCS remained distributed to the ER and Golgi, whereas JC continued to show ER localization (Fig. 13F, second to fourth rows, red). Again, whenever μHC (4×C>S) was involved, subunit distribution no longer showed the cytoplasmic punctate staining that was a possible feature of the covalent μHC multimerization in the ER (Fig. 13F, top row, green).

The complete ablation of all inter-chain disulfide bonds among all participating subunit chains abolished the formation of any detectable covalently linked assembly intermediates. As a result, μHC (4×C>S) subunit was secreted as monomers without covalently associated with other co-expressed subunits.

### 3.14. Polymeric IgM product integrity is partly maintained through non-covalent forces even when certain inter-chain disulfide bonds are missing

Biochemical studies carried out by Parkhouse et al. [87] in 1970 showed that polymeric IgM molecules can be reconstituted from a mixture of partially reduced and alkylated IgM monomers obtained from mouse plasmacytoma MOPC 104E. Kownatzki [88] reported similar results in 1973 that after a full reduction of pentameric IgMs into HCs, LCs, and JCs, pentameric IgMs were restored by removing the reducing agent to allow oxidation. Similarly, in 1975, Parkhouse [89] reported that a reduced and alkylated mix of IgM monomers and H-L half-molecules could associate through non-covalent forces to form an IgM species that sedimented at 19S, which reflected the re-formation of polymeric IgMs. Furthermore, Eskeland [90] showed in 1978 that IgM polymers can arise from a mixture of free H-L half-molecules and free JCs obtained after reduction but without alkylation. In the same study, Eskeland not only found that sulfhydryl groups must be intact and zinc ion needs to be present for the non-covalent polymerization process to take place but also demonstrated that non-covalently associated polymers could gradually become disulfide-linked polymers during prolonged dialysis. While these past studies highlighted the importance of non-covalent forces when reconstituting polymeric IgMs from a different mixture of intermediates and subunits in test tubes, it is not known whether a multimeric IgM can be generated or maintained solely or partially through non-covalent forces during biosynthesis in the ER and secretory trafficking.

Up to this point, the product quality of secreted IgMs, IgM-like molecules, and assembly intermediates were analyzed by resolving the proteins in SDS-PAGE under non-reducing conditions. Although this method preserves pre-existing covalent protein-protein interactions via disulfide bridges, non-covalent associations are disrupted because SDS is a denaturant. In order to assess which specific cysteine residues and which inter-chain covalent interactions are dispensable (or critical) in the biosynthetic formation of polymeric IgM, this time, we examined the quality of secreted products by analytical SEC under non-denaturing and physiological pH assay conditions. We presumed that most of the involved non-covalent protein-protein associations would be preserved in such assay settings and provide us with a more accurate account of the state of secretory products. Our goal is to determine which specific cysteine can be eliminated without losing the ability to assemble and secrete polymeric IgMs solely or partly through non-covalent forces.

To prepare samples for analytical SEC, the parental μHC and its five different mutant variants (i.e., C137S, C337S, C575S, C414/575S, and 4×C>S) were co-expressed with the parental λLC or the λLC-ΔCS mutant side by side in the 2-chain expression settings. On day-7 post-transfection, cell culture media were harvested and analyzed by SDS-PAGE to assess protein secretion and product quality before they were subjected to analytical SEC. Firstly, consistent with all the previous results, secretory protein levels looked comparable across the board when analyzed under reducing conditions (Fig. 14A, lanes 1‒12). Secondly, under non-reducing conditions, IgMs, IgM-like proteins, and various assembly intermediates resolved into reproducible banding patterns according to the expected characteristics unique to each mutant (Fig. 14B, lanes 1‒12).

Analytical SEC clearly showed that parental [μHC + λLC] pair co-expression yielded polymeric IgMs and monomeric IgMs as the two recognizable main products in addition to the LC dimer, LC monomer, and one unknown intermediate that eluted at ∼7 min 45 sec (Fig. 14C, panel 1). Similarly, the [μHC + λLC-ΔCS] pair produced and secreted polymeric IgMs, monomeric IgMs, and the LC monomers (Fig. 15C, panel 2). Even when tested in the 3-chain expression setting to produce pentameric IgMs, parental λLC and λLC-ΔCS, both produced similar-sized polymeric IgM products in analytical SEC (Suppl. 6, A‒C). Likewise, μHC (C137S) mutant retained the ability to assemble polymeric IgM species (through non-covalent interactions between HC and LC) when co-expressed with the parental λLC (Fig. 14C, panel 3) or λLC-ΔCS (Fig. 14C, panel 4). These first four SEC results (or six SEC results including Suppl. 6) solidified the earlier conclusions that the inter-chain disulfide bond between μHC and λLC was dispensable for the polymeric IgM assembly, product integrity, and secretion.

How about the inter-chain disulfide bond that holds the pair of μHC monomers within an IgM monomer unit? Is this inter-chain disulfide bond required for polymeric IgM assembly? Although we did not detect any high molecular weight range products in non-reducing SDS-PAGE when μHC (C337S) mutant was tested (Fig. 14B, lanes 5‒6), SEC data demonstrated that μHC (C337S) subunits retained the capacity to assemble into polymeric IgMs through non-covalent forces (Fig. 14C, panels 5 and 6). Therefore, the inter-chain disulfide bridge formed between the two Cys-337 residues of the juxtaposing μHC monomer pair was not an absolute requirement for the maintenance of multimeric IgM product.

As shown repeatedly in Fig. 5D (lanes 2 and 8) and also in Fig. 14B (lanes 7‒8), μHC (C575S) mutant marginally retained the ability to generate covalently-associated polymeric IgM products via inter-chain disulfide interactions mediated by the intact Cys-414 residue. SEC under physiological conditions also showed that μHC (C575S) mutant retained the ability to form polymeric IgM species partly through non-covalent interactions (Fig. 14C, panels 7 and 8). However, when both Cys-414 and Cys-575 were mutated simultaneously, such μHC (C414/575S) double mutant subunit was no longer able to produce polymeric IgM species either through covalent or non-covalent interactions (Fig. 14C, panels 9 and 10). Instead, the monomer unit of IgM predominated the secretory products, and there was no detectable peak that corresponded to the polymeric IgMs. Similarly, when all four cysteine residues involved in the inter-chain disulfide formation were mutated to serine, μHC (4×C>S) mutant was almost exclusively secreted as μHC (4×C>S) monomers and lost the ability to form any assembly intermediates altogether (Fig. 14C, panels 11 and 12). In addition, the λLC-ΔCS mutant was also expressed by itself (Fig. 14 AB, lane 13) and analyzed by SEC (Fig. 14C, panel 13). The SEC results demonstrated that non-covalent association alone could not generate LC dimers, clearly indicating an obligatory role of the disulfide bridge formation in λLC dimerization. On a side note, a similar SEC analysis was also performed for the pentameric IgM mutant series using harvested cell culture media. Unfortunately, most likely due to covalently aggregated free JCs in the culture media, the elution peak area corresponding to pentameric IgM was masked (data not shown) and thus hampered our data analysis.

In summary, single point mutations such as C137S, C337S, or C575S on μHC and ΔCS on λLC were tolerated for polymeric IgM assembly and product secretion. In other words, not all the prescribed inter-chain disulfide bonds need to be in place for polymeric IgM assembly and secretion. Inter-chain interactions mediated by non-covalent forces can withstand the loss of certain inter-chain disulfide bonds to maintain the multimeric IgM product integrity. By contrast, the simultaneous loss of two inter-chain covalent interactions by the C414/575S double mutant was too disrupting for the non-covalent forces to maintain the polymeric IgM product integrity during biosynthesis and secretion.

## 4. Discussion

IgMs are megadalton-size secretory proteins that continue to attract cell biologists to investigate how they negotiate their abundant secretion with the ER quality control mechanism despite their structural complexity and biosynthetic burden [18, 57, 70, 72, 91]. To further probe the details of IgM assembly and secretion requirements, we characterized the biosynthetic process of human IgM in a fully recombinant setting using a natural human IgM called SAM-6 as our model cargo. By creating a series of mutant subunit constructs that disrupt the formation of specific inter-chain disulfide bonds in a targeted manner, we tested their effects on various aspects of IgM biosynthesis in 48 combinations. Specifically, we investigated the effect of mutations on (1) protein expression level and secretion output, (2) steady state subcellular distribution of subunits during biosynthesis, (3) associated intra-ER inclusion body formation, and (4) secreted product quality and types. Through our comparative approach that uniquely combined the analysis of intracellular phenotypes and extracellular secreted product quality, we not only uncovered new features that govern the courses of IgM assembly and secretion, but also clarified underlying reasons behind conflicting reports on Russell body induction during IgM expression. This study also helped solidify a series of previously proposed concepts for IgM.

### 4.1. Not all the inter-chain interactions need to be stabilized by the covalent disulfide linkage to produce polymeric IgMs

One of the unexpected findings was the evidence that not all the prescribed inter-chain disulfide bridges were required for the polymeric IgM formation. In other words, a complete set of inter-chain disulfide formations is not the obligatory requirement for the assembly and secretion of polymeric IgMs. Although seemingly important, inter-chain disulfide bond formation does not appear to be the most critical attribute for the ER quality control surveillance system to assess the product quality and secretion readiness of assembling IgMs. The evidence suggested that inter-chain interactions can be stabilized through non-covalent protein-protein interactions at least in three locations: (1) between μHC (Cys-137) and λLC (Cys-213) both for hexamers and pentamers, (2) between the pair of juxtaposing μHC monomers via Cys-337 at least for hexamers, and (3) between the two μHC covalent homodimers via Cys-575 at least for hexamers. We did not evaluate the effect of single point mutation on Cys-414 based on the conclusion from Wiersma and Shulman [92] suggesting that “C414 is not of great importance in assembly” of mouse IgM. Importantly, the C414/575S double mutation revealed that non-covalent interactions alone were not strong enough to compensate for the simultaneous loss of two inter-chain disulfide bonds. It is fascinating to imagine that non-covalently held products can fulfill the ER quality control criteria and remain assembled during secretory pathway trafficking and even after the products are released to the extracellular space. This feature highlighted the vital roles of underlying non-covalent forces that not only orchestrate the initial subunit interactions, but also maintain the polymeric IgM product integrity during and after secretion. We propose that the IgM assembly process is far more robust than it appears and equipped with a stop-gap measure that tolerates the secretion of polymeric IgM even when a particular set of inter-chain disulfide bond formation is incomplete. Such biosynthetic robustness may underwrite the abundant product secretion not just only from a fully differentiated professional secretory plasma cell, but also from a generic cell like HEK293.

While the dispensability of inter-chain disulfides seemed striking initially, it was not unusual from the standpoint of immunoglobulin evolution in vertebrate species. If we look beyond the human IgMs, there are many examples of normal immunoglobulins in which interactions between HC and LC are non-covalently maintained. For instance, in mouse IgA [93] and human IgA2 m(1) allotype [94, 95], a disulfide-linked LC dimer (LC‒LC) and a disulfide-linked HC dimer (HC‒HC) assemble non-covalently to produce a functional IgA antibody without inter-chain disulfides between HCs and LCs. Similarly, IgA purified from chicken bile lacked the inter-chain disulfide bridges between HC and LC, and the IgA was shown to dissociate into αHC homodimers and free LC monomers without breaking disulfide bonds [96]. Canine, equine, and porcine milk also contain IgA that dissociates into HCs and LCs without disulfide reduction [96]. In an extreme case, four different mouse monoclonal IgA mAbs were shown to dissociate into free HC monomers and free LC monomers in SDS-PAGE under non-reducing conditions; and upon SDS removal, these monomers of HC and LC assembled back into functional IgAs that retained the antigen binding function [97]. In the studies on the primitive jawless vertebrates such as hagfish [98] and sea lamprey [99], antibodies seem to exist in 6.6S and 14S forms and behaved like monomeric and polymeric IgMs, respectively, but they lacked inter-chain disulfide linkage between HCs and LCs. Furthermore, in bullfrogs (*Rana catesbeiana*), free LC monomers are non-covalently associated with the disulfide-linked HC homodimers in all classes of immunoglobulins [100]. A similar feature was reported for the IgY of cane toad (*Bufo marinus*) [101]. There is yet another trick to do away with the inter-chain disulfide between HC and LC. In the μHC primary sequences of green anole lizard (*Anolis carolinensis*) and Chinese crocodile lizard (*Shinisaurus crocodilurus*), the critical Cys residue supposed to be present in the CH1 domain is absent to begin with [102, 103]. Moreover, in Chinese crocodile lizard, the LC sequence additionally lacked the critical C-terminal Cys residue [103] responsible for the inter-chain disulfide bonding. In this lizard species, the inter-chain disulfide bond formation between HC and LC is reciprocally blocked on both subunits as if making sure to thwart the formation of an inter-chain disulfide bond. As a result, the HC and LC of lizard polymeric IgM are held together only by non-covalent forces [102, 103]. Unlike the Cys residues involved in intra-domain disulfide bond formation that are stringently conserved among different immunoglobulins, the heterogeneity in inter-chain disulfide connectivity, position, and numbers among different immunoglobulins suggests that “these bonds may be of lesser importance to the antibody functions of mammalian immunoglobulins [94].” In mouse IgG2b, for example, when the critical Cys-128 in the CH1 was mutated to Ser, the covalent γ2b-HC homodimers retained the ability to assemble non-covalently with the covalent homodimers of mouse λLC into a secretion competent, functional antibody [85]. Therefore, what we uncovered in this study for human IgM is evolutionarily and mechanistically prevalent in some branches of the animal kingdom or different classes of immunoglobulins. These examples suggest that the LCs do not always need to associate with the HCs via a covalent disulfide bond to generate a functional immunoglobulin. The blueprint for polymeric IgM assembly is therefore embedded in the individual subunits, but more than one type of polymeric IgM product can be manufactured and classified as “normal” by the ER quality control system.

Is there any benefit of releasing such “incomplete” IgM species in haste without dotting the i’s and crossing the t’s? ― We speculate that, especially during an urgent response against the threat of infectious agents, the benefit of secreting such molecules may outweigh the risk of delaying the quantitative response. Besides, because the ER quality control mechanisms have already endorsed their quality for abundant secretion, there would be little harm to the host even if plasma cells released such molecules. However, the precise effects of missing a specific inter-chain disulfide bond (e.g., between μHC and λLC) on antigen binding as well as another inter-chain disulfide (e.g., between μHC monomers mediated by Cys-337) on cytolytic activity or epithelial transport are not understood―and are worth investigating.

### 4.2. Full list of secretory products and intermediates

This study revealed the production and release of various assembly intermediates and by-products during the parental and mutant IgM expression. Interestingly, some products were stringently retained in the cell due to the strict ER quality control mechanism, while others were secreted to the culture media abundantly, depending on which subunit was expressed and which mutation was introduced to the subunit. To holistically view the collection of products and by-products identified and characterized in this study, we collated the snapshot information and displayed them as an inclusive reference chart (see Fig. 15). The chart shows (1) steady state subcellular cargo distribution during overexpression, (2) the types of inclusion body the cargo can induce, (3) the types of main secretory products and major by-products, (4) a subset of common assembly intermediates released to the culture media, and (5) secretion outputs. We made this chart hoping to serve as a reference guide to determine what can happen to IgM assembly and secretion when specific mutations are introduced to HCs and LCs and when a certain subunit chain expression is omitted. Our study increases the awareness of IgM and IgM-like proteins as a promising modality option for biotechnology applications by lowering the hurdle to working with recombinant IgMs.

### 4.3. Prospects of IgM-like scaffold for therapeutic protein design

Because a pool of secreted IgM products may contain a small fraction of IgM molecules lacking the inter-chain disulfide bonds between HC and LC or elsewhere (and can go unnoticed), manufacturing a biochemically, biophysically, and functionally homogeneous drug substance and maintaining a consistent product quality can be challenging. While establishing a panel of qualifying assays for both physicochemical attributes and biological functions would be necessary without a doubt, coming up with ingenious protein engineering strategies may be another viable approach to prevent such potential product heterogeneity introduced by the lack of certain inter-chain disulfide bridges, especially between HC and LC.

One simple approach to circumvent such product heterogeneity may be to stop using the LC subunit altogether and re-design the IgM’s antigen binding domain using the VH fragment from a class of HC-only antibodies. For example, UniDab^TM^ [104] and Nanobody® [105] can recognize antigens at high affinity as a self-sufficient unit without the need for VL-derived sequences (unlike Fab, scFab, or scFv). Such VH-based binders are convenient modules when generating multivalent IgM-like molecules both in hexameric and pentameric forms by simply fusing to the μHC’s constant region (see Suppl. 7B). Compared to the conventional IgMs (Suppl. 7A), this molecule retains the benefit of antigen binding avidity while eliminating the need for μHC‒LC pairing. In addition, because the single-domain antibody is regarded as a promising building block when designing a multi-specific antibody [104, 105], different UniDabs or Nanobodies can be fused directly to the N- or C-terminus (or both termini) of the JC subunit to generate a multivalent bi- or tri-specific molecules (Suppl. 7C). Furthermore, if we combine the VH fragment binding domains with a heterodimerizing Fc technology, one can envision to create a variety of multi-specific IgM molecules (Suppl. 7D). Because we no longer need to use LC-derived sequences, using a VH based binder module should simplify the molecular design, on one hand, and increase the design versatility, on the other hand, while simultaneously eliminating the HC‒LC interaction heterogeneity. Although the simpler design makes good sense, given that SAM-6 μHC-ΔCH1 alone (Fig. 15A) and [SAM-6 μHC-ΔCH1 + JC] co-expression (Fig. 15B, second row) failed to yield secretion-competent products. Whether these novel molecules built from UniDab™ or similar VH domain-only binder (see Suppl. 7 B‒D) would achieve secretion competency must be fully addressed.

While the JC subunit is an important piece of the IgM molecular design puzzle, it has remained largely unexplored as the substrate for protein engineering. JC subunit is shared with another multivalent immunoglobulin class, IgA, and is best known for its interaction with pIgR to facilitate epithelial transport [15, 16]. However, when it comes to its biosynthesis and native conformation, it has not been understood well [106]. The fact that JC is also expressed in the cells that do not express IgM or IgA, or even in the cells that do not express HCs or LCs, is vexing, and it might suggest that JC may have additional functions [107]. In fact, by just being present in the ER lumen, JC modulated the intracellular solubility of other subunits such as [μHC-ΔCH1 + λLC pair] by changing their steady state punctate localization into inducing Russell bodies. JC also altered the steady state dynamics of μHC (C414/575S) and μHC-ΔCH1 (C414/575S) to induce droplet inclusion bodies. How JC subunits modulate other subunits’ solution behavior is not understood. Likewise, the mechanism for why only one JC subunit is incorporated into an assembling pentameric IgM is not fully understood. JC is also a unique protein in that (literally) it does not show homology to any known proteins except for the JCs from other species, and as such, it cannot be placed into a structurally related protein family [77, 106]. A single JC molecule has 8 cysteine residues involved in 3 intra-molecule disulfide bridge formation [77]. While disulfide bond connectivity is understood in the context of pentameric IgM and IgA [83, 108], JC’s structural features (PDB ID: 7K0C for IgM, 6UE7 for IgA) suggested that both termini are amenable to fuse with a functional protein module without hindering the incorporation into a pentameric IgM. This approach has already been tested in the examples of IGM Bioscience’s (https://igmbio.com/) molecular design, where they engineer multivalent bi-specific IgM by fusing an anti-CD3 scFv to the JC subunit [24, 32]. However, details need to be determined empirically regarding the size limit of guest proteins that can be fused on either terminus or both termini.

Another important aspect of IgM we did not cover in this study was the non-covalent interactions between the J-chain of a pentameric IgM and the secretory component (SC) [109]. To enable mucosal and nasal delivery of biologic drugs, optimizing a method to produce a secretory IgM complex (i.e., pentamer IgM + SC) would be crucial to mimic how nature delivers antibodies to external secretions by epithelial transport. Given that the association of SC was shown to protect the IgA from proteolytic degradation in a rather harsh environment of mucosal surface [110], SC is expected to provide similar protection to pentameric IgM, thereby potentially prolonging the half-life of secretory IgM in the lumen and mucosal surface.

### 4.4. Requirements for the specialized chaperones upregulated in IgM-producing plasma cells

Despite the intrinsic molecular complexity and the demand for cellular resources to produce IgM molecules, an early study published in 1986 reported that plasma cells can produce and secrete ∼25,000 IgM molecules per cell per second [19]. To explain the cellular feat like this, it has been postulated that plasma cells acquire enhanced secretory capacity by upregulating specialized ER resident chaperones during a differentiation process into a mature professional secretory cell to assist the assembly of IgM molecules [58–60, 111]. Interestingly, Cattaneo and Neuberger [74] showed that glioma, phaeochromocytoma, and other non-lymphoid cell lines were able to secrete polymeric IgM upon transfection of μHC and LC to a comparable level as plasmacytoma cell hosts. Likewise, Niles et al. [76] reported that IgM can be assembled and secreted by a murine AtT20 pituitary cell line regardless of developed or rudimentary secretory apparatus and without requiring specialized factors only available in plasma cells. Cell types of non-lymphoid origin are also sufficiently equipped to produce both pentameric and hexameric IgMs if the cells were transfected with μHC and λLC constructs with or without the JC subunit.

Now, because our model HEK293 cell was also able to assemble and secrete both pentameric and hexameric forms of IgM readily, our study agreed that B-cell and plasma cell-specific ER resident proteins and chaperones were not required for the formation and secretion of polymeric IgMs. Although these factors may not be necessary for IgM secretion per se, it is still an intriguing possibility that plasma cell-specific factors play more critical roles in increasing the success rate of polymeric IgM assembly while decreasing the release of assembly intermediates such as monomeric IgMs (μ2λ2) and half-molecules (μλ). Although this possibility was not tested in this study, we are curious about the specific effects of pER1p [58, 60] and ERp44 [59, 111] on the secreted IgM product quality. It will be essential to demonstrate whether the ectopic expression of those factors maximizes the secretion of fully assembled IgM products from a non-professional secretory cell such as HEK293 or CHO and minimizes the release of assembly intermediates.

### 4.5. Russell body-related inclusion body phenotypes during IgM biosynthesis

Induction and suppression of Russell body phenotype during SAM-6 IgM biosynthesis was mostly predictable and agreed well with previously reported findings from the studies on various recombinant IgG mAbs characterized in our laboratory [50–56, 84]. We said “mostly” because there were cases where Russell body phenotypes and cargo secretion needed to be interpreted carefully because of the unique characteristics presented by the penultimate Cys residue located in the μHC’s C-terminal tailpiece; the Cys that plays roles in JC association and IgM polymerization and that is also a known substrate of free thiol mediated ER retention mechanism by forming a reversible mixed disulfide bond with ERp44 [111].

Just like a group of γHC subunits characterized previously, μHC expression by itself invariably induced Russell body even when the Cys residues involved in inter-chain disulfide bonds were mutated singly or doubly unless all the four Cys residues were mutated simultaneously. In this respect, Russell body formation served as an unequivocal visual cue that reports the lack of cargo secretion. Likewise, the μHC-ΔCH1 mutant induced Russell body and its secretion was suppressed until both Cys-414 and Cys-575 were mutated simultaneously, upon which the Russell body phenotype was dissolved, and the cargo secretion was restored. Similarly, co-expression of μHC and JC (as well as μHC-ΔCH1 and JC) led to the co-aggregation of both subunits in the ER and induced Russell body. Upon mutating the Cys-575 into Ser, JC became excluded from the Russell body and returned to its normal ER distribution. Although it has been well known that μHC is not secreted by itself in the absence of LC co-expression, our work was the first instance where the underlying “cell phenotype‒secretion relationship” was clearly demonstrated and explained visually by immunofluorescent microscopy.

The ΔCH1 version of immunoglobulin μHCs has been used as a convenient model to induce Russell body phenotype in some influential studies [68, 69] and led to a widespread belief that HC’s sequence abnormality like ΔCH1 is the main cause of Russell body formation. This notion was later refuted because structurally and functionally normal human IgGs can also induce Russell bodies depending on the variable domain sequences, cellular protein homeostasis, and IgG’s physicochemical properties [50, 51, 53]. In any event, Valetti et al. [68] used two different homologous cell models in which endogenous immunoglobulin genes were differentially expressed: mouse myeloma J558L (H–/L+/J+) and mouse myeloma NS0 (H–/L–/J+). They also tested in a heterologous cell model, rat glioma cells, in which no endogenous immunoglobulin genes are expressed (H–/L–/J–). After ectopically expressing full-length μHC, μHC-ΔCH1, and λLC in those three cellular models, Valetti et al. [68] concluded that the assembly and polymerization of [μHC-ΔCH1 + λLC] was required to induce Russell body. They also asserted that the Russell body formation by [μHC-ΔCH1 and λLC] was independent of JC availability because rat glioma cells showed punctate cargo accumulation which they interpreted as Russell body.

In a more recent study, Corcos et al. [67] reported a result that conflicted with Valetti’s conclusion. Corcos et al. compared the Russell body forming frequency of CD138+ plasma cells derived from two different strains of mice. The first was knock-in mice (denoted μNR) expressing a truncated μHC lacking the CH1-CH2 domains (ΔCH1-CH2). The second was from μNRL−/− mice that again express the same truncated μHC (ΔCH1-CH2) but in an LC null background. The results revealed that plasma cells of μNRL−/− mice induced Russell bodies extensively, whereas the same μHC (ΔCH1-CH2) protein did not aggregate into Russell body when LC was present in the μNR mice background. Here, although the truncated μHCs and the LCs did not assemble covalently because of the ΔCH1-CH2 truncation, the results suggested a protective role of LCs in preventing the truncated μHCs from aggregating into Russell bodies during IgM biosynthesis in vivo. Because the authors gave no account of the JC expression in their cells, whether the JC subunit played any role in the Russell body formation was unknown from their study.

Now, a widely used heterologous cell host HEK293 turned out advantageous over homologous cellular models to explain these conflicting results because HEK293 cells do not express endogenous immunoglobulin genes that may complicate or mislead the data interpretation. When similar experiments were carried out in a straightforward recombinant setting, co-expression of cognate LC did prevent the Russell body formation of μHC-ΔCH1 (compare Fig. 3B and 3C; see also Fig. 5B). In this regard, the cognate LC expression indeed exhibited the proposed protective effects against Russell body formation. Because the λLC and μHC-ΔCH1 subunits do not covalently assemble, non-covalent inter-chain interactions between them exerted the effects to prevent aggregation into Russell body. This result endorsed Corcos’ conclusion while disagreed with that of Valetti. The interesting twist was our finding that the protective effect of LCs on μHC-ΔCH1 aggregation was easily canceled when the JC subunit was also present in the same assembly milieu in the ER (Fig. 9F, top). Because μHC-ΔCH1 and JC interacted covalently via a disulfide bond at Cys-575 (see Fig. 8 B C), once these two proteins covalently aggregated into Russell body, an attempt to prevent the Russell body formation by non-covalent forces provided by the LC is either no longer effective or too weak to exercise its effect. In this case, Corcos’ claim that the LC is protective against Russell body formation is no longer valid. Yet, Valetti’s claim that LC and μHC-ΔCH1 aggregate into Russell body regardless of JC expression was not supported either. Additionally, our work demonstrated that C575S mutation can easily cancel the Russell body formation in the 3-chain expression of [μHC-ΔCH1 (C575S) + λLC + JC] (see Fig. 9F, second row). It is therefore conclusive that the covalent aggregation between JC and the Cys-575 of μHC-ΔCH1 did override the protective benefits of λLC. Interestingly, the deletion of CH1 was required in this [μHC-ΔCH1 + λLC + JC] 3-chain Russell body formation event because a specific blockade of HC–LC inter-chain disulfide bond formation by a minimally invasive μHC (C137S) was not able to induce Russell body phenotype in the [μHC (C137S) + λLC + JC] 3-chain co-expression setting.

Whenever a secretion-competent polymeric IgM (both hexamer and pentamer) was synthesized, the assembling IgM cargo was detected in the punctate structures that distributed broadly in the cytoplasm of expressing cells (Fig. 1A, 3C, 5A, 9E, 10F, 10G, 11D, 11E, 12E, 12F, 13E, 13F). Only after the blockade of covalent polymerization by C414/575S did the punctate localizations disappear and the cargo distributed to the Golgi and ER (Fig. 5A). Interestingly, a poorly secreted cargo (e.g., [μHC-ΔCH1 + LC] pair co-expression) also showed such cytoplasmic puncta to a similar extent (Fig. 5B, first row). Similarly, μHC (C137S) and μHC (C337S) mutants induced such punctate distribution when co-expressed with LC. We propose that punctate localization is a sign that μHC or μHC-ΔCH1 is achieving a multimeric state in the ER (covalent or not) and is independent of cargo secretion competence. Then, what are these cytoplasmic punctate structures that appear when polymeric IgMs assemble in the ER? Although we did not probe their identity, from the characteristic morphology, distribution pattern, and functional relevance, the punctate structures are likely to be the ER exit sites—which are often marked by Sec23, SURF4, ERGIC-53, ERp44, etc. [18, 112].

In addition to Russell body, Protein droplet inclusion bodies were also induced during IgM expression when the JC subunit was co-expressed with μHC (C414/575S) or μHC-ΔCH1 (C414/575S) mutant subunits (Fig. 8, D and G). While μHC (C414/575S) mutant induced Russell body when it was expressed alone (Fig. 4B), the co-expression of JC subunit led to protein droplet formation (Fig. 8C). Similarly, although μHC-ΔCH1 (C414/575S) localized to ER and Golgi at steady state (Fig. 4B), JC co-expression abruptly changed the solubility of μHC-ΔCH1 (C414/575S) mutant and induced protein droplet inclusion bodies in the ER (Fig. 8G). Instead of just sitting there as an inert bystander, JCs clearly interacted with these mutant μHC subunits in the ER non-covalently and modulated their condensation propensity. Unlike the previously characterized scFv-Fc-stp [84], we did not readily find evidence suggesting that μHC (C414/575S) and μHC-ΔCH1 (C414/575S) have notable solubility problems at high concentrations or cryoglobulin-like characteristics at temperatures below 37°C. The mechanistic process of how JC facilitated the droplet inclusion body formation was elusive. Whether or not LLPS played roles in this droplet formation was also unknown.

While these protein droplets can be readily distinguishable from the Russell bodies by using DIC microscopy and immunofluorescence microscopy, there have long been confusions in the scientific and clinical literature in that both types of inclusion bodies are indiscriminately referred to as Russell bodies merely based on their shared spherical shapes. On the one hand, we agree that Russell body, protein droplet, and crystalline inclusion bodies are variations of a common theme related to the intra-ER inclusion bodies of immunoglobulins. On the other hand, a series of characterizations on various IgG mAbs have started unveiling that there are some rules governing the relationship between the types of induced inclusion bodies and the physicochemical characteristics of individual immunoglobulin clones that induce them [50–56, 84, 113]. Suppose we can comprehend the underlying rules for different inclusion body formations and correlate them with different condensation propensities of individual mAbs. In that case, such knowledge will become indispensable when assessing the manufacturing suitability of a therapeutic antibody because we can forecast the antibody’s intrinsic high condensation propensity by looking at the cell phenotypes during recombinant overexpression [51]. Likewise, the knowledge can be applied in disease diagnosis when assessing the risk of renal toxicity for multiple myeloma patients because the inclusion body detected in the patient’s plasma cells would predict the severity of intrinsic condensation propensity embedded in the overproduced monoclonal antibody [51].

Can IgMs also induce intracellular protein crystals in the ER? Are IgMs too complex or heterogeneous (in terms of disulfides, N-glycans, and hexamer/pentamer mixture) to induce intracellular crystallization events? At least, SAM-6 did not have such notable characteristics. However, in the clinical literature, there are numerous reports on intra-ER protein crystals composed of IgMs [114–117]. Examples of key determinants responsible for intra-ER crystallization have been elucidated for various IgG mAb clones using computational modeling and targeted mutagenesis [51, 52, 54, 56, 113]. We still hardly know what structural determinants or sequence-based motifs underscore such intra-ER crystallization events for IgMs. Given the potential values of crystalline immunoglobulins as a promising method to achieve high-concentration biologics formulation [118] and to facilitate alternative downstream processing [119], understanding the process of IgM crystallization would add tremendous value when developing IgM-based protein therapeutics cost-effectively.

## Abbreviations

CH1: heavy chain constant domain-1
CH2: heavy chain constant domain-2
DIC: differential interference contrast
ER: endoplasmic reticulum
Fab: antigen-binding fragment
Fc: crystallizable fragment
Fv: variable fragment
HEK: human embryonic kidney
HC: heavy chain
IF: immunofluorescent
Ig: immunoglobulin
JC: J-chain
mAb: monoclonal antibody
LC: light chain
LLPS: liquid–liquid phase separation
MALS: multi-angle light scattering
PBS: Phosphate-buffered saline
pIgR: polymeric immunoglobulin receptor
SC: secretory component
scFab: single-chain Fab
scFv: single-chain Fv
SDS-PAGE: sodium dodecyl sulfate polyacrylamide gel electrophoresis
SEC: size-exclusion chromatography
VH: heavy chain variable region
VL: light chain variable domain

## 5. Acknowledgments

The authors thank Alice Bakker for cross-site work coordination at the beginning of this project and Christy Tinberg for analyzing the publicly reported SAM-6 VL and VH primary sequences for potential abnormality. We thank our colleagues at SARC for an initial attempt to optimize the hexamer IgM purification method. HH is personally grateful to Yoko Azumi for her continuous encouragement.

## 6. CRediT authorship contribution statement

**Haruki Hasegawa:** Conceptualization, Methodology, Validation, Investigation, Data Curation, Writing - Original Draft, Writing - Review & Editing, Visualization, Supervision, Project administration. **Songyu Wang:** Resources, Investigation, Data Curation. **Eddie Kast:** Investigation, Data Curation. **Hui-Ting Chou:** Investigation, Data Curation. **Mehma Kaur:** Resources. **Tanakorn Janlaor:** Resources. **Mina Mostafavi:** Resources. **Yi-Ling Wang:** Resources. **Peng Li:** Resources.

---

**Supplement 1.**
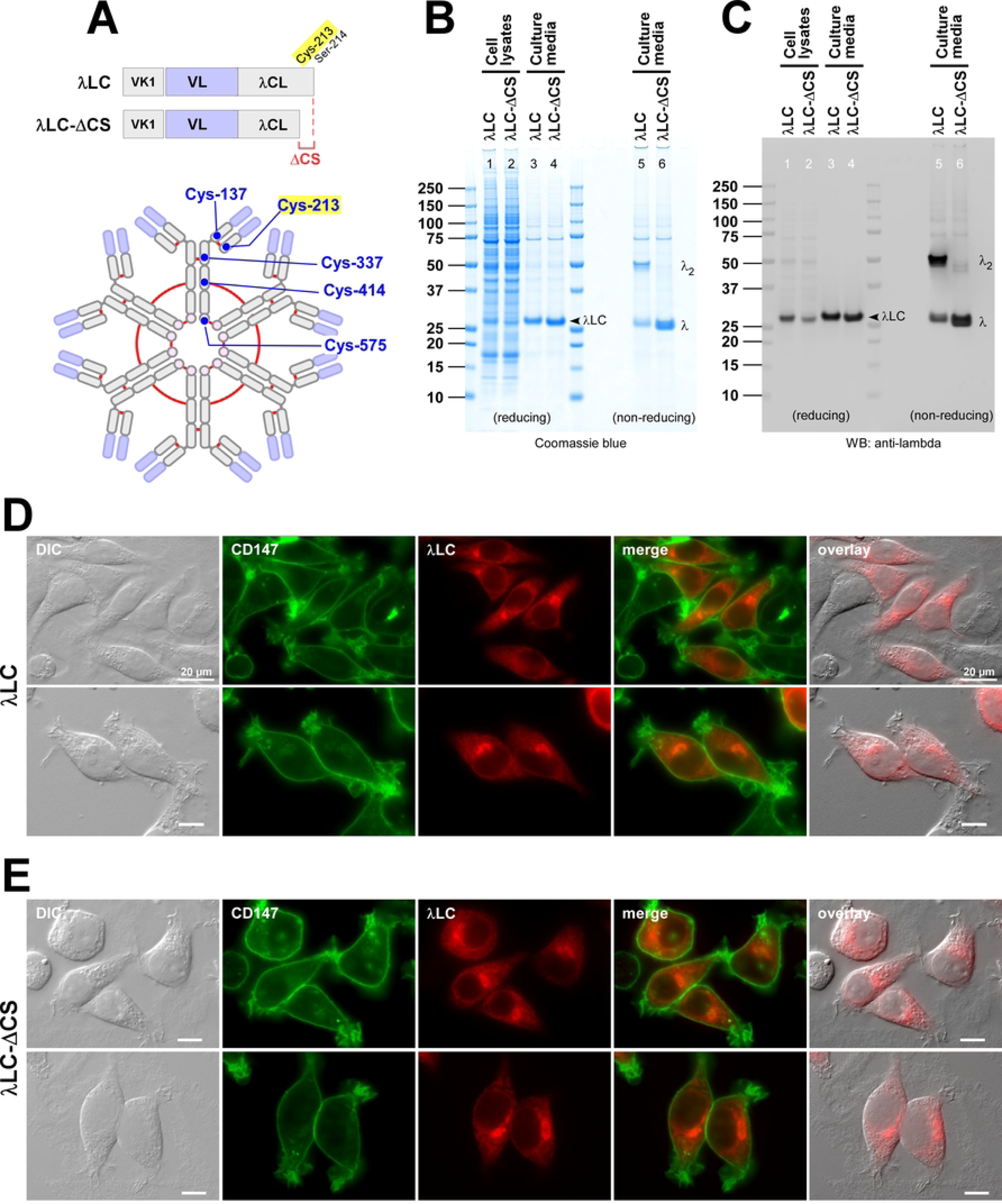
The penultimate Cys-213 of the λLC is required for inter-chain disulfide bond formation. (A, top) Schematic representation of the full-length SAM-6 λLC (top row) and its ΔCS mutant (second row) in which two C-terminal amino acids (Cys-213 and Ser-214) are deleted. (A, bottom) The position of Cys-213 residue involved in the HC‒LC inter-chain disulfide bond is highlighted in yellow in the context of the hexameric IgM diagram. Solid red lines represent the inter-chain disulfide bond connectivity. (B, C) HEK293 cells were transfected with full-length λLC (lanes 1, 3, 5) or its ΔCS mutant (lanes 2, 4, 6). At day-7 post-transfection, cell lysates (lanes 1 and 2) and cell culture media samples (lanes 3 and 4) were prepared and resolved by SDS-PAGE under reducing conditions followed by Coomassie blue staining (B) or by Western blotting (C). The day-7 cell culture media were also analyzed by Coomassie staining or Western blotting after resolving the proteins under non-reducing conditions (B and C, lanes 5 and 6). Blotted membranes were probed with polyclonal anti-λLC antibodies. The corresponding protein band for the λLC subunit is pointed by an arrowhead and labeled (lanes 1‒4). Monomeric and dimeric free λLC subunit is labeled next to lane 6. (D, E) Fluorescent micrographs of HEK293 cells transfected with full-length λLC (D) or ΔCS mutant (E). On day-3 post-transfection, cells were fixed, permeabilized, and co-stained with FITC-labeled anti-CD147 antibody and Texas Red-labeled anti-human λLC antibody. Green and red image fields were superimposed to create ‘merge’ views. DIC and red image fields were superimposed to create ‘overlay’ views.

**Supplement 2.**
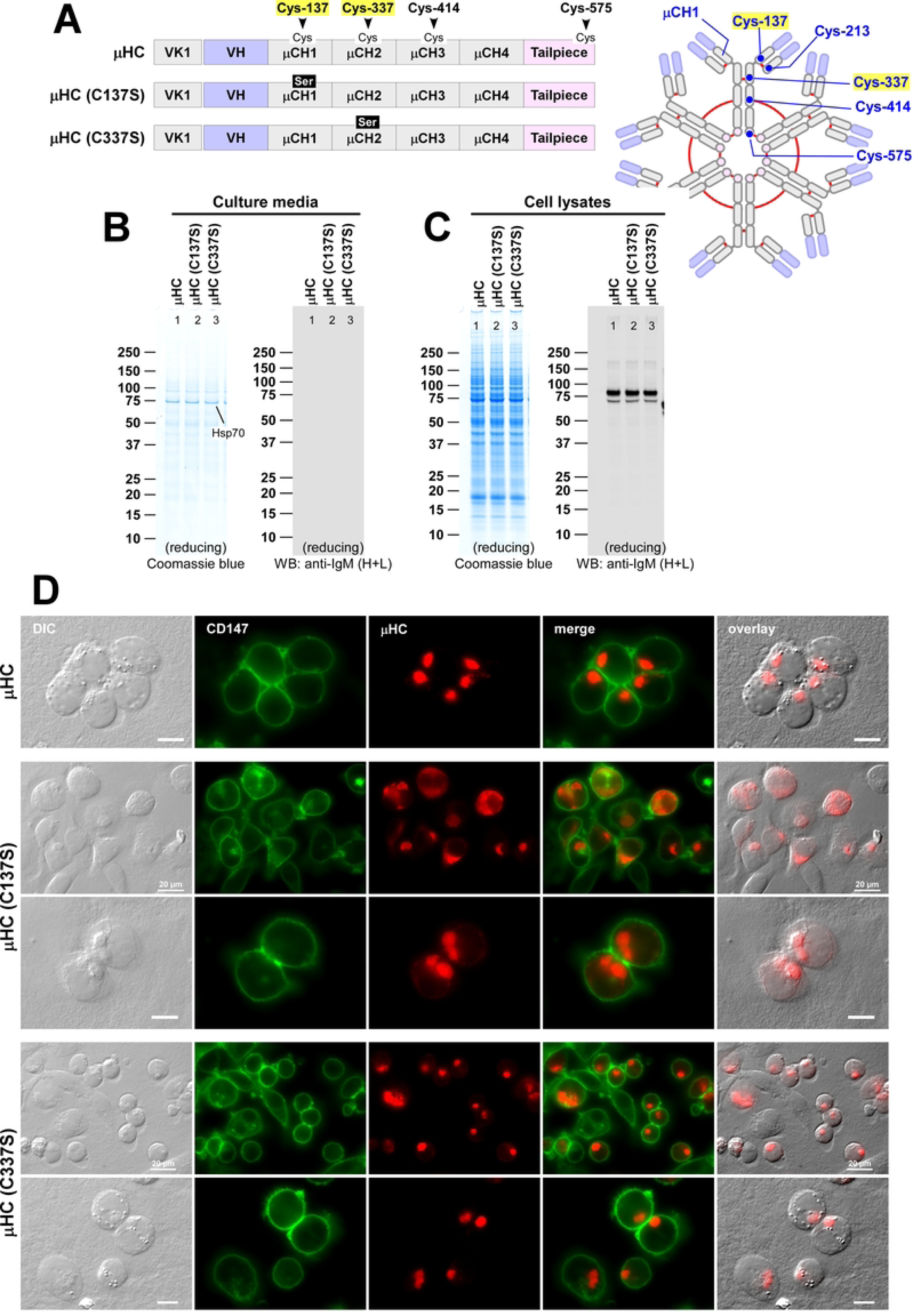
Effect of C137S and C337S point mutations on the expression, secretion, and subcellular localization of μHC subunit. (A, left) Schematic representation of parental SAM-6 μHC (top row) and its C137S and C337S mutants (second and third rows). (A, right) The position of Cys-137 and Cys-337 residues is highlighted in yellow in the context of the hexameric IgM diagram. Solid red lines represent the inter-chain disulfide bond connectivity. (B, C) HEK293 cells were transfected with parental μHC and its mutants, as shown at the top of each lane. At day-7 post-transfection, cell culture media (B) and cell lysates (C) were prepared and resolved by SDS-PAGE under reducing conditions followed by Coomassie blue staining (B, C, left panels) or by Western blotting (B, C, right panels). Blotted membranes in B and C were probed with polyclonal anti-IgM (H+L) antibodies. Both parental and mutant μHCs failed to secrete to the culture media. (D) Fluorescent micrographs of HEK293 cells transfected with parental μHC (top row), μHC (C137S) mutant (second and third rows), or μHC (C337S) mutant (fourth and fifth rows). On day-3 post-transfection, cells were fixed, permeabilized, and co-stained with FITC-labeled anti-CD147 antibody and Texas Red-labeled anti-human μHC antibody. Green and red image fields were superimposed to create ‘merge’ views. DIC and red image fields were superimposed to create ‘overlay’ views.

**Supplement 3.**
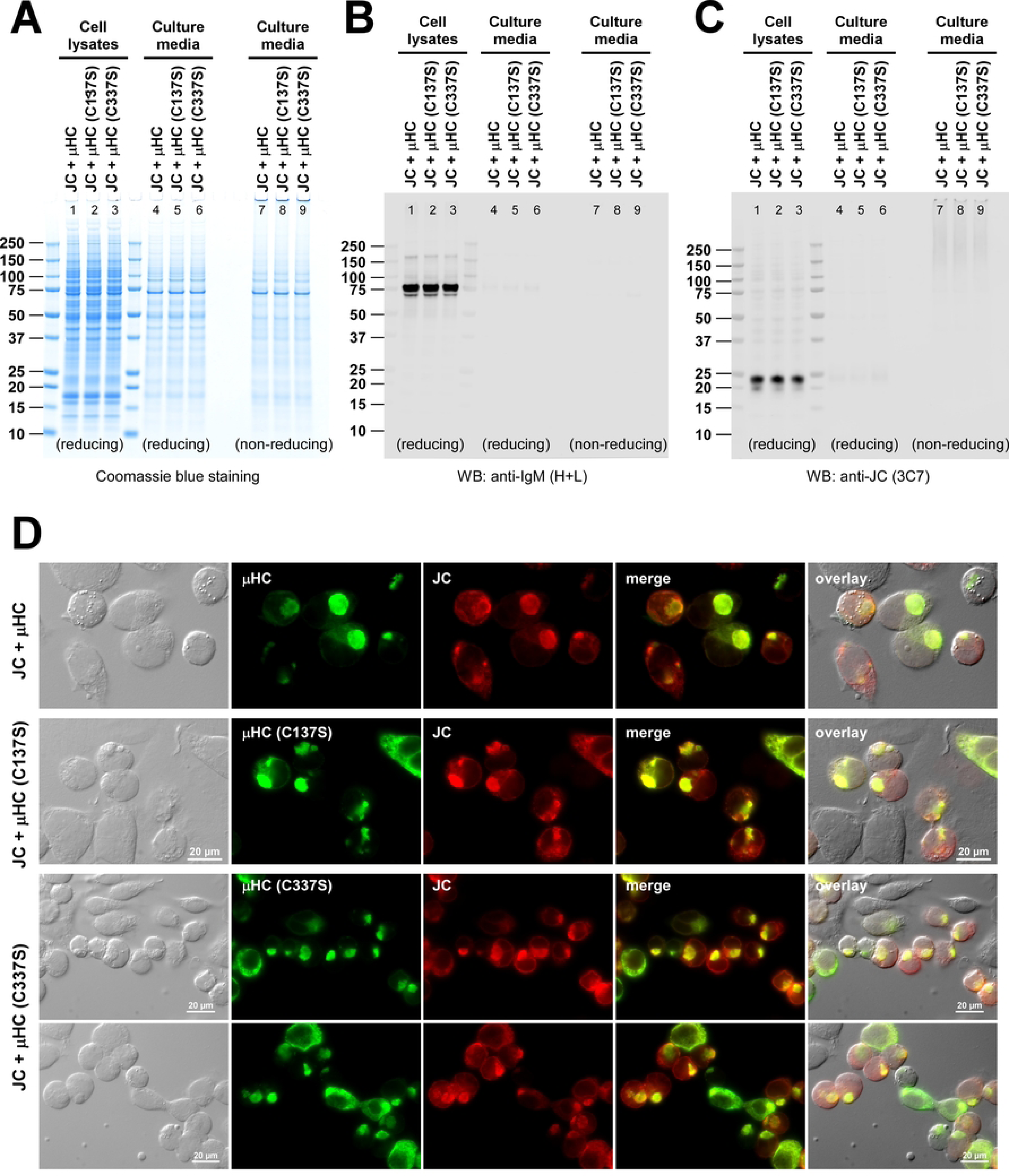
Co-aggregation of J-chain with C137S and C337S mutant μHC subunits. (A‒C) HEK293 cells were co-transfected with JC and one of the following μHC constructs: parental μHC (lanes 1, 4, 7), μHC (C137S) (lanes 2, 5, 8) and μHC (C337S) (lanes 3, 6, 9). At day-7 post-transfection, cell lysate samples were prepared (lanes 1‒3), and cell culture media were harvested (lanes 4‒9) to run SDS-PAGE under reducing conditions (lanes 1‒6) or non-reducing conditions (lanes 7‒9) followed by Coomassie blue staining (panel A) and Western blotting (panels B and C). Blotted membranes were probed with (B) polyclonal anti-IgM (H+L) antibodies or (C) monoclonal anti-JC antibody. A co-transfected construct pair is shown at the top of each lane. (D) Fluorescent micrographs of HEK293 cells co-transfected with JC and parental μHC (top row), JC and μHC (C137S) (second row), and JC and μHC (C337S) (third and fourth rows). A co-transfected construct pair is also shown on the left of each row. On day-3 post-transfection, cells were fixed, permeabilized, and co-stained with FITC-labeled anti-human μHC antibody (green) and monoclonal anti-human JC (red). Green and red image fields were superimposed to create ‘merge’ views. DIC and ‘merge’ were superimposed to create ‘overlay’ views.

**Supplement 4.**
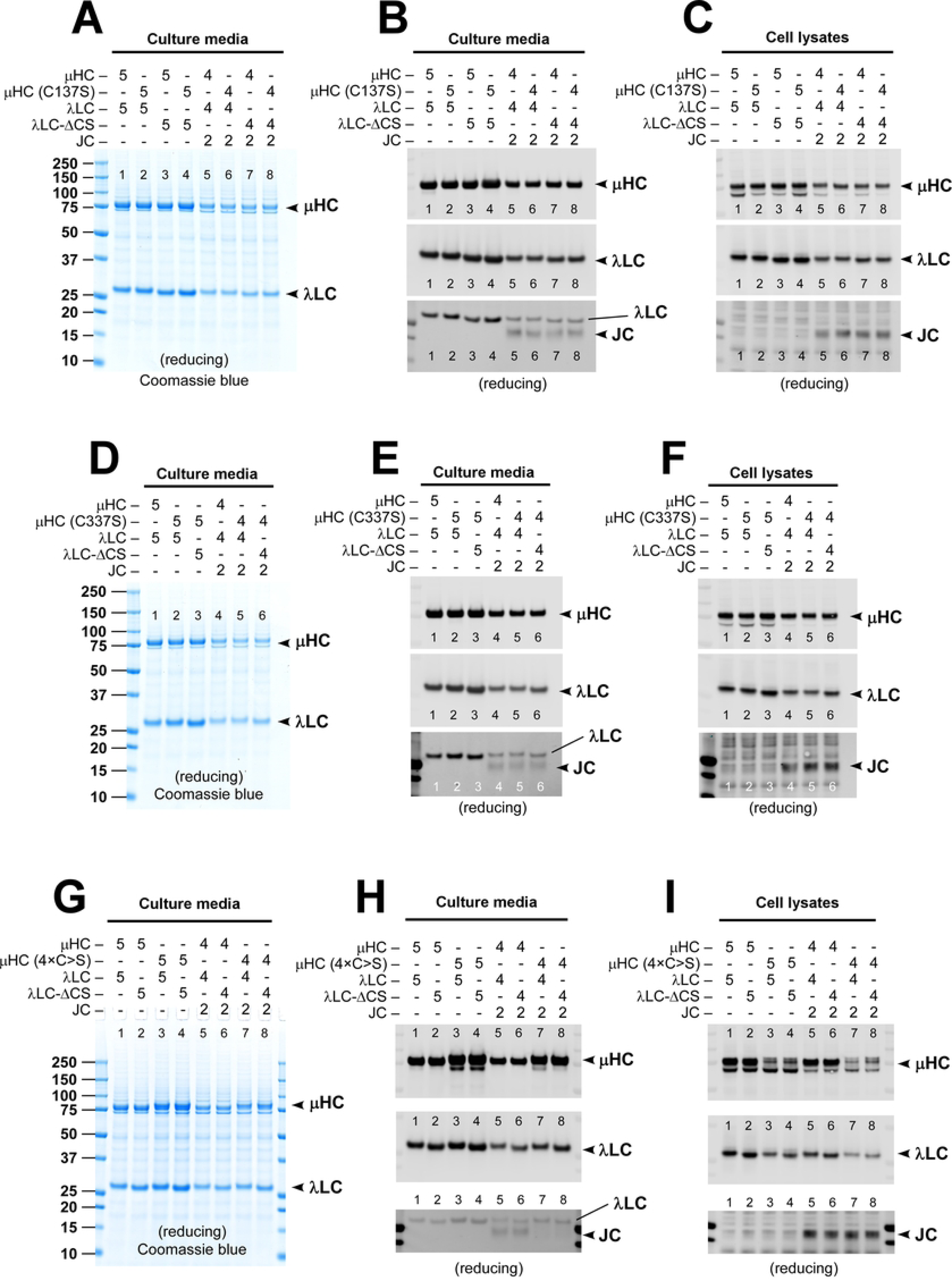
Effects of μHC (C137S), μHC (C337S), and μHC (4×C>S) mutation on IgM protein expression and secretion. (A‒C) The effect of μHC (C137S) mutant subunit on polymeric IgM expression and secretion was assessed in a 2-chain co-expression scheme (lanes 1‒4) and a 3-chain co-expression setting (lanes 5‒8). Parental subunit chains and mutants were co-transfected at the DNA ratio indicated at the top of each lane. At day-7 post-transfection, cell culture media (A, B) and cell lysates (C) were prepared and resolved by SDS-PAGE under reducing conditions followed by Coomassie blue staining (A) or Western blotting (B, C). Blotted membranes were probed with polyclonal anti-IgM (H+L) antibodies to detect μHC or μHC (C137S) (top panel) and λLC or λLC-ΔCS (second panel) or with monoclonal anti-JC antibody (third panel). (D‒F) The role of Cys-337 in polymeric IgM expression was tested by replacing the parental μHC with μHC (C337S) mutant subunit in a 2-chain co-expression scheme (lanes 1‒3) and a 3-chain co-expression setting (lanes 4‒6). Subunit chains were co-transfected at the DNA ratio indicated at the top of each lane. At day-7 post-transfection, cell culture media (D, E) and cell lysates (F) were prepared and resolved by SDS-PAGE under reducing conditions, followed by Coomassie blue staining and Western blotting. Blotted membranes were probed with polyclonal anti-IgM (H+L) antibodies to detect μHC or μHC (C337S) mutant (E, F; top panel) and λLC or λLC-ΔCS (E, F, second panel) or with monoclonal anti-JC antibody (E, F, third panel). (G‒I) The effect of μHC (4×C>S) mutant on polymeric IgM expression was tested in a 2-chain co-expression scheme (lanes 1‒4) and a 3-chain co-expression setting (lanes 5‒8). Subunit chains were co-transfected at the DNA ratio indicated at the top of each lane. At day-7 post-transfection, cell culture media (G, H) and cell lysates (I) were prepared and resolved by SDS-PAGE under reducing conditions, followed by Coomassie blue staining (G) and Western blotting (H, I). Blotted membranes were probed with polyclonal anti-IgM (H+L) antibodies to detect μHC or μHC (4×C>S) mutant (H, I; top panel) and λLC or λLC-ΔCS (H, I; second panel) or with monoclonal anti-JC antibody (H, I; third panel).

**Supplement 5.**
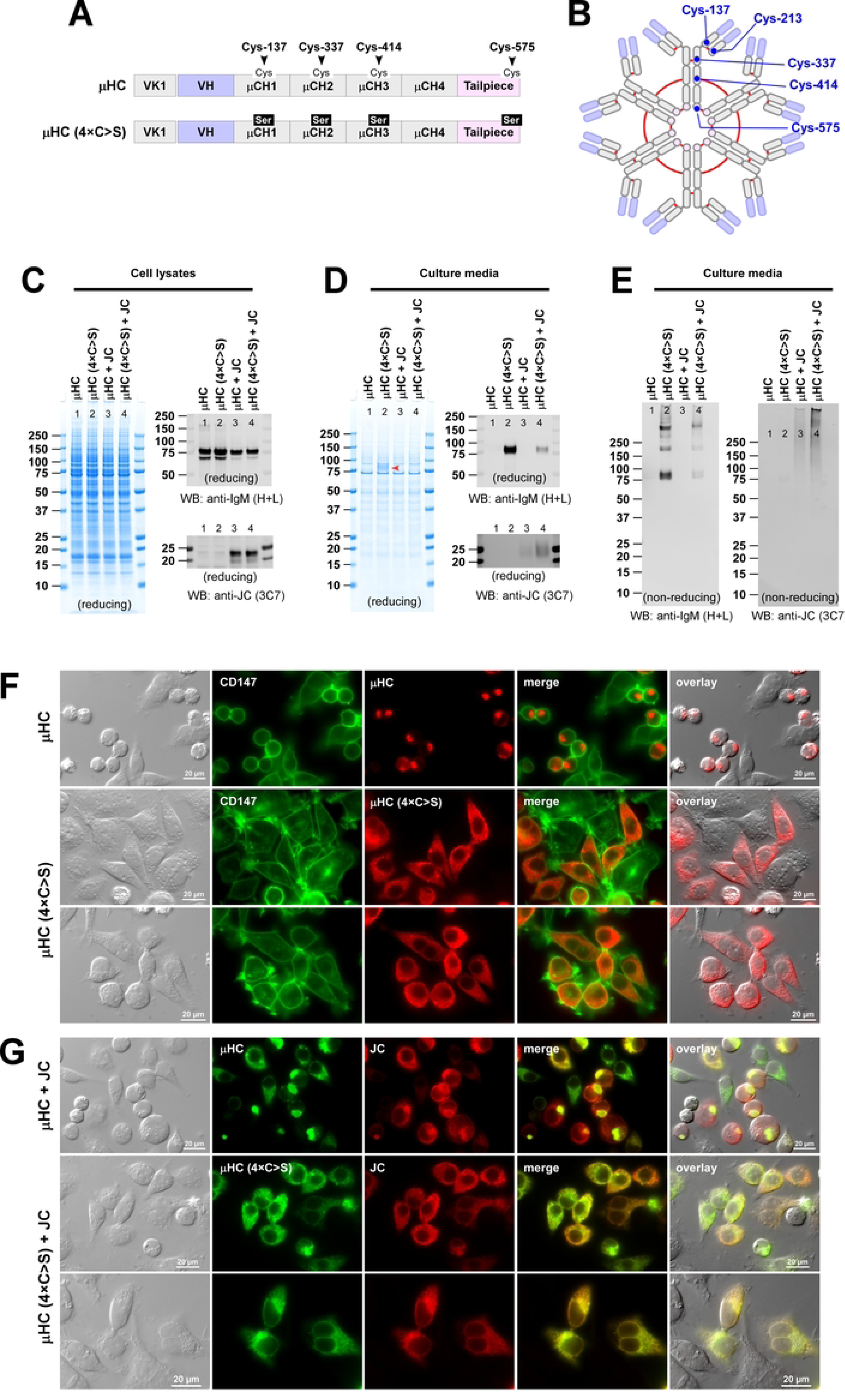
Effect of μHC (4×C>S) quadruple mutations on the expression, secretion, and subcellular distribution of μHC subunit. (A) Schematic representation of parental SAM-6 μHC (top row) and its 4×C>S mutant (second row). The positions of key cysteine residues important for the inter-chain disulfide formation are marked on the parental μHC. (B) The position of all four Cys residues involved in inter-chain disulfide bond formation on μHC and Cys-213 on λLC are depicted in the context of the hexameric IgM diagram. Solid red lines represent the inter-chain disulfide bond connectivity. (C, D) HEK293 cells were transfected with parental μHC alone (lane 1) or its 4×C>S mutant alone (lane 2). Likewise, the cells are co-transfected with μHC and JC (lane 3) or μHC (4×C>S) and JC (lane 4). At day-7 post-transfection, cell lysates (C) and culture media (D) were resolved by SDS-PAGE under reducing conditions followed by Coomassie blue staining (C, D; left panel) or by Western blotting (C, D; right panels). Blotted membranes in C and D were probed with polyclonal anti-IgM (H+L) antibody (top panel) or monoclonal anti-JC antibody (bottom panel). (E) Day-7 culture media were also analyzed by Western blotting after proteins were resolved under non-reducing conditions. Blotted membranes were probed with polyclonal anti-IgM (H+L) antibody (left panel) or monoclonal anti-JC antibody (right panel). (F) Fluorescent micrographs of HEK293 cells transfected with parental μHC (top row) or μHC (4×C>S) mutant (second and third rows). On day-3 post-transfection, cells were fixed, permeabilized, and co-stained with FITC-labeled anti-CD147 antibody and Texas Red-labeled anti-human μHC antibody. Green and red image fields were superimposed to create ‘merge’ views. DIC and red image fields were superimposed to create ‘overlay’ views. (G) Fluorescent micrographs of HEK293 cells co-transfected with μHC and JC (top row) or μHC (4×C>S) mutant and JC (second and third rows). Cells were co-stained FITC-labeled anti-human μHC (shown in green) and monoclonal anti-JC antibody followed by AlexaFluor594-conjugated secondary antibody (shown in red). Green and red image fields were superimposed to create ‘merge’ views. DIC and ‘merge’ were superimposed to create ‘overlay’ views.

**Supplement 6.**
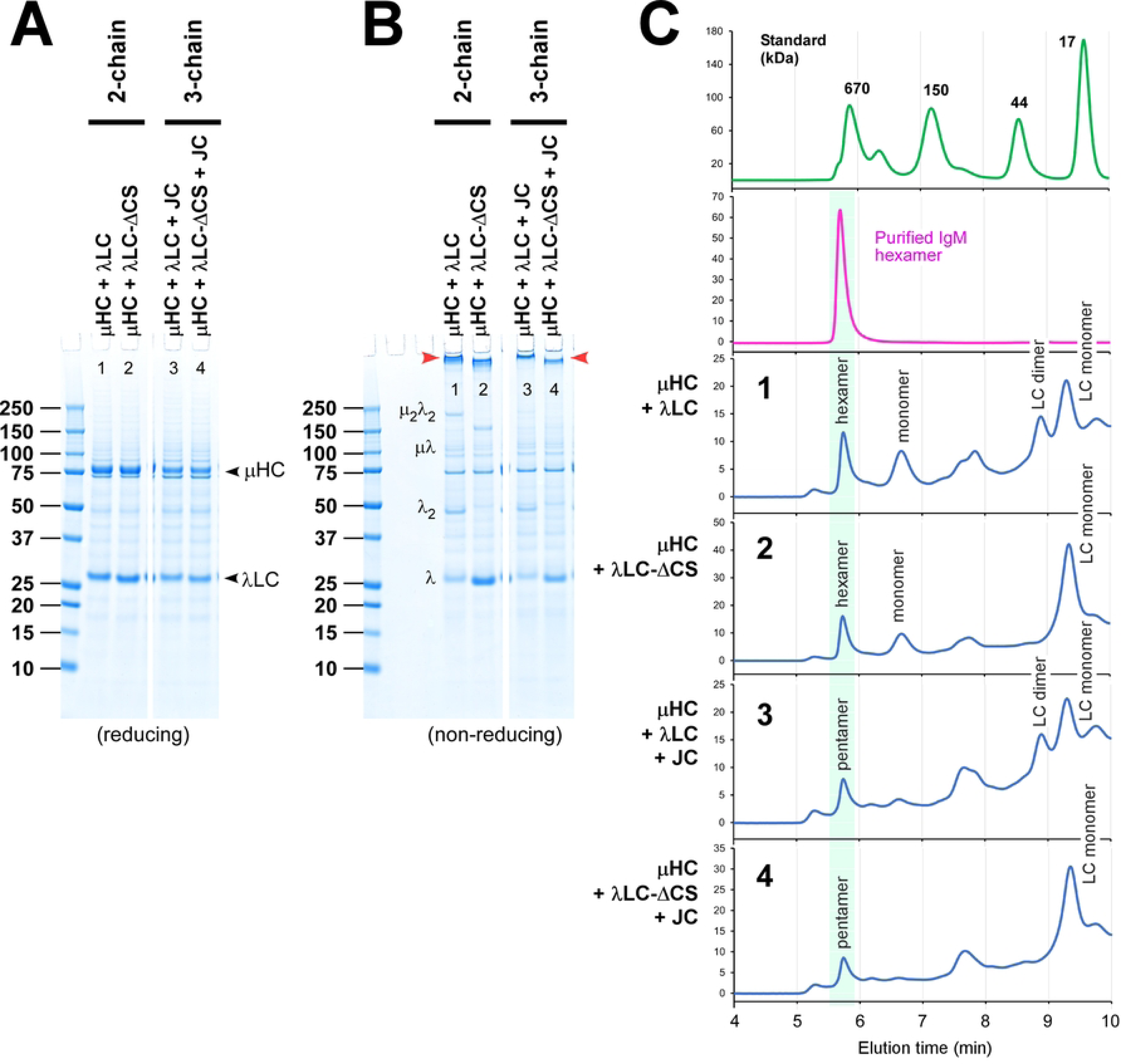
Assembly and maintenance of pentameric IgM-like products partially through non-covalent forces. (A, B) Secretion level and product quality of pentameric IgMs, IgM-like molecules, and assembly intermediates were compared in SDS-PAGE by resolving the day-7 cell culture media under (A) reducing and (B) non-reducing conditions. In the 3-chain expression setting to produce pentameric IgMs, parental μHC and JC were co-transfected either with the parental λLC (lane 3) or λLC-ΔCS (lane 4). In panels A and B, protein samples were analyzed in one single gel, but the gel image was digitally cropped to remove intervening lanes not directly related to this analysis. Covalently associated polymeric IgM and IgM-like products preserved in SDS-PAGE are marked by red arrowhead in panel B. (C) The quality of secreted products was examined by analytical SEC under non-denaturing, physiological pH assay conditions. SEC elution profiles for culture media shown in B, lanes 1‒4, are displayed in the chromatograms with corresponding numbers in C, panels 1‒4. The transfected subunit chain combination is also shown on the left side of each chromatogram. The elution peak corresponding to the designated polymeric IgM is shaded in light green in individual chromatograms.

**Supplement 7.**
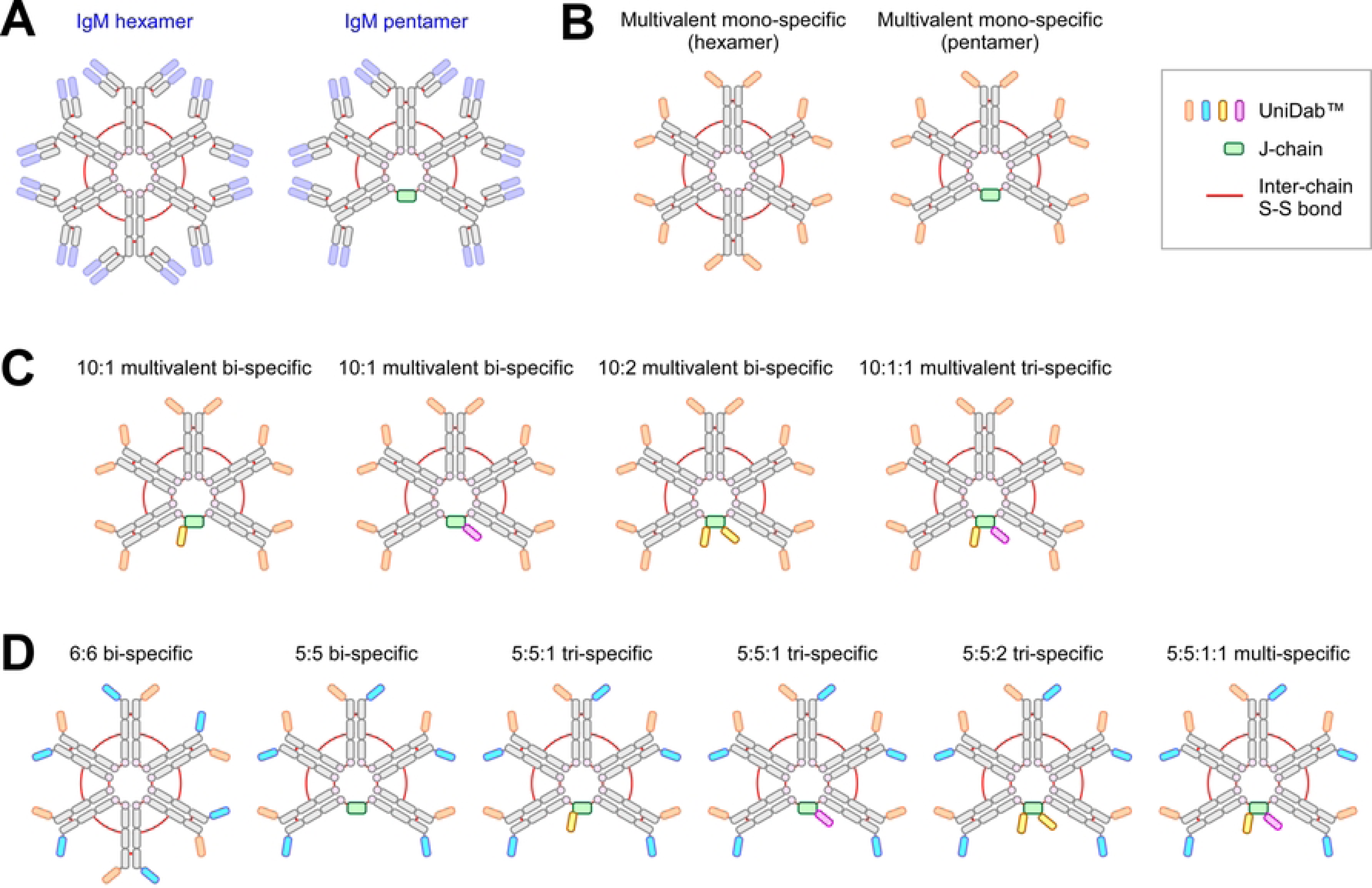
Diagrams of hypothetical IgM-like product design using the VH-domains derived from HC-only antibodies. (A) Conventional hexameric and pentameric IgM. (B) The Fab region of the conventional IgM was replaced by a UniDab™ module to illustrate the potential molecular designs for multivalent monospecific hexamer and pentamer. (C) Additional UniDab™ was fused to the N-terminus, C-terminus, or both termini of the JC subunit to produce multivalent bi-specific or tri-specific IgM pentamers. (D) Hypothetical molecule variations when UniDab™ based multi-specific design was combined with a hetero-Fc technology that can generate asymmetric monomer units that comprise the multivalent IgM-like molecules.

